# Who Grows There? A Course-based Undergraduate Research Experience to explore the human microbiome through 16S DNA metabarcoding

**DOI:** 10.1101/2024.07.22.600610

**Authors:** Graham S Sellers, Merideth Freiheit, Michael R Winter, Domino A Joyce, Darron A Cullen, David H Lunt, Katharine E Hubbard

## Abstract

We describe a two week Course-based Undergraduate Research Experience (CURE) to introduce students to next-generation DNA sequencing, molecular biology methods and a bioinformatic workflow. The CURE is designed to take students with little to no technical and bioinformatic experience through key steps of the protocol through scaffolded laboratory and computational practicals. Our students extract and amplify human microbiome DNA using 16S ribosomal RNA specific primers, then construct a sequencing library for Oxford Nanopore based sequencing. They taxonomically assign the sequencing reads, and determine the ecological community composition using relevant software packages. Our students were able to successfully prepare sequencing libraries and analyse the data to produce relevant figures, demonstrating they met the learning objectives of the CURE. Students identified that they had developed higher level learning as defined by Bloom’s taxonomy, and that their confidence in practical work significantly increased as a result of doing the CURE. We share recommendations for implementation of the CURE in undergraduate curricula, and adaptations of the methods for use in schools outreach. Our CURE successfully provides training for students in genetic analysis in an enjoyable and relatively time and cost efficient manner, preparing them for future research or careers in modern molecular biology techniques.

## Introduction

Next-generation DNA sequencing technologies have revolutionised our understanding of biosciences, with applications from epidemiology to ecological monitoring (1–4). The advent of these technologies has meant that it is increasingly quick, easy and inexpensive to generate large amounts of DNA sequencing data for bioinformatic analysis. There is therefore a need to train undergraduate biologists in these technologies. However, the methods involved require high levels of understanding, technical skill and accuracy. Sequencing methods generate large datasets, so also require training in bioinformatic workflows and ‘big data’ analysis methods. This paper describes a Course-based Undergraduate Research Experience (CURE) to introduce undergraduates to wet-lab and computational skills required to answer research questions using sequencing methods.

It has long been recognised that inquiry-driven and/or research-led approaches to teaching scientific practicals have many advantages over traditional ‘cook-book’ style practicals (5). An increasingly common model is the CURE, whereby students are exposed to authentic research-driven projects that emphasise the scientific process, discovery, real world relevance and collaboration as much as technical skill (6, 7). These take place within credit bearing modules, typically with one instructor supervising multiple students, so are more scalable and inclusive than requiring students to find individual research internships (7–9). Instructors in a variety of bioscience contexts have designed CUREs to embed research-led experimental work (10–13). These CUREs vary in their scientific content, but also in the degree of student independence afforded. Brownell and Kloser identify four ‘levels’ of CURE, ranging from ‘structured’ CUREs where students have limited choice in the methods used but have independence over analysis, through to ‘authentic’ CUREs where students have full autonomy throughout the research process (6). CUREs have been associated with increased student ownership of their work (14), scientific thinking (15), data analysis skills (15), intention to continue in research (16) and even graduation rate (17). It has been suggested that these positive impacts are felt most by students from historically or currently marginalised groups (8). As such, there are numerous benefits to the CURE model as authentic research-led training within bioscience.

Our CURE is designed as an introduction to DNA metabarcoding, a high throughput sequencing technique to taxonomically identify organisms from the sequence of a specific marker gene (18, 19). It is a relatively cost-efficient, accurate, sensitive and rapid method of species determination, and less prone to bias than traditional ecological sampling (20). It has been used successfully in contexts including freshwater ecology (21), ecological network analysis (22), forensic science (4)(23) and soil microbiology (23). In a typical workflow DNA is extracted, PCR amplified using adapted 16S rRNA primers, a sequencing library prepared and sequenced using Oxford Nanopore technology (24). Bioinformatic analysis taxonomically defines each sequence, with the community composition determined from read frequency (25). Bacterial classification typically uses Operational Taxonomic Units (OTUs) defined at species, genus or other level as appropriate (25). Our CURE uses these techniques to explore an individual human microbiome. The microbiome is highly personalised and dynamic, reflecting the sample site, lifestyle and diet of the individual (26), and also with various health conditions (26, 27). Microbiome studies therefore lend themselves to CUREs as there is no ‘right’ or ‘known’ result, enabling students to undertake genuine open-ended inquiry based research in a relatively short space of time and cost effective way.

### Intended Audience

This CURE is designed for use with second year undergraduates on bioscience programmes at a medium sized UK university; we present data for two cohorts of students. The CURE forms half of a second year ‘Genetic Analysis’ module, with the other half covering theoretical content. Students taking our CURE have previously studied one introductory genetics module, which does not include inquiry-driven practicals. We introduced this CURE to (i) give students training in fundamental and modern genetic analysis techniques and (ii) to provide their first exposure to research-led experimental work. The module is compulsory for students on BSc Biology and BSc Biochemistry, and optional for BSc Zoology students. The CURE is delivered by a lead academic and postgraduate demonstrators supported by technicians. Students complete the work in groups of two or three to promote peer learning and build confidence.

### Learning Time

Our CURE takes place over an intensive two week period, preceded by online activities to develop understanding of core principles, followed by a write-up period (Figure 1). Wet-lab practicals build student confidence and skill in fundamental techniques before they prepare sequencing libraries. Sequencing takes place over the weekend, providing data for computational practicals in the second week; these two weeks could be separated if required.

**Figure 1:**
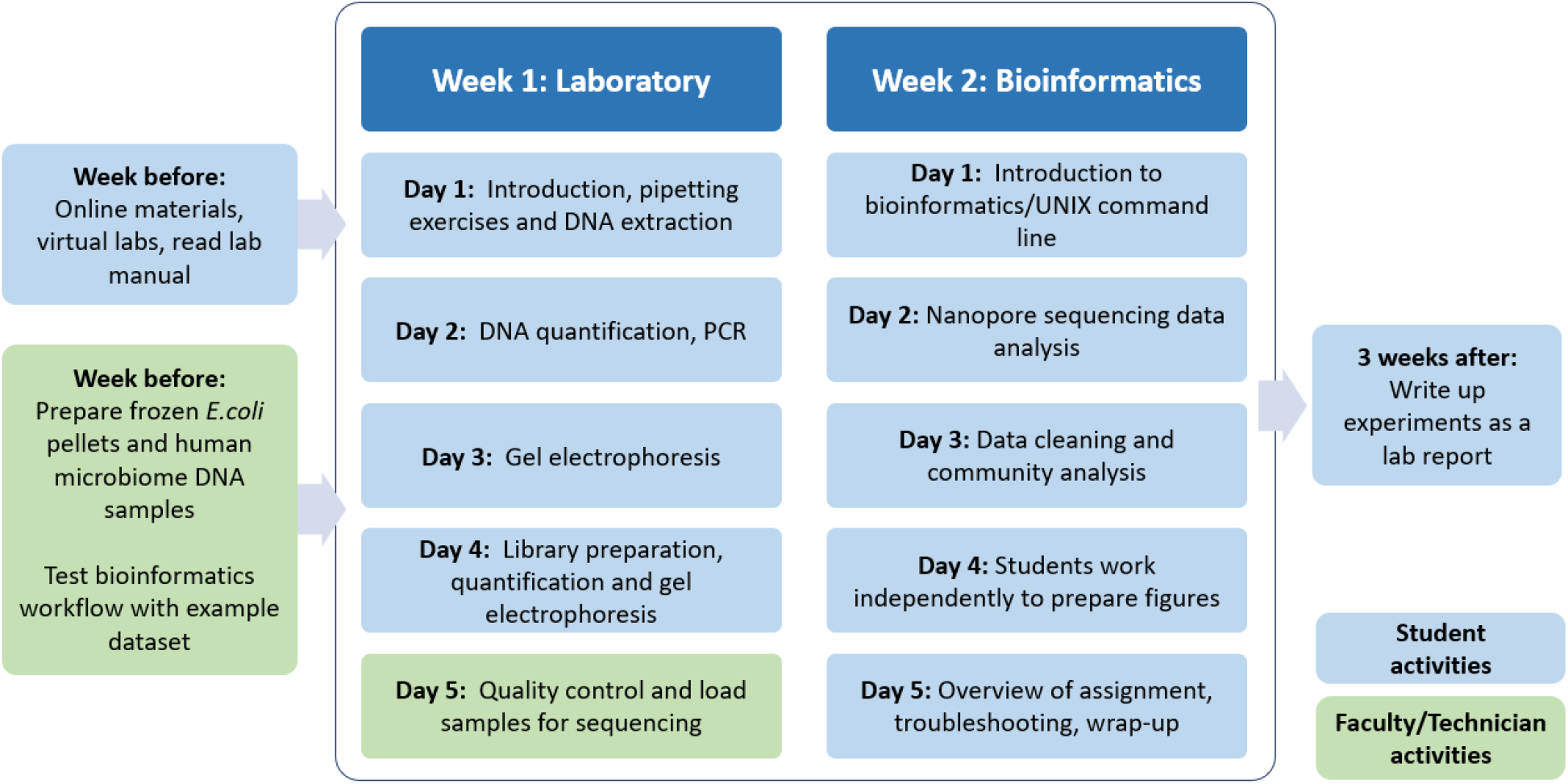
Overview of CURE design.

### Prerequisite student knowledge

Students require some knowledge of fundamental genetics and genetic methods (e.g. PCR, gel electrophoresis). In our context this is covered in the first half of the Genetic Analysis module and in the pre-lab content. It would be possible to teach these topics as an intensive short course at the start of the CURE. Our students have some prior experience of using Rstudio for basic data analysis (e.g. t-tests, box-and-whisker plots), but this could be taught within the context of the CURE given appropriate time and support. To help our students prepare for the practicals they may complete interactive simulations provided by Learning Science Ltd before coming to the laboratory. Students are also provided with two research papers to read in advance, a review paper on the human skin microbiome (28), and an experimental paper describing DNA extraction (29).

### Learning Objectives

By the end of the CURE, students should be able to demonstrate the following competencies:

- Accurately use a micropipette.
- Extract and quantify purified DNA from a bacterial culture.
- Run and analyse a PCR using bacterial 16S markers via gel electrophoresis.
- Use the above skills to prepare and quantify a genetic library for Flongle sequencing.
- Learn the basic utilities and functions of the Unix command line.
- Experience bioinformatic analysis of Flongle sequencing data including sequencing data quality control and taxonomic assignment in Python.
- Analyse bioinformatic outputs and produce descriptive figures in R.
- Write a scientific report that focuses on a specific research question within the context of the experiments performed.

### Procedures

The CURE is designed as an introduction to modern genomics methods and bioinformatics, for a cohort of students with limited experience. We therefore designed a workflow where students learn key techniques, and are exposed to more advanced technical aspects, but do not complete the entire procedure themselves. While CUREs should promote engagement with scientific thinking, there is a need for balance between open-endedness and scaffolded learning so that students can develop autonomy without being overwhelmed by a lack of structure (30).

To build student confidence students use ‘practice’ samples of *E*.*coli* to extract DNA, perform PCR and gel electrophoresis, before using the same techniques to prepare and quantify their microbiome sequencing libraries. Students prepare libraries from pre-prepared human microbiome DNA samples and check the quality and quantity of their libraries, but staff load the sequencer to ensure correct procedure (Figure 1). Similarly, the bioinformatic analysis is designed so that students perform some steps themselves (e.g. data import, construction of figures), but more complex aspects of the workflow (e.g. taxonomic assignment) use a pre-written data analysis pipeline. Where possible, we use methods that are relatively inexpensive and reliable that can be used at scale by students with limited molecular biology experience. For example, we use Modular universal DNA (Mu-DNA) extraction as a low-cost, high-throughput method (29). We also use Oxford Nanopore’s Flongle flow cells as a cost effective alternative to the MinION for smaller sequencing libraries (31).

## Materials

To minimise biological risk we prepared biological samples for students in advance. For the initial DNA extraction samples we prepared overnight cultures of *E. coli* K12 in liquid LB media, which were then aliquoted and flash-frozen to give dead pellets for extraction. For the microbiome samples, 12 swabs were taken from the human subject (3x left arm, 3x right arm, 3x left leg, 3x right leg) and DNA extracted using the Mu-DNA method (29) before giving to students for library preparation.

For the bioinformatics workflow we prepared Jupyter notebooks for the students comprising a mixture of explanation, instruction, and their bioinformatic analysis activities. These notebooks were hosted in a JupyterLab environment on a University of Hull Linux server. This JupyterLab application is designed to manage multiple user sessions in interactive computing (32, 33) and is well suited to class teaching with individual logins. The computational environment for bioinformatics required use of a bash shell, R, python, and several bioinformatics analysis programmes. These are listed as part of the resources, and installed using the Conda package manager (34). The documentation, data, environment description, and workflow was managed and distributed to the students via use of git version control on GitHub (35). These are available under a permissive CC-BY licence at https://github.com/davelunt/genetic-analysis-bioinformatics-practical

### Laboratory procedures (Week 1)

**Day 1**. Students are first introduced to the practical course via a short presentation, including health and safety information. Students then complete a series of pipetting exercises to re-familiarise themselves with a micropipette, reinforcing both volume control and spatial accuracy. Students then each extract DNA from a frozen bacterial pellet, which is then stored at 4°C overnight.

**Day 2**. DNA samples are quantified on a Qubit 3.0 fluorometer using the High-sensitivity (HS) dsDNA assay (Invitrogen). Technical staff prepare a Qubit mastermix containing standardised concentrations of the required dyes in advance; students mix the mastermix and their DNA sample and quantify on the fluorometer. Samples with a minimum of 1 ng/μl or above indicate a successful DNA extraction. Students then prepare 16S PCRs from their samples and a positive and negative control, which amplify overnight.

**Day 3**. PCR products are visualised via gel electrophoresis. Students mix PCR products with loading dye and load samples onto sodium borate gels containing GelRed nucleic acid stain (Biotium), using EasyLadder I DNA ladder (BioLine) to identify fragment lengths. Students run and visualise gels, with successful PCRs giving a clearly visible band (Figure 2A).

**Figure 2:**
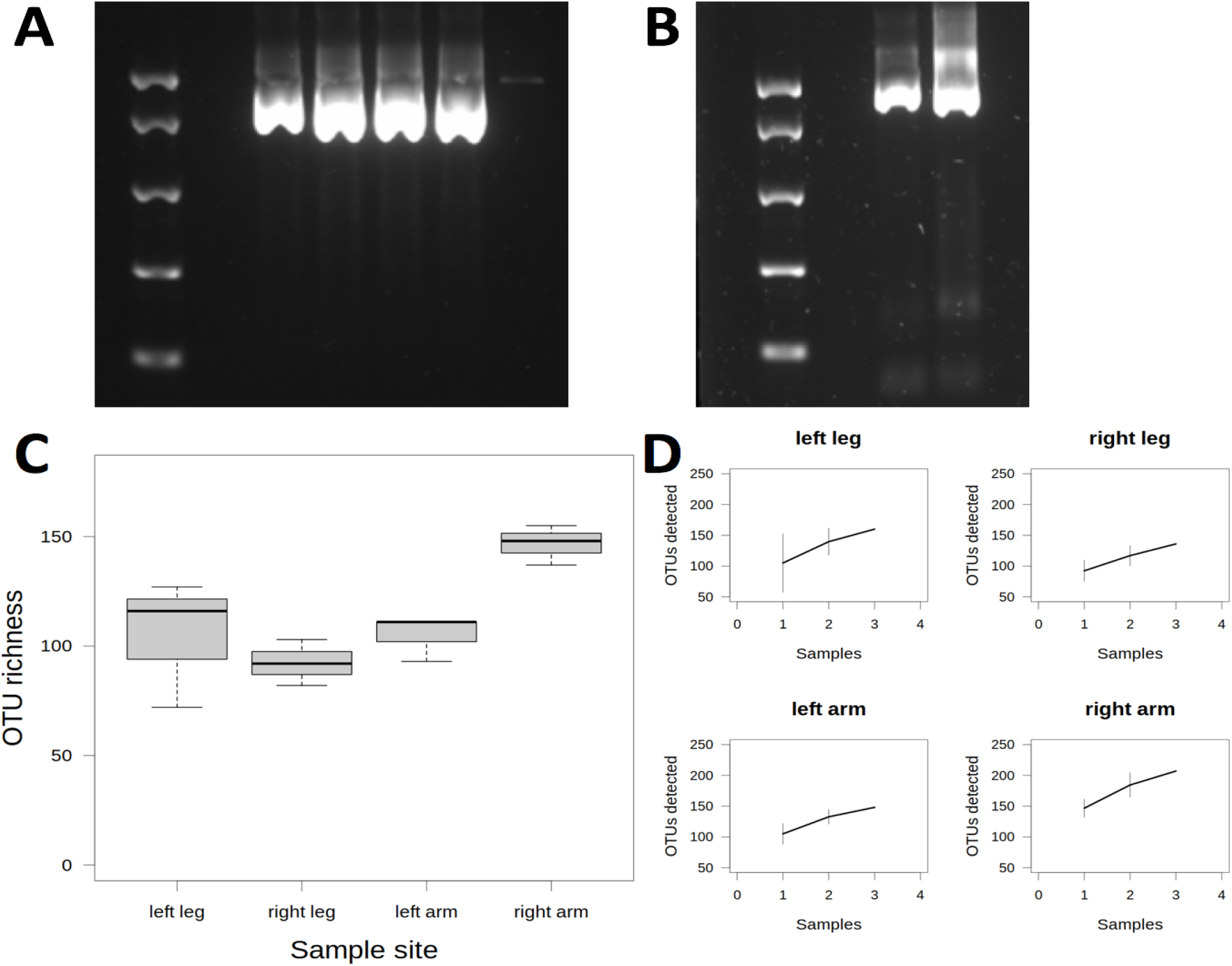
Examples of student data analysis. A: Gel electrophoresis image of student 16S PCR products. Lane 1 contains EasyLadder I with bands at 2000, 1000, 500, 250 and 100 bp; Lanes 3-5 contain student produced 16S samples, lanes 6 and 7 are PCR positive and negative controls respectively. Student extracted samples have a clear band at ∼1500 bp. B: Gel electrophoresis image of student sequencing library preparation. Lane 1 contains EasyLadder I, lanes 3-4 library PCR products at ∼1700 bp. C: Boxplot of OTU (Operational Taxonomic Unit) richness at the four sampling sites and D: OTU accumulation plots, showing that OTU saturation had not been reached within the three replicate samples at each site.

**Day 4**. Having learned the essential techniques with the practice samples on Days 1-3, students then prepare their genetic sequencing libraries. Pre-extracted DNA samples from the human microbiome samples are provided to students. Each group of 2-3 students prepares 16S sequencing libraries for 2 out of the 12 microbiome samples. Libraries are prepared with the Oxford Nanopore Technologies 16S Barcoding Kit (SQK-RAB204), which uses a PCR based method to add sequencing primers to the end of each DNA sample. We add a unique primer to each sample to allow for multiplexing of samples on the nanopore (i.e. all 12 samples added simultaneously). After library preparation, samples are quantified with the Day 2 Qubit protocol, and visualised by gel electrophoresis using the Day 3 protocol (Figure 2B). Students may calculate molar concentration of each sample, as samples need to be of equal molarity for multiplexing.

**Day 5**. Students are not required to come to the lab. Technical staff check DNA library concentrations, and mix samples together for multiplexing. Samples are loaded onto the Flongle and sequencing occurs over the weekend.

Full details are available in Supplemental Information 1: Lab Manuals

### Bioinformatic analysis (Week 2)

Our students had no previous experience of bioinformatics prior to the CURE, so activities were structured to introduce key concepts from using the command line to generating microbial community analyses. The bioinformatic analysis was conducted in person via interactive Jupyter virtual workbooks (33), but could be adapted for remote participation if required. The analysis pipeline was partly conducted on our university high performance computing cluster, required for the handling of large datasets. Example smaller datasets were constructed for whole class use (e.g. by removing OTUs that only occurred at low frequencies) to reduce the amount of processing power required while still allowing students to develop relevant computational skills. Teaching strategies included live demonstrations of the key steps by the instructors, notes written for each step, and practical activities with example code and step by step instruction. Students checked their work regularly with demonstrators, and any students who were struggling were given 1:1 support.

**Day 1**. Students were supported in logging in to the JupyterLab server, git cloning the repository of data and workbooks, and familiarising themselves with Jupyter notebooks (NB the time required for these initial steps can be larger than expected). Students carry out self-paced demonstrator-supported activities to learn the Unix command line for bioinformatics. We have built in stop points, where the class can be brought back together, and where discussions are used to reinforce the learning. Instructors encourage self-directed learning by helping students to use essential tools, and then presenting them with an opportunity to use the same skills independently for a biological problem.

**Day 2**. Working in Jupyter notebooks with a mixture of written instruction and space for analysis, students begin data analysis. Students first explore the fastq file format to examine their data and how it is stored. They next perform quality control steps, cleaning the data with fastp (36) and SeqKit (37). They are then guided through taxonomic assignment of sequences using Kraken2 (25) and use a pre-written script to reformat the taxonomic output as .csv so that the data is easy to explore using R.

**Day 3**. Students conduct ecological community analysis in R, first using example datasets and then their own data. They complete exercises to refamiliarise themselves with coding in R, and then examine the taxonomic data from Day 2. They then extend these results by examining subsets such as proportional read presence and absence and dividing by sample site. Using these analyses, the students perform ecological analysis of the microbial community, determining OTU richness in each sample site. The final steps of ecological analysis use Vegan (38), an R package for community analysis.

**Day 4**. Students have a day of independent study to work on their analysis and identify any issues they want to discuss with staff on the final day.

**Day 5**. Students first review the main skills from Days 1-3, including R basics, data cleaning, and ecological analysis. Demonstrators discuss publication quality figure creation for the assignment. Students then independently elaborate on example code to create figures in R (Figure 2C,D). Students are brought together for a final in-person session for troubleshooting and questions, and review of the assignment requirements. A seminar is held to review key theoretical aspects of microbiomes (e.g. commonly found species), and to give details of the human subject which might be relevant to the microbiome results.

Full details are available in Supplemental Information 2: Bioinformatic Manuals

### Safety and ethical considerations

The CURE uses standard molecular biology methods (PCR, gel electrophoresis). We minimise chemical hazards through use of GelRed nucleic acid stain rather than ethidium bromide, and by technicians pre-aliquoting small volumes of reagents for students to use. We minimise biological hazards by preparing frozen pellets of *E*.*coli* K12 bacteria for DNA extractions; if students are preparing live cultures then a biological risk assessment may be required. Risk assessments are provided in advance (Supplemental Information 3), and critical safety instructions added to the lab manual and presentation slides. Due to local ethical approvals, students were unable to sample their own microbiomes. We therefore sampled bacteria from one of our postgraduates who was covered by existing research ethical approval.

### Assessment methods for determining student learning

Our CURE is assessed through students submitting a formal experimental write-up of 2000 words, including Introduction, Methods, Results and Discussion, and at least one results figure. Students must focus their report on a specific research question of choice, allowing individualisation within the whole-class dataset. Examples include comparing microbiome samples from different locations, or whether being a pet owner had influenced the microbial community of the human subject. Our students are provided with a clear assessment brief and mark scheme (Supplemental Information 4 and 5). This assessment method gives students training in an authentic research dissemination format, a common feature of CUREs (6). Alternative assessment formats could be a research poster, oral presentation or examination.

## Discussion

Our CURE provides students with inquiry-driven scientific training in modern molecular biology methods and bioinformatic analysis, using the human microbiome as a relevant research context. In Brownell and Kloser’s classification, our design would be considered as a ‘structured’ CURE as students do not have autonomy over the methods used, but have independence when it comes to data analysis and interpretation and communication of results (6). However, our CURE does not have a known answer, making it more typical of ‘open’ or ‘authentic’ CUREs.

### Evidence of Student Learning

The development of student technical skills is demonstrated through their experimental results (Figure 2). All but one group of students successfully PCR amplified the 16S gene from the frozen *E*.*coli* pellets (Figure 2A). Most groups prepared a genetic library that was successfully sequenced, demonstrating correct library preparation.

Demonstration of student understanding and bioinformatic competencies can be found in their experimental write-ups, particularly in the presentation and description of results (Figure 2). Reports were marked within the VLE by academics according to a structured mark scheme (Supplementary Information X), and a subset of reports were checkmarked by an independent academic member of staff. We used slightly different markschemes for the two cohorts, but there was no significant difference in marks between years (t-test t = 0.103, p = 0.92). The mean mark was 56.5% with a minimum mark of 21% and maximum of 84% (the pass mark was 40%). 42 of 49 students passed the assignment at the first attempt (86%), with the remaining students offered a later resit attempt. This indicates that the majority of the cohort met the learning outcomes.

### Student perceptions

We were also interested in student perceptions of the CURE so conducted a short survey, which was completed by 38 out of the 49 students from two cohorts. 3 students completed the survey but did not give consent so were removed from the dataset, resulting in a total of 35 participants (71% response rate). As such responses may not represent the whole class, but capture the majority of students. Ethical oversight and data analysis are described in detail in Supplementary Information 6 alongside the questions.

We first asked students what they had learned from the practicals as a free text question. Students identified techniques that they had learned (e.g. PCR, gel electrophoresis), but also identified more general scientific skills such as accuracy, safe working and analysis of results (Figure 3). Some students also commented on how the hands-on work had been more beneficial to their understanding than theoretical content:

**Figure 3:**
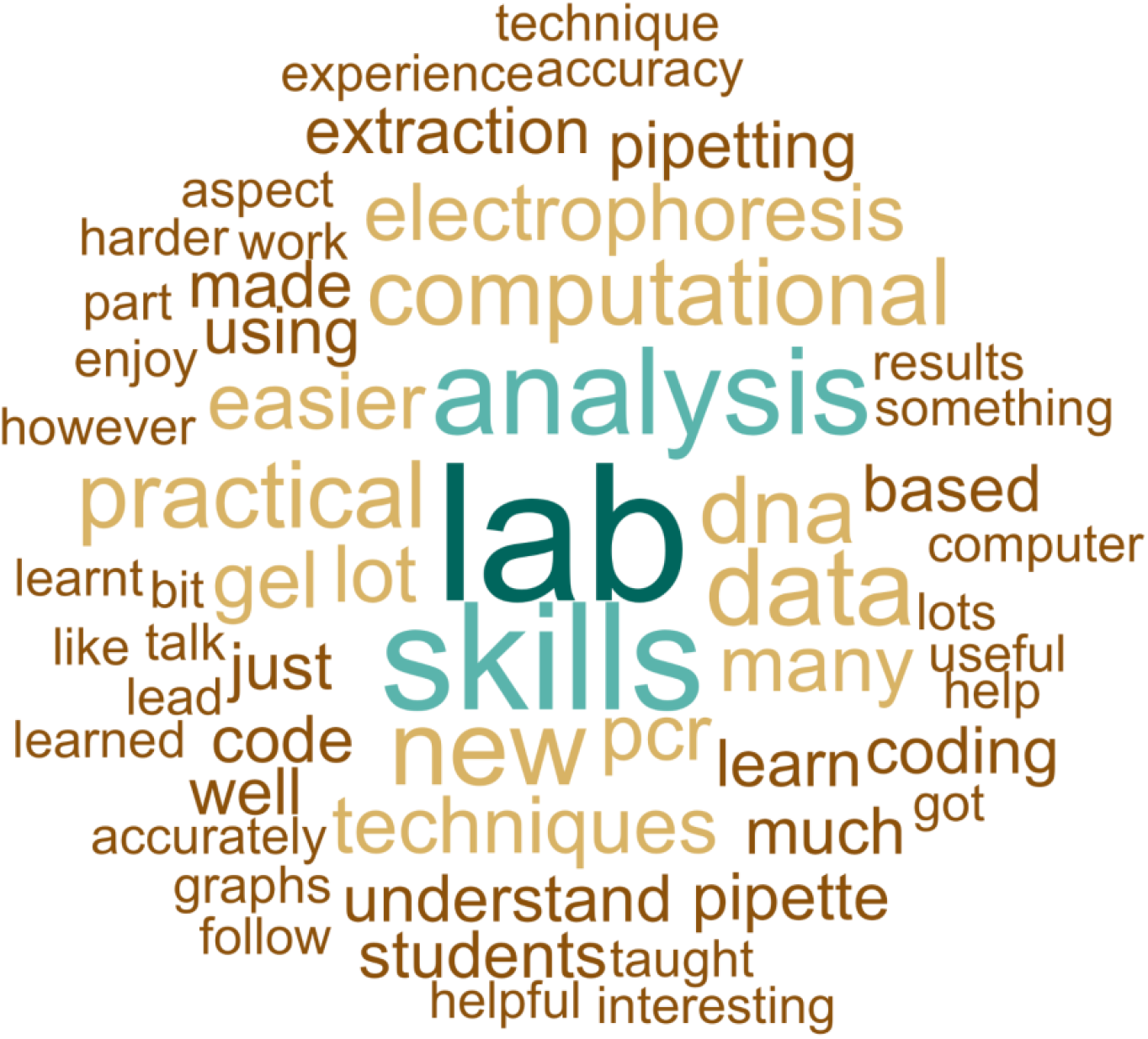
Student identification of what they learned from the CURE. Words are presented in proportion to the frequency with which they were mentioned. Words included if mentioned by 2 or more students.

> “PCR reaction, pipetting skills, running gels, working with tiny amounts of DNA - needing accuracy. Learning a new bit of code, exposed to python, vegan and other packages”
>
> “I learned a lot about how an experiment is actually conducted, rather than just learning or reading about them in a textbook.”

We then evaluated student perceptions of their learning aligned to Bloom’s Revised Taxonomy (39), using a series of previously validated Likert questions (40). Bloom’s taxonomy defines six increasingly advanced cognitive levels (Remember, Understand, Apply, Analyse, Evaluate and Create) and is commonly used to assess student learning. There were high levels of agreement (60% or above) for all levels of the Taxonomy with the exception of ‘Create’ (Figure 4A). The highest levels of ‘Strongly agree’ were for Understand (31%) and Analyse (29%). Students therefore agreed that the CURE had developed both lower and higher level learning.

**Figure 4:**
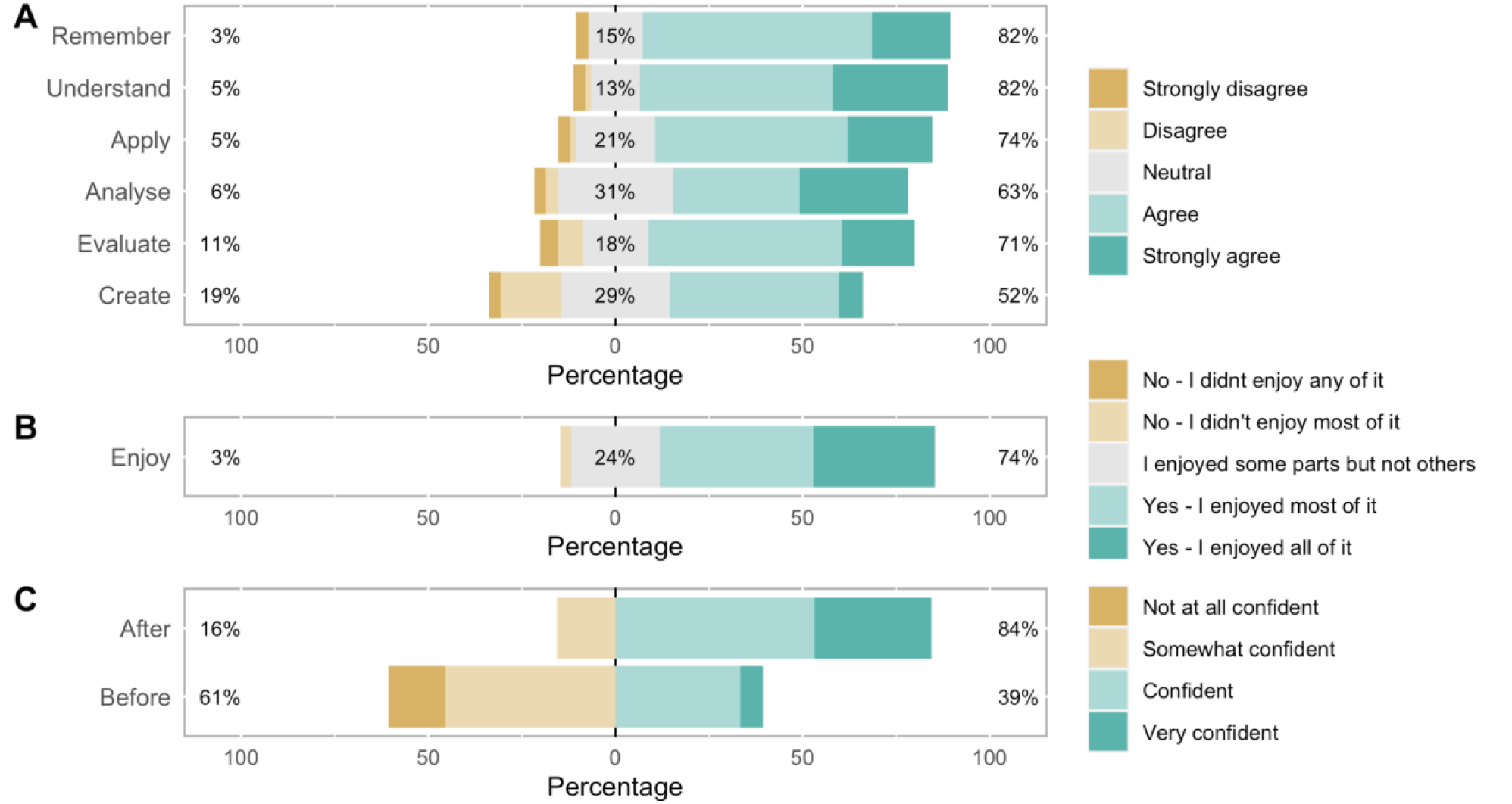
Student perceptions of the CURE labs. A: Student learning aligned to Bloom’s revised taxonomy. B: Student enjoyment of the labs. C: Change in student self-reported confidence in their laboratory work as a result of doing the CURE. Data presented for the 30 students who gave answers to all likert prompts. For the Bloom’s taxonomy questions (A) each student answered two questions which were combined, so n = 30×2 = 60 responses for each question.

We then asked students if they had enjoyed the CURE. 74% said they had enjoyed all or most aspects of the practicals (Figure 4B). After completion of the CURE there was also a significant increase in self-reported confidence (Figure 4C). 39% said they felt confident or very confident about lab work before doing the course, increasing to 84% after the course (Wilcoxon paired rank sum test V = 0, p-value <0.01).

### Potential Modifications and Extensions

The methods used are appropriate for any microbiome study, so could be used in the context of food hygiene, environmental microbiology, human or animal health. Students could also potentially sample their own microbiomes. Future iterations of our CURE may include these alternatives to give students more independence over their research questions, but will still use the same methods and workflow. It could also include classical microbiology (e.g. streak plates, overnight cultures) to compare with modern sequencing methods. We have also used a shortened version of these labs as an outreach activity with local schools. In this format students undertake a 2.5 hour practical where they perform gel electrophoresis of pre-prepared 16S PCR products, have a demonstration of the Flongle and then explore an example microbial community analysis through an interactive display, identifying dominant taxa in each sample.

## Conclusion

Our CURE takes undergraduate students with little or no experience of modern molecular methods to research-led inquiry within a short timeframe. It is designed in a format that is scalable and enjoyable for students and staff. Our CURE is embedded into a compulsory lower level undergraduate class, providing an inclusive introduction to research-led practical work. The CURE provides students with authentic training in modern molecular biology in a supported environment, equipping them to understand and engage with the potential of next-generation sequencing.

## Supporting information

Supplementary 4: Report instructions

Supplementary 5: Report marking scheme

Supplementary 6: Questionnaire methods

Supplementary 1: Genetic analysis lab manual

Supplementary 2: Bioinformatics manuals

Supplementary 3: Risk assessment

## Acknowledgements

We are thankful to the technical staff at our institution who have provided invaluable support in the development and running of these practicals. We also thank all students involved for their level of engagement and constructive feedback. We thank the VIPER High Performance Computing facility of the University of Hull.

