## Supplementary 4: Report instructions for "Who Grows There? A Course-based Undergraduate Research Experience to explore the human microbiome through 16S DNA metabarcoding"

### Supplemental Material 4:

### Microbiome Report Assignment Instructions

Produce a report on the results from the data generated during the practical. This will be in the format of ***a scientific report*** and will have the following sections:

**Introduction:** Background to the human skin microbiome and its diversity. How DNA metabarcoding can be used to detect this diversity. Brief outline of the experimental design and aims.

**Methods:** Description of the methods for generating the sequencing data. Include the experimental design and sampling strategy. Software and parameters used for bioinformatics and community analysis.

**Results:** Description of the results and sequencing outputs. All relevant figures should be included here.

**Discussion:** Findings are introduced. Results are interpreted, discussed and conclusions made.

**Referencing:** Harvard-formatted citations.

**Vital background information for the report can be found below**

### Microbiome practical methods and results summary

This forms an overview of the methods used and the results generated from the Microbiome practical. It is to be used as a reference point and is vital to the students who were unable to attend the practical. The layout of this document is deliberately sparse to ensure students look through the relevant materials available on Canvas for each week of the practical. These materials are linked where necessary. This is to ensure no student can copy and paste the following summaries and achieve a high mark - there is still effort on their part to achieve this.

### Methods

The following were specifically discussed during the practicals

##### Sample collection:

Equal sampling effort (time, pressure and swab rotation) with sterile swabs on 2cm square sections at each sample location.
Bioinformatics:
***fastp***: minimum read length = 1000
***fastp***: maximum read length = 1700
***Kraken 2*** standard Bacteria database was used for taxonomic assignment. See Kraken 2 manual for detailed information
***R*** data wrangling: only OTUs with ≥ 5 reads were kept per sample for final analysis
***R Vegan*** was used for OTU accumulation and community based analysis via NMDS

##### Bioinformatic analysis Jupyter Notebooks on GitHub

The links below and the relevant described sections are useful:
ONT_sequencing_data_analysis: fastp: "3.2.2. *fastp* qc of raw sequencing data"
R_analysis: Sequencing read depth threshold: "3.2. Cleaning the data"

### Results

##### DNA sequencing statistics

The details of the sequencing results are shown in the Genetic Analysis (500697) Week 7 module: Bioinformatics day 4 - slides 26 - 27. These slides show the outputs in the MinKnow software. There is some interpretation required.
There is more information in the "genetic_analysis_cleaned_data.xlsx" Excel file located in Genetic Analysis (500697) Week 7 module section "PRACTICAL OUTPUTS". This file is the final curated sequencing results. In the file, samples are columns, OTUs are rows (species first, then higher taxonomic ranks). Each cell is the number of reads of an OTU in each sample. The total number of assigned reads should be included in the report. To understand "OTU", see R_analysis: "1.2. Exploring the data" or Google the term.

##### R figures

Graphical outputs from the ***R*** analysis and loose descriptions and notes for each.
OTU richness per sample per site sampled:

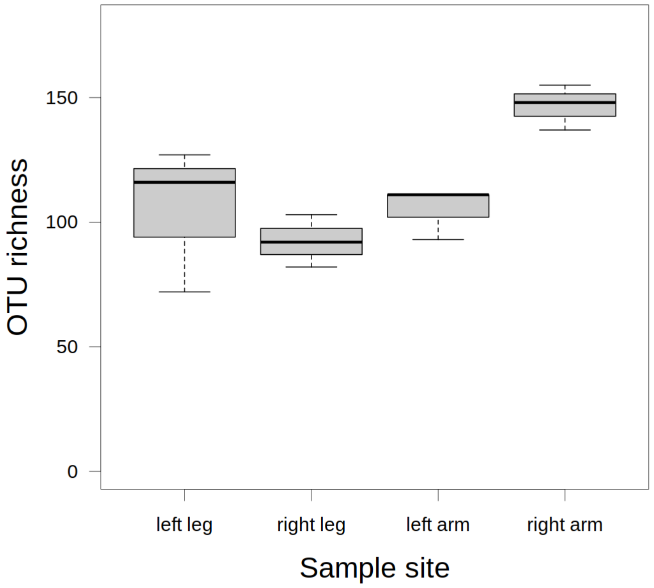


Total OTU richness per site sampled:

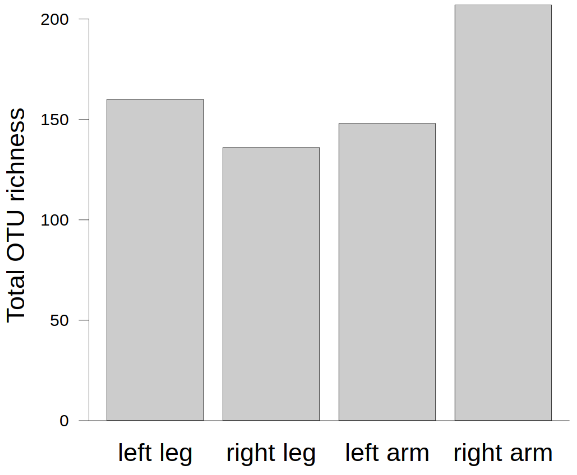


OTU accumulation:

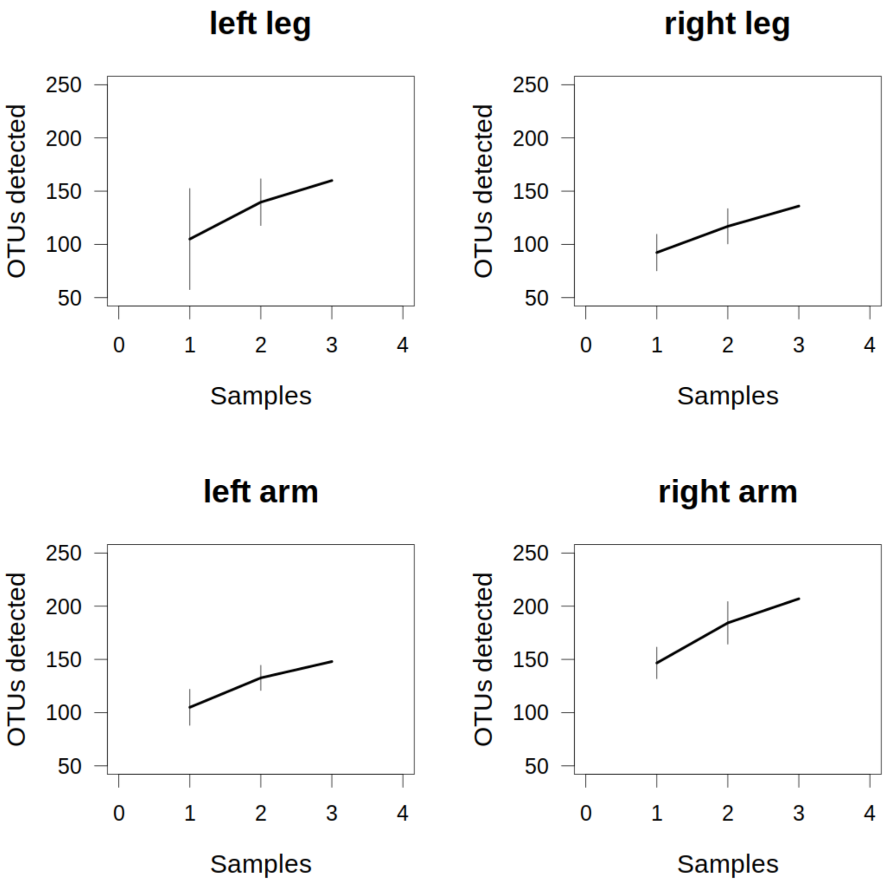


**Notes:** The above figure was generated from the data with **R Vegan** and shows the accumulated total OTUs per sample size. The vertical lines indicate standard deviation (*SD*) at each sample size.

**Important:** For the purpose of sampling completeness, a point to consider in the report, do the curves look like they have reached a plateau or would more samples be needed? Is it sufficient, i.e. how far from a plateau is it? Consider the size of the sample area - would it be feasible? Google “Species accumulation” for more information.

Non-metric multidimensional scaling ordination (NMDS):

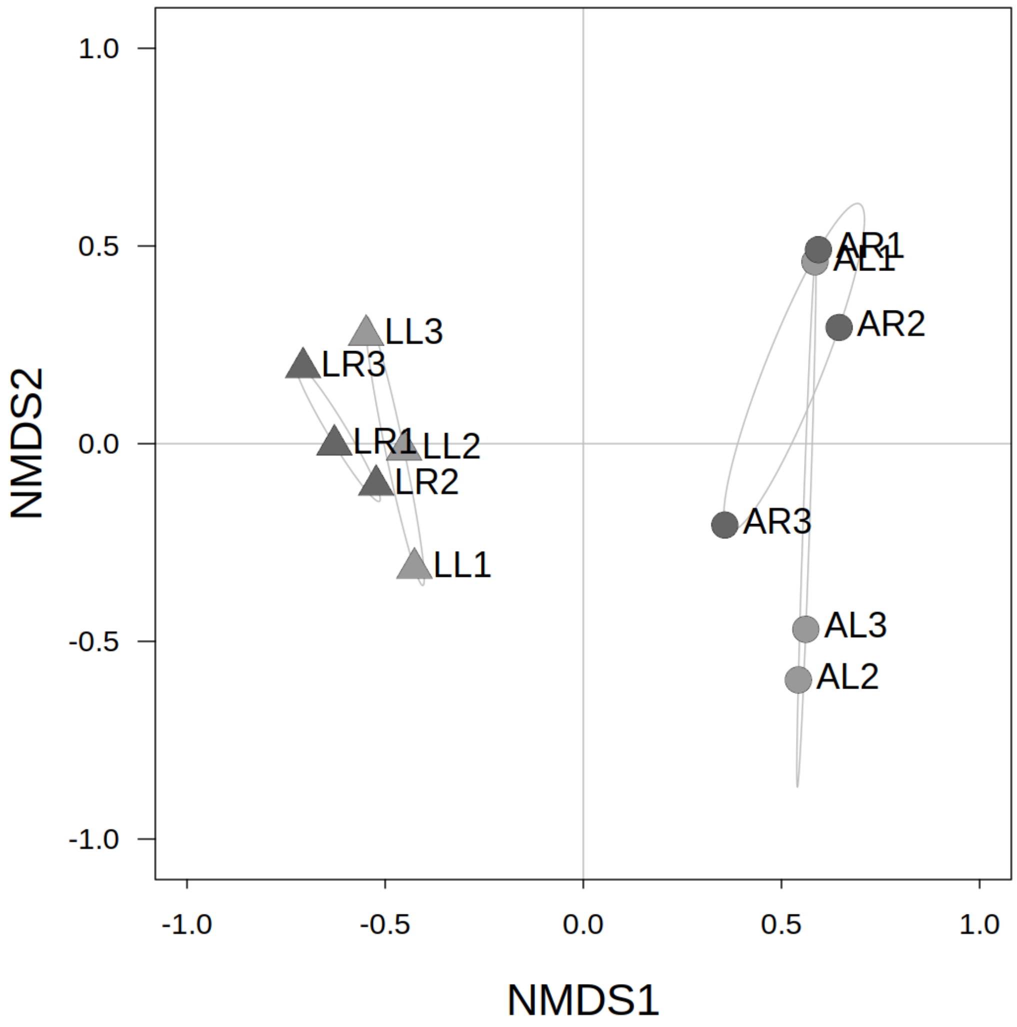


**Notes:** The figure was created from results generated from the data with ***R Vegan***. It shows the difference in community composition between each sample. The distance between each point shows the level of bacterial community difference. Google how to interpret it correctly.

#### Narrative

The results should not just be based solely on those above. There is much more information in the "genetic_analysis_cleaned_data.xlsx" Excel file (see above). Students should look through it and find any OTUs of interest, those that show a strong difference between arm and leg sample sites or similar are a good starting point. These could be used as the basis for a "story" - an underlying narrative for the report. Ideally, this would, ideally, reflect the known life history of the test subject (See Genetic analysis wrap-up - slide 12).

Interesting points were made by guest speaker Merideth Freiheit, a PhD student studying the human microbiome.
High diversity = good
*Cutibacterium acnes* - causes acne, common
*Micrococcus luteus* - common to the mouth, nose, and skin

Porphyromonas gingivalis, Capnocytophaga gingivalis - oral, cause gingivitis
*Staphylococcus aureus* - common on skin, opportunistic - MRSA (clinical nurse)
*Dolosigranulum pigrum* - upper respiratory (clinical nurse)
Streptococcus canis - dogs
Moraxella osloensis - Google it
Anaerococcus prevotii, Finegoldia magna - normal to skin microbiome

Students are encouraged to Google the results and gain an understanding of the complexity of the human microbiome. Students should then find relevant articles to cite for the report. **It should form a minor narrative aspect of the report**. Consider this aspect briefly in the introduction, report on it in the results, and expand on it in the discussion. A minimal approach is advised here, choose a single aspect - it should not overshadow the primary results.

The information contained here is deemed sufficient for all students to complete a suitable report. Those students that attended the practicals, this is to be used as a reminder. Those that could not attend, use the information provided here as a basis to gain an understanding of the analysis performed. All the above figures are available in the Genetic Analysis (500697) Week 7 Module, marks will not be deducted for their use.
