## Supplementary 5: Report marking scheme for "Who Grows There? A Course-based Undergraduate Research Experience to explore the human microbiome through 16S DNA metabarcoding"

### Supplemental Information 5:
### Who Grows There Genetic Analysis Report Markscheme

| Criteria | High 1st | 1st | 2i | 2i | 3rd | Fail | No marks |
|---|---|---|---|---|---|---|---|
| **Introduction (10%)** | **10 Pts** | **8 Pts** | **6 Pts** | **5 Pts** | **3 Pts** | **1 Pts** | **0 Pts** |
| Background to the human skin microbiome, its diversity between individuals and different body areas of an individual. How DNA metabarcoding can be used to detect this diversity. Brief outline of the experimental design and aims. | An exceptionally clear and accurate background showing a good understanding of the human skin microbiome diversity. Clearly describes and demonstrates the application of DNA metabarcoding to the field of study. Concise overview of the experimental design and aim of the project. | A clear background showing an understanding of the human skin microbiome diversity. Describes the application of DNA metabarcoding to the field of study. Clear overview of the experimental design and aim of the project. | A good background to the human skin microbiome diversity. Describes the application of DNA metabarcoding to the field of study. Overview of the experimental design for the project may be slightly incomplete. | Includes a background of the human skin microbiome diversity but it is not in great detail. Describes the application of DNA metabarcoding to the field of study but may miss some points. The experimental design overview for the project is incomplete or shows little rational. | Background is incomplete missing some key points. DNA metabarcoding is described but with little understanding. The experimental design overview is not present. | Background is superficial. DNA metabarcoding is not described well. The experimental design overview is not present. | Section is missing or contains nothing of relevance. |
| **Methods (30%)** | **30 Pts** | **26 Pts** | **21 Pts** | **16 Pts** | **10 Pts** | **5 Pts** | **0 Pts** |
| Introduction to the experimental design and sampling strategy. Description of the methods for generating the sequencing data. Software and parameters used for bioinformatics and community analysis. | The experimental design is introduced fully. The test subject's life history is described in detail. Sample locations and methodology are clearly explained. All methods for sequencing data generation are clearly described. All software used in the analysis is clearly stated, all parameters are described and includes any rational where applicable. Citations are included correctly for all methods. Methods are presented in the correct order and have a logical structure. | The experimental design is introduced acceptably. The test subject's life history is mentioned. Sample locations and methodology are explained. All methods for sequencing data generation are described. All software used in the analysis is clearly stated, parameters are described and some rational is present. Citations are included for all methods. Methods are presented in the correct order and have logical structure. | The experimental design is loosely introduced. The test subject's life history is mentioned, but some details are missing. All methods for sequencing data generation are described acceptably. All software used in the analysis is stated, parameters are described. Citations are included for most methods. Methods are presented in the correct order. | The experimental design is introduced very loosely or have large omissions. The test subject's life history is not clearly described. Methods for sequencing data generation are not described clearly. Most software used in the analysis is stated, some parameters are mentioned. Citations are included but may be missing for some methods. Methods are not all presented in the correct order. | The experimental design is poorly introduced if at all. The test subject's details are missing or very poorly described. Methods for sequencing data generation are mentioned but not described well enough. Some software used in the analysis is stated, parameters may not be present. Citations may not be present. Methods may be jumbled. | The experimental design is not present. No mention of the test subject' details. Methods for sequencing data generation are not clear or missing. Some software used in the analysis is stated, parameters are missing or not suitably stated. Citations are not present. Methods may have no structure. | Section is missing or contains nothing of relevance. |

| Results (25%) | 25 Pts | 22 Pts | 18 Pts | 14 Pts | 9 Pts | 4 Pts | 0 Pts |
|---|---|---|---|---|---|---|---|
| Description of the results and sequencing outputs. All relevant figures are included. | Results are described correctly and clearly in a succinct manner. Sequencing statistics are included and bioinformatic results are stated. Taxonomic assignment success is described. All relevant figures are included along with additional descriptive ones. All figures are clearly referred to correctly in text. Figure legends are correctly formatted and describe the figure clearly. Figure design and colour is well considered. | Results are described clearly in a succinct manner. Sequencing statistics are included and bioinformatic results are stated. Taxonomic assignment success is described. All relevant figures are included, may have some additional descriptive ones. Figures are clearly referred to in text. Figure legends describe the figure clearly. Figure design and colour is well considered. | Results are described acceptably. Sequencing statistics and bioinformatic results may be stated, but not correctly. Taxonomic assignment success may be present. All relevant figures are included. Figures are mostly referred to in text. Figure legends acceptably describe the figure. Figure design and colour is acceptable. | Results are described but may have omissions. Sequencing statistics and bioinformatic results are not present or may be unclear. Taxonomic assignment success is not clearly stated. All relevant figures are included. Some figures are referred to in text. Figure legends are present, but may not be clear. Figure design and colour has been poorly considered. | Results are poorly described but have many omissions. Sequencing statistics and bioinformatic results are not present. Taxonomic assignment success is not stated. Some figures may be missing. Remaining figures may not be referred to in text. Figure legends are not clear. Figure design has not been considered at all. | Results are poorly described or in a jumbled manner. Sequencing statistics and bioinformatic results are not present. Taxonomic assignment success is not stated. Figures may be missing. Figures are not referred to in text. Figure legends are incomplete. | Section is missing or contains nothing of relevance. |

| Discussion (20%) | 20 Pts | 16 Pts | 12 Pts | 10 Pts | 6 Pts | 2 Pts | 0 Pts |
|---|---|---|---|---|---|---|---|
| Findings are introduced. Results are interpreted, discussed and conclusions made. | Findings are introduced clearly. Results are interpreted and discussed in relation to the literature. Shortcomings of the experimental design are considered. Concludes well with a final overview and potential future direction. | Findings are introduced well. Results are interpreted acceptably and discussed in some relation to the literature. Some shortcomings of the experimental design are mentioned. Concludes with a final overview. | Findings are introduced acceptably. Results are interpreted but not always in relation to the literature. Experimental design loosely critiqued. Any conclusions are superficial. | Findings are introduced loosely. Results are interpreted but not clearly or in relation to the literature. Experimental design is not considered. Concludes abruptly. | Findings are not introduced well, if at all. Results are interpreted poorly. Experimental design is not considered. No clear conclusions. | Findings are not introduced. Results are interpreted poorly or in a jumbled manner. No conclusions. | Section is missing or contains nothing of relevance. |

| References (10%) | 10 Pts | 8 Pts | 6 Pts | 5 Pts | 3 Pts | 1 Pts | 0 Pts |
|---|---|---|---|---|---|---|---|
| A good number of relevant peer-reviewed scientific sources are cited; the in-text citations and reference list are correctly formatted in Harvard style. | Fully supported by peer-reviewed references in Harvard format within text. | Mostly supported by peer-reviewed references in Harvard format within text. | For the most part supported by peer-reviewed references in Harvard format within text, though some statements are not. | Some statements supported by peer-reviewed references within text. | Few statements supported by peer-reviewed references within text. | Almost no statements supported by peer-reviewed references within text. | Section is missing or contains nothing of relevance. |

| Structure and writing (5%) | 5 Pts | 4 Pts | 3 Pts | 2.5 Pts | 1.5 Pts | 0.5 Pts | 0 Pts |
|---|---|---|---|---|---|---|---|
| Structure and writing style of the report. Clear, scientific and concise. | Clearly structured with subheadings where needed. Writing style is scientific and has a good flow between sections. | Well structured and writing style is scientific and has a good flow between sections. | Structure is acceptable. Writing style may lack some flow but is otherwise clear and sufficiently scientific. | Structure may be lacking. Writing style may lack flow but is mostly clear and acceptably scientific. | Structure may be jumbled and some parts may be irrelevant. Writing style lacks flow but is not suitably scientific. | No, or very poor structure. Some parts may be irrelevant. Writing style is not very clear or scientific. | Section is missing or contains nothing of relevance. |
