## Supplementary 6: Questionnaire methods for "Who Grows There? A Course-based Undergraduate Research Experience to explore the human microbiome through 16S DNA metabarcoding"

### Supplementary Information 6: Questionnaire details

#### Ethical Oversight

Ethical approval for the survey was granted by the Faculty of Science and Engineering ethics committee (Project FEC_2023_21, Amendment FEC_2023_88). Participation in the survey was voluntary, and no personal identification or demographic data was collected to protect student anonymity. Study information was provided before the questionnaire, and students indicated consent for participation via a compulsory question. All data was stored in accordance with local data protection standards.

#### Data analysis

Data was analysed and visualised in RStudio (1) using the packages tidyr (2), dplyr (3), ggplot2 (4), cowplot (5), psych (6) splithalfr (7) and likert (8). Data was assumed to be non-normally distributed, so non-parametric statistics were used throughout.

##### Bloom's Taxonomy

Bloom’s revised taxonomy (9) is a commonly used framework for evaluating student learning in a variety of settings including undergraduate contexts (10). The six levels of the taxonomy (Remember, Understand, Apply, Analyse, Evaluate and Create) capture increasingly sophisticated cognitive processes. Bloom’s taxonomy must be applied in a local context; what ‘analyse’ looks like for a graduate level biochemistry course will be different to a high school level science class (10). Remember and Understand are usually considered ‘lower’ order thinking, and Analyse, Evaluate and Create considered ‘higher’ level thinking. Apply can be thought of either higher or lower depending on the specific context; in this case we consider it to be intermediate.

We asked two Likert style questions per level of the taxonomy, which we converted to ordinal data for quantitative analysis . The twelve Likert questions relating to Blooms’ taxonomy had a Chronbach alpha value of 0.932 indicating high levels of internal consistency. We also calculated Spearman-Brown reliability coefficients for each pair of questions (Table S1), as recommended for two-item scales by Eisinga et al. (11). Given the high internal consistency, for the 30 students who answered all Likert questions we combined each pair of questions per level of the taxonomy, giving 60 responses for each level represented graphically in Figure 4.

Table S1: Internal Consistency Analysis for the twelve Blooms Taxonomy Questions (n = 30)

| Taxonomy Level | Lower/ Higher Learning | Question Prompt | Internal Consistency |
|---|---|---|---|
| Remember | Lower | Remember concepts relevant to my course<br>Memorise experimental techniques | 0.764 |
| Understand | Lower | Understand concepts relevant to my course<br>Appreciate how experimental techniques work | 0.793 |
| Apply | Lower/ Higher | Apply knowledge from my course to a new situation<br>Use information from my course to understand a new situation | 0.865 |
| Analyse | Higher | Analyse data using techniques from my course<br>Perform numerical or graphical analysis of data | 0.939 |
| Evaluate | Higher | Evaluate the success of the experimental technique used<br>Identify the strengths and weaknesses of an experimental technique | 0.863 |
| Create | Higher | Create a new method to answer a scientific question<br>Design a new experimental strategy to investigate a scientific question | 0.793 |
| **All levels (Chronbach alpha)** | | | **0.932** |

### Questions

Purple text indicates answer format

1. If you give permission to participate in this study, please indicate using the options below: [YES – I understand the purpose of the study and how my data will be used. I give consent to participate; NO - I do not give consent to participate]
2. Which degree programme are you on? [List of relevant degree programmes]
3. Which module have you been asked to complete this questionnaire in? [List of relevant modules]
4. Did you enjoy doing the practicals in this module? Include any lab work or computational practicals [Yes - I enjoyed all of it; Yes - I enjoyed most of it; I enjoyed some parts but not others; No - I didn't enjoy most of it; No - I didn't enjoy any of it]
5. What do you think you learned in laboratory based practicals in this module? Include both lab work and computational aspects of the practical. [Free Text]
6. To what extent do you agree with the following statements: [Strongly Disagree; Disagree; Neutral; Agree; Strongly Agree] The practicals helped me to:
    a. Remember concepts relevant to my course
    b. Memorise experimental techniques
    c. Understand concepts relevant to my course
    d. Appreciate how experimental techniques work
    e. Apply knowledge from my course to a new situation
    f. Use information from my course to understand a new situation
    g. Analyse data using techniques from my course
    h. Perform numerical or graphical analysis of data
    i. Evaluate the success of the experimental technique used
    j. Identify the strengths and weaknesses of an experimental technique
    k. Create a new method to answer a scientific question
    l. Design a new experimental strategy to investigate a scientific question
7. How confident did you feel about performing practical work [Very confident; Confident; Somewhat confident; Not at all confident]
    a. before doing the practicals in this module
    b. after doing the practicals in this module
8. These questions are about your general opinions of laboratory work. Please indicate to what extent you agree with the following statements: [Strongly Disagree; Disagree; Neutral; Agree; Strongly Agree]
    a. Laboratory work is something that I enjoy
    b. I would rather have lectures than do laboratory practical work
    c. I always feel well prepared for laboratory practical classes
    d. Laboratory work teaches me valuable skills
    e. It would be better to be shown a video of a practical being done than to do it ourselves
    f. I would recommend laboratory practical classes to others
    g. I feel safe while undertaking laboratory work
9. If you have anything that you would like to comment, including any improvements you think we could make to the practicals, please use the box below. [Free text]
