## Supplementary 1: Genetic analysis lab manual for "Who Grows There? A Course-based Undergraduate Research Experience to explore the human microbiome through 16S DNA metabarcoding"

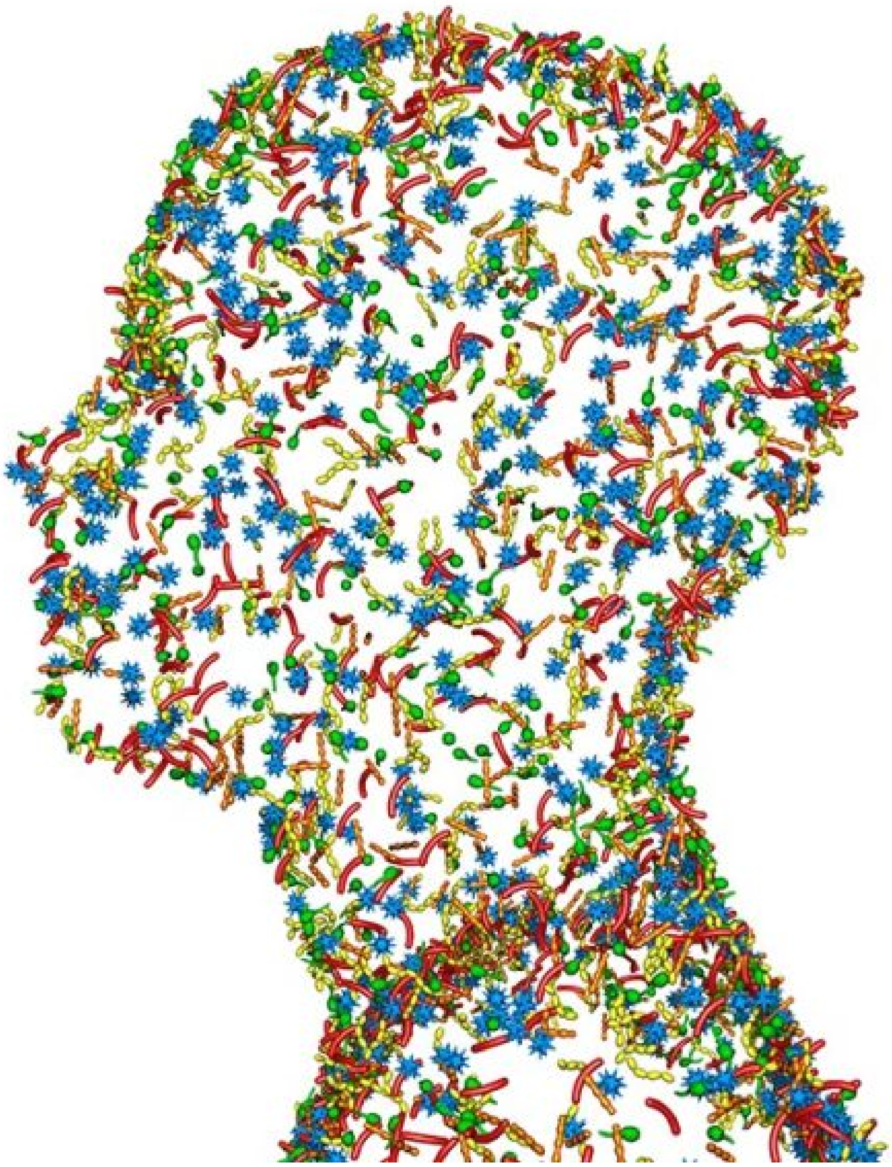

### Metabarcoding the Human Microbiome

Dr Graham S Sellers

### Genetic Analysis Lab Manual

#### Overview of Week 1

| **Day** | | **Activity Plan** |
|---|---|---|
| Monday | Morning | Introduction. Pipetting exercises. DNA extraction. |
| | Afternoon | |
| Tuesday | Morning | DNA quantification. PCR. |
| | Afternoon | |
| Wednesday | Morning | Gel electrophoresis. Basic calculations exercises. Recap. |
| | Afternoon | |
| Thursday | Morning | Sequencing library prep. pt1 (PCR). |
| | Afternoon | Sequencing library prep. pt2 (gel electrophoresis and quantification). |
| Friday | Morning | |
| | Afternoon | |

#### Achievements of Week 1

This week you will:

1. Extract purified DNA from a bacterial culture
2. Quantify the extracted DNA
3. Run a PCR using bacterial 16S markers
4. Analysed PCR results via gel electrophoresis and imaging
5. Use the above skills to prepare and quantify a genetic library for MinION sequencing

#### Overview

Bacterial 16S DNA metabarcoding of the human microbiome collected from skin swabs. This week we concentrate on generating the genetic data for computational analysis next week. DNA sequencing will be carried out using Oxford Nanopore's Minion sequencing platform, the most recent addition to next-generation sequencing technologies.

#### Aims and outcomes

Students will learn clinically relevant laboratory based skills through preparation and quantification of DNA sequencing libraries. This will include PCR preparation, gel electrophoresis, and spectrofluorometry. The sequencing data generated from this practical will be the basis for the computational analysis in the second week of the practical and the final project write-up.

### Genetic Analysis day 1

#### Overview

| Time | Activity |
|---|---|
| 09:00 - 09:30 | Introduction.<br>*Overview of the day, health and safety in the lab.* |
| 09:30 - 10:00 | Pipetting exercises.<br>*Best laboratory practices.*<br>*Level up your pipetting skills in preparation for the week ahead.* |
| 10:00 - 12:00 | DNA extraction.<br>*Important safety measures and best practices.*<br>*Isolate DNA from E. coli samples.* |

### Pipetting exercises

Unsurprisingly, the correct use of a pipette is an important skill in laboratory based genetics. The reagents used in many techniques must be accurately pipetted in tiny volumes, any errors can lead to the failure of the process. In some cases there may only be sufficient of a certain reagent or sample that is available, and to have this fail due to human error would not be favourable.

So, to ensure everyone is up to speed and can operate the Gilson pipettes that will be used throughout this week's practical, you will now perform some pipetting competency exercises.

#### The Gilson pipette

The ubiquitous pipette. Used in most laboratories worldwide.

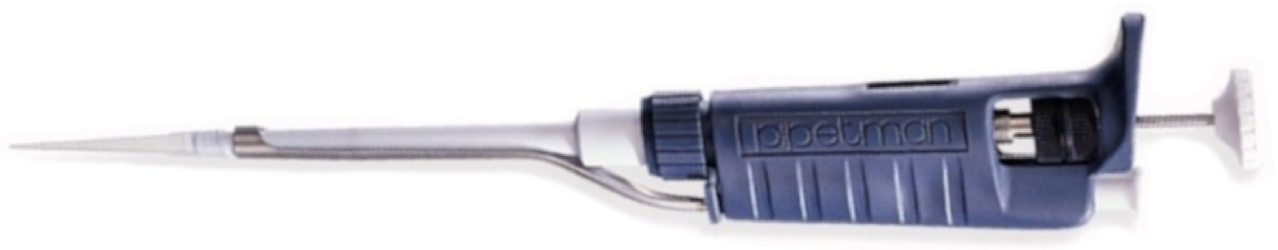


A quick refresher of microlitres (μl):

**1 ml = 1000 μl, or, 1 μl = 0.001ml**

The Gilson pipettes you will be using will be of these sizes:

**P1000** = max volume of **1000 μl** = 1 ml

**P200** = max volume of **200 μl** = 0.2 ml

**P100** = max volume of **100 μl** = 0.1 ml

**P10** = max volume of **10 μl** = 0.01 ml

#### Materials

In front of you you will find the following equipment:

- ☐ P1000, P200, P100 and P20 Gilson pipettes and pipette tips
- ☐ 1.5 ml tubes
- ☐ Purified water
- ☐ Parafilm strip

***NOTE:*** Check if everything required is present. If you cannot see or are missing any of the required materials, ask for assistance - we are here to help.

Now, if everything's in order, let's do the pipetting competency exercises on the following page.

### Pipetting competency exercises

You will now have some time to familiarise yourself with pipette use. Work your way through the following 3 exercises. Any issues, please ask for assistance.

***NOTE:*** These exercises may seem trivial, but each one is required later in this week's practical. Competency in pipette use will make all the difference.

#### 1. Volume control

To get you familiar with the increments and volumes of each pipette, you will practise transferring different volumes of water between tubes.

- ☐ Add 800 µl water to a 1.5 ml tube
- ☐ Add 180 µl water to the tube
- ☐ Add 55 µl water to the tube
- ☐ Add 15 µl water to the tube

Remove 1000 µl from the tube, how much water should there be left?

Now, take a pipette set to your expected value and remove that amount of water.

Is there any water remaining? Why?

#### 2. Target practice

There are many instances during laboratory work where accurate handling of the pipette is required, such as gel electrophoresis. Now, we're going to replicate this.

You will see there are some small squares drawn on the parafilm strip in front of you. Take a P20 pipette and practice loading 5 µl of water into this square. Do not touch the parafilm with your pipette tip while doing this. Do this a few times until you have the stability required.

Once you are comfortable doing this move onto the next exercise.

#### 3. Pixel art

It's time to write your initials in tiny droplets of water on the tape!

Take the P20 and fill it with 20 µl of water. Slowly push the pipette plunger down to release the smallest possible drop onto the parafilm. Do not touch the parafilm with your pipette tip while doing this.

#### Some questions

1. What's the fastest way to pipette 1.2 ml using a P1000?
2. Do you compress the pipette plunger to stop 1 or 2 prior to loading it with liquid?
3. What is the smallest volume you should pipette with a P200?

You should now be sufficiently good with a pipette.

---

***PAUSE:*** *Do not go ahead to the next section. Please wait until I have talked about the next stages.*

---

### DNA extraction

For any kind of genetic analysis, the first, and most important step, is extracting pure nucleic acids from the sample. In this practical we are interested in DNA as it is the basis of many molecular approaches for identification.

There are many methods for extracting DNA from samples, however, we will be following a modified version of the MU-DNA extraction protocol of Sellers et al. (2018). The extraction protocol can be broken down into simple steps as shown below.

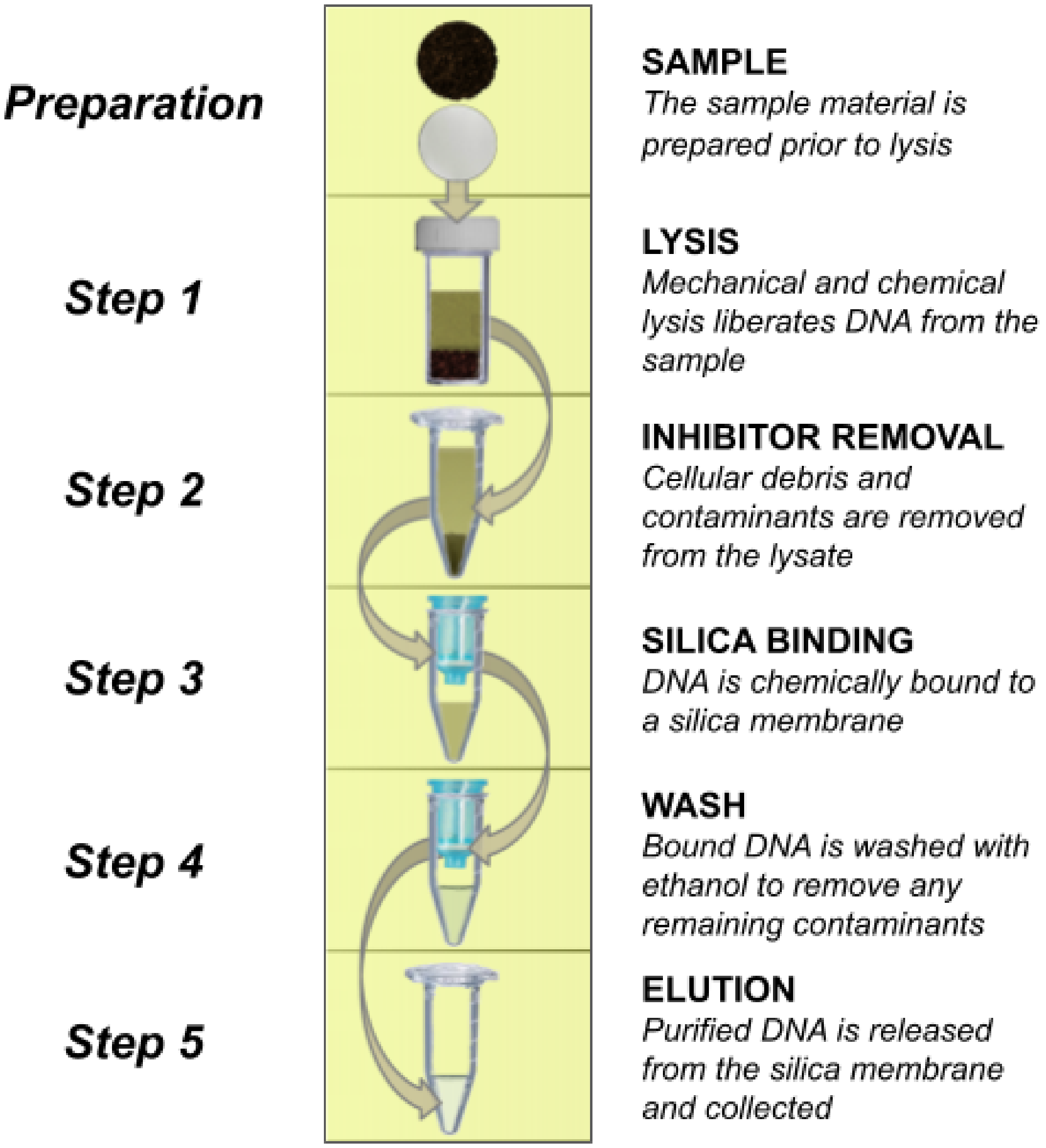

### Materials and methods

You will now perform a DNA extraction protocol on some specially grown liquid medium cultures of *Escherichia coli (E. coli)*. This is to allow you to experience DNA extraction - get a genuine, first-hand understanding of the procedure.

***NOTE:*** The *E. coli* samples you will be using have all been frozen at -20°C and are to be considered inactive (i.e. destroyed). They form no threat to your health, but, please treat them responsibly.

---

***DO NOT PANIC:*** You will be following a straightforward and established DNA extraction protocol. If you have any trouble/concerns, please ask. There are some (very) experienced demonstrators on hand (and myself), so feel free to approach them (or myself) for any assistance.

---

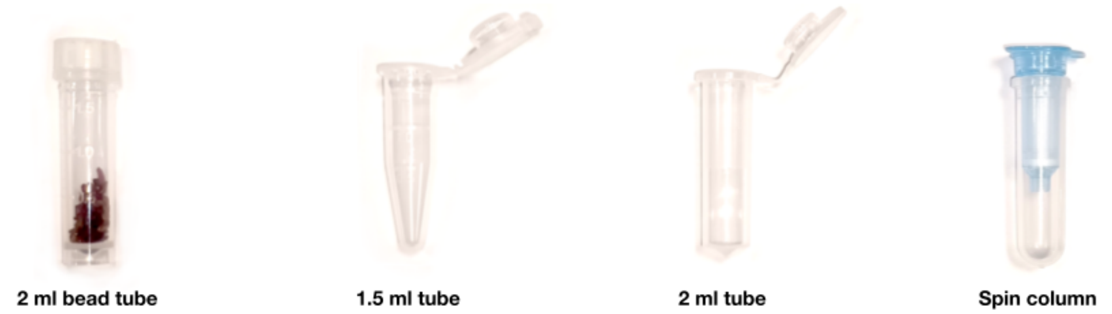


#### Materials

In front of you you will find the following equipment:

- [ ] P1000, P200, P100 and P20 Gilson pipettes and pipette tips
- [ ] Bead tubes: 2 ml bead tubes
- [ ] 1.5 ml tubes
- [ ] 2 ml tubes
- [ ] Spin columns and collection tubes
- [ ] Benchtop centrifuge
- [ ] Microcentrifuge
- [ ] Vortex mixer
- [ ] Chemical waste container

Additionally you will have:

- [ ] 250 µl liquid medium cultures of *E. coli* in 1.5 ml tubes
- [ ] Ice bucket
- [ ] Reagents for DNA extraction in 5 ml tubes:
    - [ ] ***Lysis Solution***
    - [ ] ***Lysis Additive***
    - [ ] ***Flocculant Solution***
    - [ ] ***Binding Solution***
    - [ ] ***Wash Solution***
    - [ ] ***Elution Buffer***

***NOTE:*** Check if everything required is present. If you cannot see or are missing any of the required materials, ask for assistance - we are here to help.

Now, if everything's in order, let's proceed with the DNA extraction protocol on the following page.

### DNA extraction protocol

A modified MU-DNA protocol for DNA extraction of liquid medium cultured *E. coli*.

#### PREPARATION

1. Centrifuge liquid medium culture of *E. coli* at 10,000 xg for 1 min
2. Remove and discard liquid medium without disturbing pellet
3. Add 50 µl ***Lysis Solution*** to the pellet, vortex to resuspend
4. This is now the ***Prepared Sample***

#### LYSIS

1. Add 550 µl ***Lysis Solution*** and 200 µl ***Lysis Additive*** to the 2 ml bead tube
2. Add 50 µl ***Prepared Sample*** to the 2 ml bead tube
3. Close lid tightly and vortex briefly to mix
4. Place the preparation in a tube rack and ask a demonstrator to check your progress

---

***IMPORTANT:*** *Once all sample preparations have reached this point, they will be taken to be processed on the TissueLyser II. Once your samples are returned, continue with the next steps.*

---

5. Centrifuge at 10,000 xg for 1 min at room temperature
6. Transfer 600 µl supernatant to a fresh 1.5 ml tube

#### INHIBITOR REMOVAL

1. Add 200 µl of ***Flocculant Solution***, vortex briefly and incubate at 4°C or on ice for 10 mins
2. Centrifuge at 10,000 xg for 1 min
3. Without disturbing the pellet, transfer 500 µl supernatant to a 2 ml tube

#### SILICA BINDING

1. Add 1000 µl ***Binding Solution***, vortex briefly to mix
2. Transfer 650 µl of the mixture to the spin column
3. Centrifuge at ≥ 10,000 xg for 15 secs, discard the flow-through from collection tube
4. Repeat steps 2 to 3 until all the mixture has passed through the spin column

#### WASH

1. Add 500 µl of ***Wash Solution*** to the spin column
2. Centrifuge at 10,000 xg for 15 secs, discard the flow-through from collection tube
3. Repeat steps 1 and 2 a second time
4. Centrifuge at 10,000 xg for 2 min, replace collection tube with a 1.5 ml tube

#### ELUTION

1. Add 100 µl of ***Elution Buffer*** directly to the spin column membrane and incubate for 1 min at room temperature
2. Centrifuge at 10,000 xg for 1 min
3. DNA is now in the 1.5 ml tube

***IMPORTANT:*** *Ensure you have labelled your samples correctly - you will need to find them tomorrow.*

### Day 1 skills

Today you have:

- Gained proficiency in accurate pipette work
- Followed a written protocol
- Extracted DNA from a bacteria sample

Aspects of these skills and processes form the bulk of all laboratory based work for genetic analysis.

### Week 1 achievements so far:

You have now got the following Week 1 achievement:

1. Extract purified DNA from a bacterial culture ✓

WELL DONE!

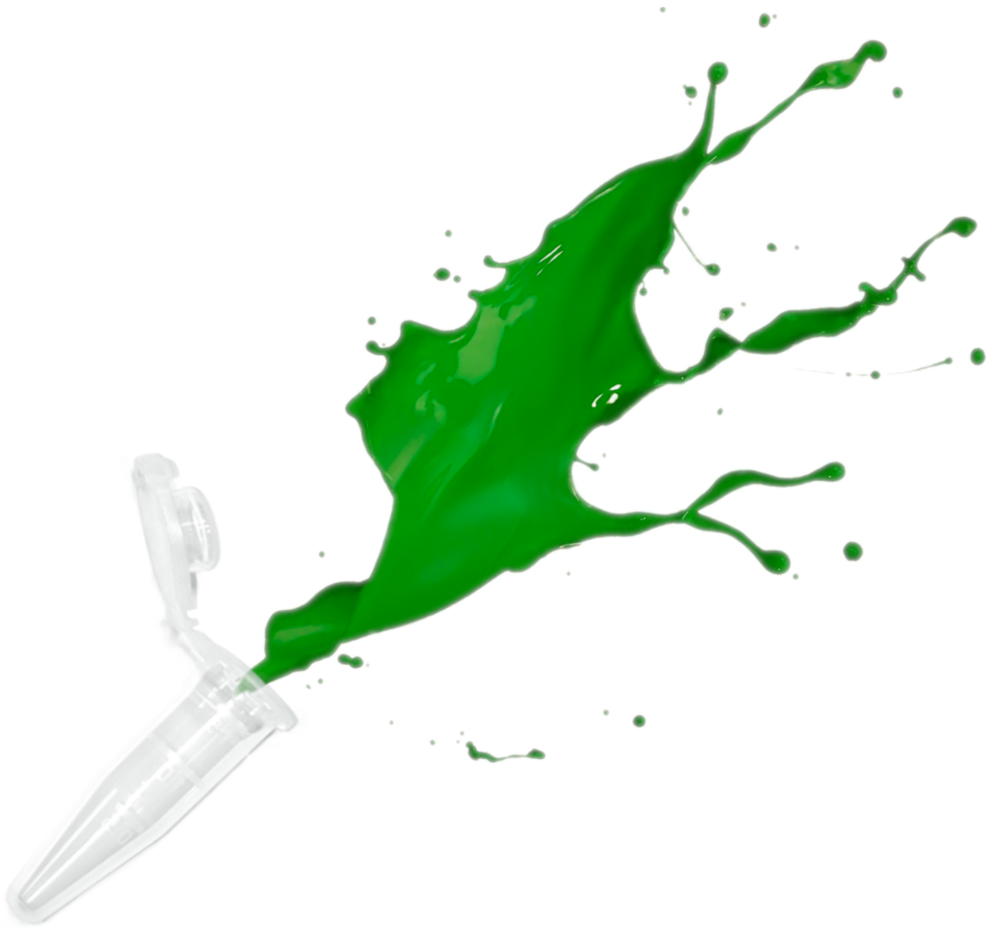

### Tomorrow

Quantification and PCR amplification of your DNA extractions.

### Genetic Analysis day 2

#### Overview

| Time | Activity |
|---|---|
| 09:00 - 09:30 | Introduction.<br>*Overview of the day, health and safety in the lab.* |
| 09:30 - 10:30 | DNA quantification.<br>*Best laboratory practices.*<br>*Using a Qubit fluorometer to see how much DNA you extracted.* |
| 10:30 - 12:00 | PCR.<br>*Important safety measures and best practices.*<br>*Amplification of bacterial 16S region via PCR.* |

### DNA quantification

For many processes in laboratory-based genetic work, knowing the concentration of the nucleic acids you are working with is essential. We are now going to measure the concentration of DNA extracted from your samples before we move on to today's next step.

DNA can be quantified in many ways, here we will use a Qubit fluorometer. The Qubit is used as an industry standard, it can accurately measure double stranded DNA (dsDNA) to concentrations as low as 0.005 ng/µl.

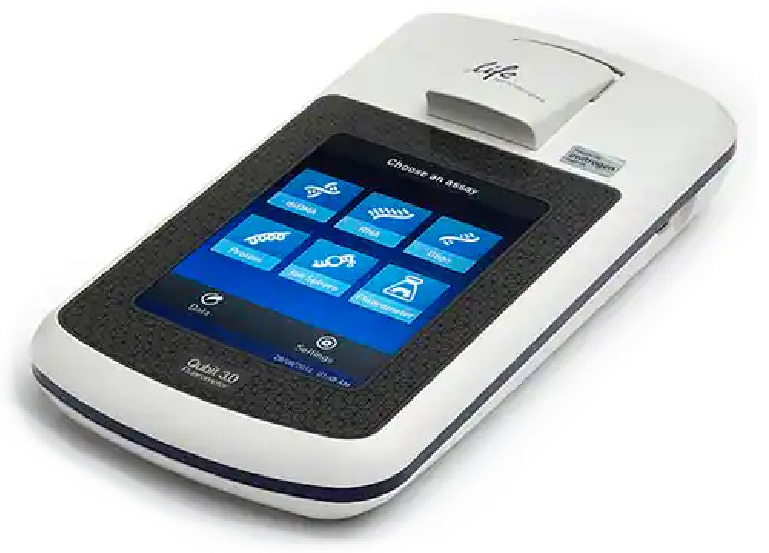


#### How it works

The Qubit uses fluorescent dyes that emit signals only when bound to the specific target molecules, in our case, dsDNA.

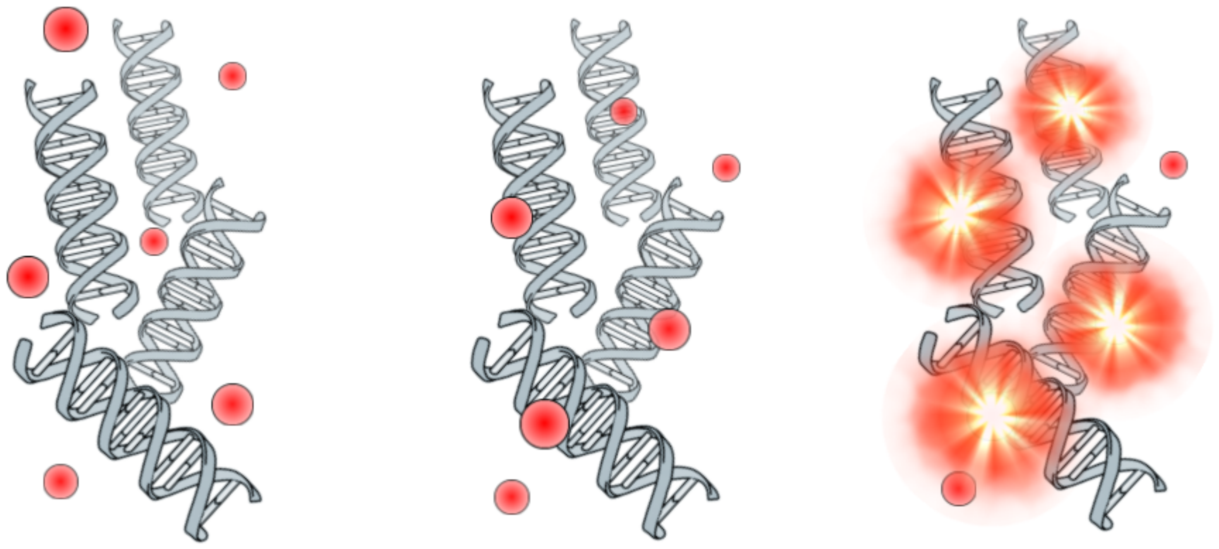

**DNA added to Qubit buffer**

**Qubit buffer dye binds to dsDNA**

**Bound dye fluorescence measured**

### Materials and methods

All the samples will be measured in a single batch using the same controls. This will make all the results directly comparable. To do this we have made a Qubit buffer master mix that you can take out the correct volume for your assay. We have also created the 2 controls required for the calibration prior to measuring the DNA concentration of your samples.

For completeness, here are the volumes required for the Qubit buffer and Qubit dye for a single sample. A larger master mix can easily be made by scaling the volumes and adding 10% extra for pipetting error.

Add 1 µl of Qubit dye to 199 µl Qubit buffer. Vortex to mix, spin down briefly.

You will quantify 2 µl of your DNA sample. The final volume used for Qubit quantification is 200 µl, so you will use 198 µl of Qubit buffer master mix (200 µl - 2 µl = 198 µl).

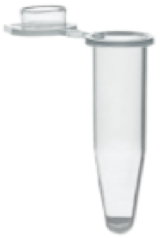

Qubit tube

#### Materials

In front of you you will find the following equipment:

- ☐ P1000, P200, P100, P20 and P10 Gilson pipettes and pipette tips
- ☐ Qubit tubes
- ☐ Microcentrifuge
- ☐ Vortex mixer

### DNA quantification protocol

1. Vortex DNA sample to mix thoroughly, spin down briefly
2. Transfer 198 µl Qubit buffer master mix to a Qubit tube
3. Add 2 µl of extracted DNA. Vortex to mix, spin down briefly
4. Incubate at room temperature for 2 mins

Once your sample is ready, ask a demonstrator and they will show you how to quantify it on the Qubit. Make a note of your sample’s DNA concentration - you will need it for later.

---

***PAUSE:*** *Do not go ahead to the next section. Please wait until I have talked about the next stages.*

---

### Polymerase Chain Reaction

Now we will prepare a Polymerase Chain Reaction (PCR). PCR utilises markers (aka primers) to replicate a specific region of DNA (aka amplicon). This could be a certain gene of interest or a potentially diagnostic sequence somewhere in an organism's genome. You will be using universal bacterial primers designed to amplify a fragment of the 16S rRNA region across most species of bacteria. The resulting amplicon will be ~1500 bp in length. This amplicon has sufficient differences between species to allow us to use it as an identification tool.

For metabarcoding, as the focus of this practical, PCR is an essential and powerful protocol for replicating large amounts of DNA. After the PCR has finished, our amplicon will be replicated to around 2 billion times its original concentration. Nice!

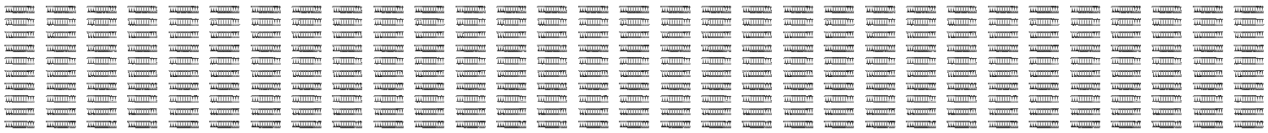

### Materials and methods

For current, modern, PCR reaction preparations we use a Taq DNA Polymerase pre-prepared buffer. This saves unnecessary pipetting, removes errors, and ultimately makes the process much simpler.

***NOTE:*** The reagents for PCR need to be kept cold. You will find the required PCR reagents in your ice bucket. Place your samples on ice too.

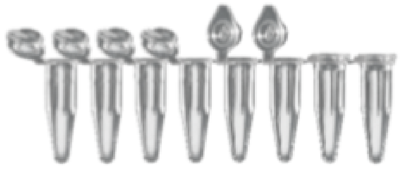

**0.2 ml PCR strip tubes**

#### Materials

In front of you you will find the following equipment:

- ☐ P1000, P200, P100, P20 and P10 Gilson pipettes and pipette tips
- ☐ 0.2 ml PCR strip tubes
- ☐ 1.5 ml tubes
- ☐ Ice bucket with PCR reagents
- ☐ Microcentrifuge
- ☐ Vortex mixer

PCR reagents in the ice bucket:

- ☐ Taq DNA Polymerase master mix
- ☐ Primers *27F* and *1492R*
- ☐ Molecular grade $H_2O$
- ☐ Positive control

***NOTE:*** Check if everything required is present. If you cannot see or are missing any of the required materials, ask for assistance - we are here to help.

### PCR preparation protocol

You will work together in your group to perform this step.

You will need to make a PCR reaction for each sample, plus a positive and negative control. If you are a group of 4, you will need to make a master mix for 6 reactions: 4 samples plus your positive and negative controls. However, as there are always pipetting errors, add in an extra reaction's worth so your master mix is for 7 reactions instead.

***NOTE:*** *We will be using molecular grade $H_2O$ for your negative control.*

Below is what is required for a single reaction:

1. 12.5 µl Taq DNA Polymerase master mix
2. 8.5 µl molecular grade $H_2O$
3. 1 µl primer 27F
4. 1 µl primer 1492R

**Total volume** = 23 µl

Calculate the volumes required for each reagent to create a PCR master mix of sufficient volume for your group.

---

***IMPORTANT:*** *Ask a demonstrator to confirm you have correctly calculated the volumes required before continuing.*

---

Once checked, create your PCR master mix in a 1.5 ml tube, label it, and place it on ice until required. Below is an example of the ideal layout for your samples in the PCR strip tubes:

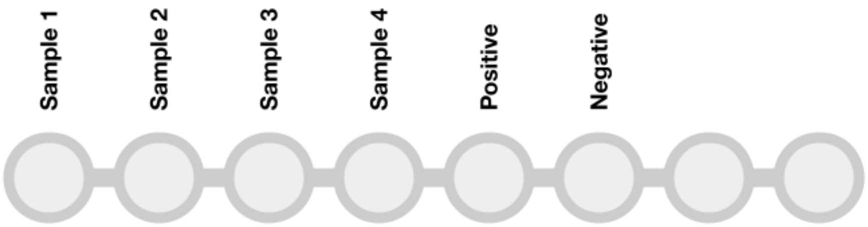


*Hint: pre labelling your tubes is a good idea*

Add 23 µl of PCR master mix to each tube in the PCR strip that is needed (samples plus controls). Now do the following:

1. Vortex all samples and positive control, spin down briefly, place on ice
2. Add 2 µl molecular grade H2O to the negative control PCR tube, close cap
3. Add 2 µl of each of your samples to the relevant PCR tube, close cap after each sample
4. Finally, add 2 µl of positive control to the positive control PCR tube, close cap
5. Ensure all caps on the PCR strip tube are closed securely
6. Gently flick PCR strip tubes to mix contents, spin down briefly, place on ice

Once completed let a demonstrator know. Make sure you have labelled your tubes clearly.

#### Now what?

Your PCR mixtures will be taken and placed in a thermal cycler. It's going to take 1 hr 30 mins for the PCR to be completed. Your final PCR product will be stored overnight in a fridge.

Just for completeness, here are the thermal cycler conditions used for the PCR:

***1 min @ 95°C, 30 X (15 secs @ 95°C, 15 secs @ 55°C, 30 secs @ 72°C), 7 mins @ 72°C.***

And here is the thermal cycler the PCR will take place in:

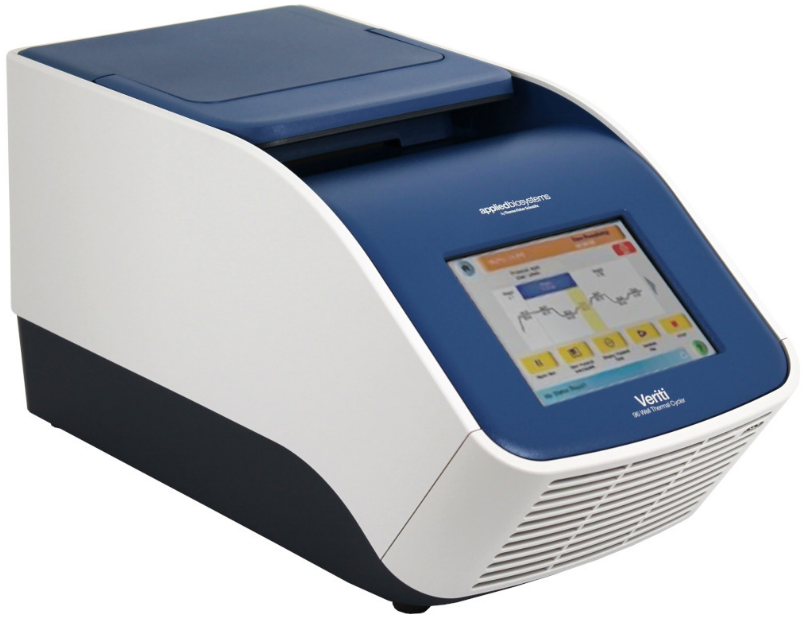


Applied Biosystems™ Veriti™ 96-well thermal cycler

### Questions

The following questions are designed to be a test of your knowledge. If you don't know, ask - this is just as good a way to learn. Just give it a go yourself first.

**Based on your Qubit results, what was the total yield of DNA (ng) of your sample?**

**During preparation, why was everything kept on ice?**

**Why were the samples added to the PCR tubes in the order they were?**

**Why were the final PCR reactions kept on ice prior to being placed on the thermal cycler?**

**Can you state what is happening at each stage of a PCR cycle?**

### Day 2 skills

Today you have:

- Quantified dsDNA
- Prepared a PCR

Again, these skills form the majority of laboratory work for genetic analysis.

### Week 1 achievements so far:

You have now got the following Week 1 achievements:

1. Extract purified DNA from a bacterial culture ✓
2. Quantify the extracted DNA ✓
3. Run a PCR using bacterial 16S markers ✓

WELL DONE!

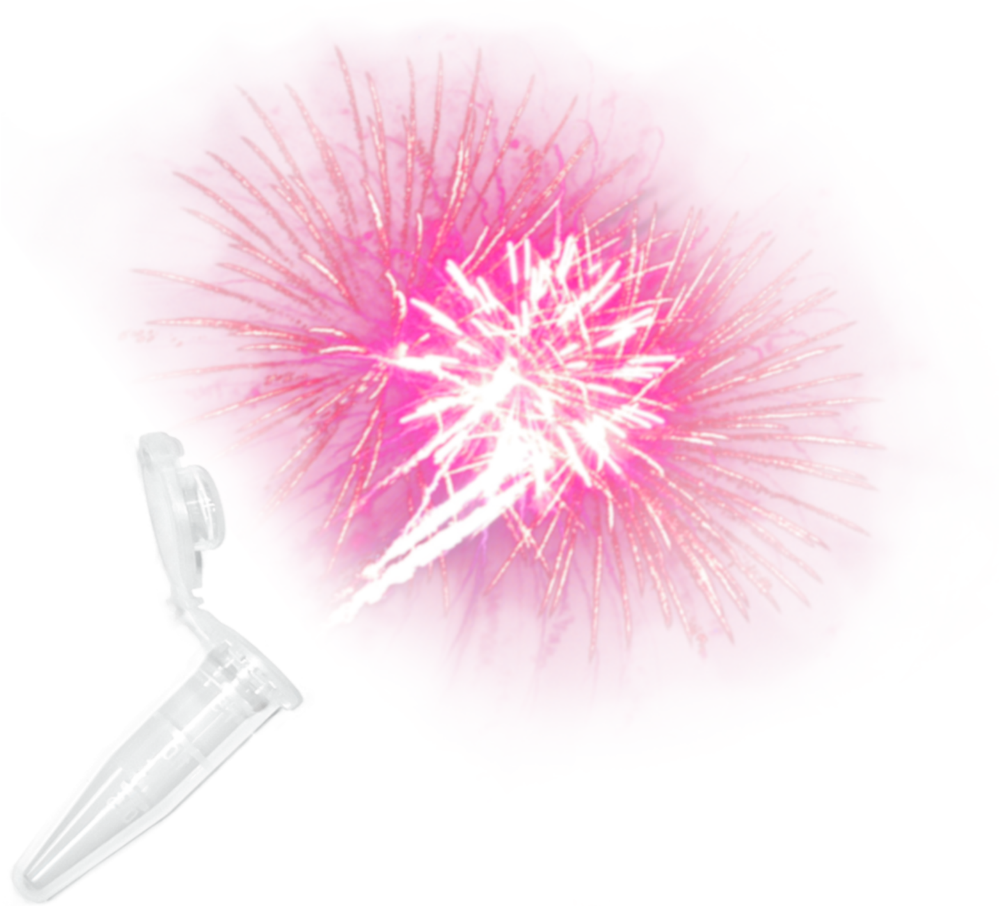

### Tomorrow

Gel electrophoresis and visualisation.

### Genetic Analysis day 3

#### Overview

| Time | Activity |
|---|---|
| 09:00 - 09:30 | Introduction.<br>*Overview of the day, health and safety in the lab.* |
| 09:30 - 10:30 | Gel electrophoresis.<br>*Loading and running a gel.* |
| 10:30 - 11:00 | Gel visualisation.<br>*Visualise gels on the UV imager.* |
| 11:00 - 12:00 | Skills roundup.<br>*Reflection on skills learnt so far.*<br>*Preparation for tomorrow.* |

### Gel electrophoresis and visualisation

You have, so far, extracted DNA from a bacterial culture. Quantified the extracted DNA. Prepared and run a PCR, using universal bacterial 16S primers, with the extracted DNA.

After all this effort we still don't know if anything has worked - not a single thing we have done so far was visible to the human eye, nor did it give any evidence it was there or something was happening. Well, with the exception of those numbers on the Qubit, right?

The success of the PCR is most important for this practical, so, let's see what needs to be done.

#### Did it work?

We must visualise the PCR product to see if PCR amplification was successful. This is a two stage process. First, we must perform gel electrophoresis on your PCR products. Then we need to image the gel to see the outcome of the gel electrophoresis.

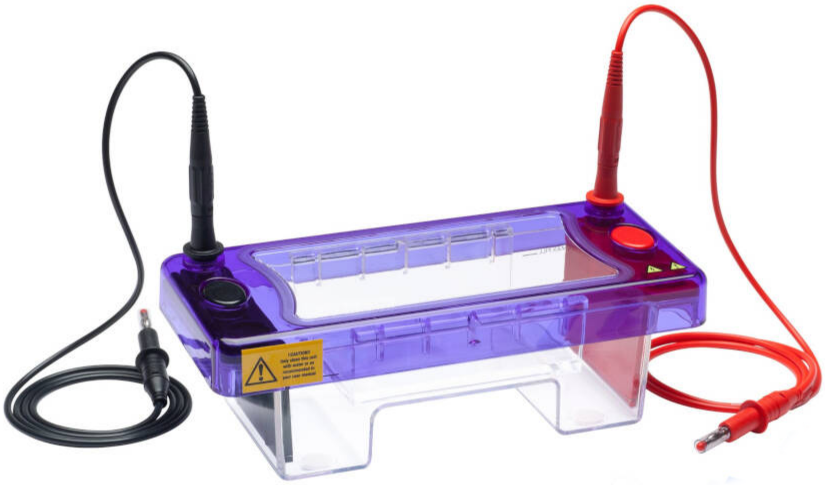

### Materials and methods

An important part of gel electrophoresis is the gel. We will be using a 1.5% agarose gel stained with GelRed® nucleic acid stain. All gels will be premade and there is one gel per group. Each gel has a 12-well comb.

#### Materials

In front of you you will find the following equipment:

- ☐ P1000, P200, P100, P20 and P10 Gilson pipettes and pipette tips
- ☐ 0.2 ml PCR strip tubes
- ☐ Gel electrophoresis tank
- ☐ Premade 1.5% agarose gel
- ☐ Sodium borate buffer
- ☐ Loading dye
- ☐ DNA ladder
- ☐ Microcentrifuge

You will also need your PCR products.

***NOTE:*** Check if everything required is present. If you cannot see or are missing any of the required materials, ask for assistance - we are here to help.

### Gel electrophoresis protocol

#### Preparing the PCR product

Adding loading dye to the PCR product makes it easier to see while loading, but also allows it to sink into the wells better.

It is a good idea to stick to the strip tube layout used in PCR preparation:

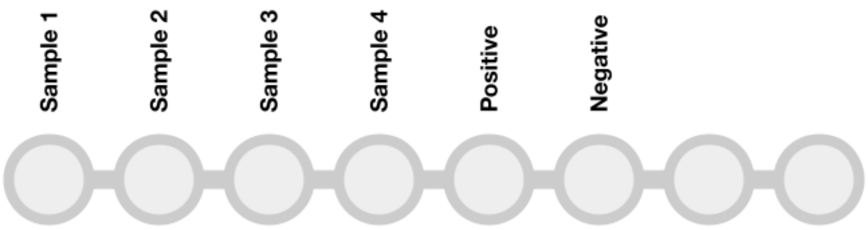


*Hint: again, pre labelling your tubes is a good idea*

1. Add 1 µl loading dye to to each tube in the PCR strip that is needed (samples plus controls)
2. Add 5 µl of each of your PCR products to the relevant PCR tube
3. Seal all caps securely
4. Gently flick PCR strip tubes to mix contents, spin down briefly

#### Preparing the gel tank

1. Taking care not to damage the gel, remove the sealing tape from both ends of the gel tray
2. Place the gel tray in the electrophoresis tank
3. Slowly fill the electrophoresis tank with sodium borate buffer until the gel is submerged
4. Carefully remove the comb from the gel

#### Loading the gel

---

***IMPORTANT:*** *While loading the gel try not to move or knock the gel tank. Do not pierce the gel with the pipette tip. Remember your pipetting exercises. Any issues please ask a demonstrator for help.*

---

1. Add 5 µl of DNA ladder to the first well
2. Add 5 µl of each of your PCR products to the adjacent wells:

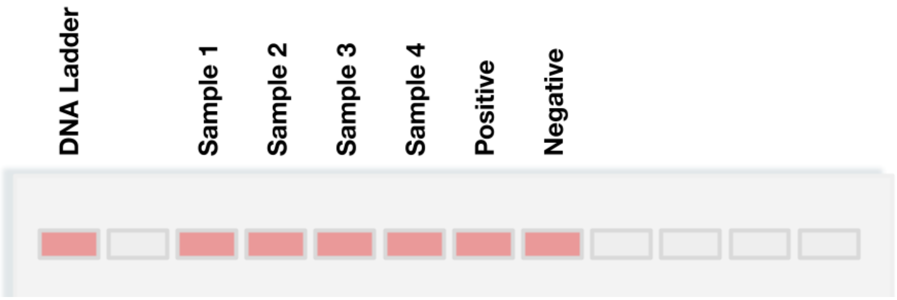


3. Replace the gel tank lid

Once completed let a demonstrator know. They will check your progress and show you the final steps to run the gel.

### A short time later…

Your gel will take 30 minutes to run, don't go anywhere though.

#### Removing the gel

Once complete, disconnect the power pack from the gel tank. Ask a demonstrator if unsure. Carefully remove the gel and tray from the gel tank and place it on a tray.

#### Imaging the gel

Find a demonstrator to show you the process of visualising the gel on the GelDoc imager.

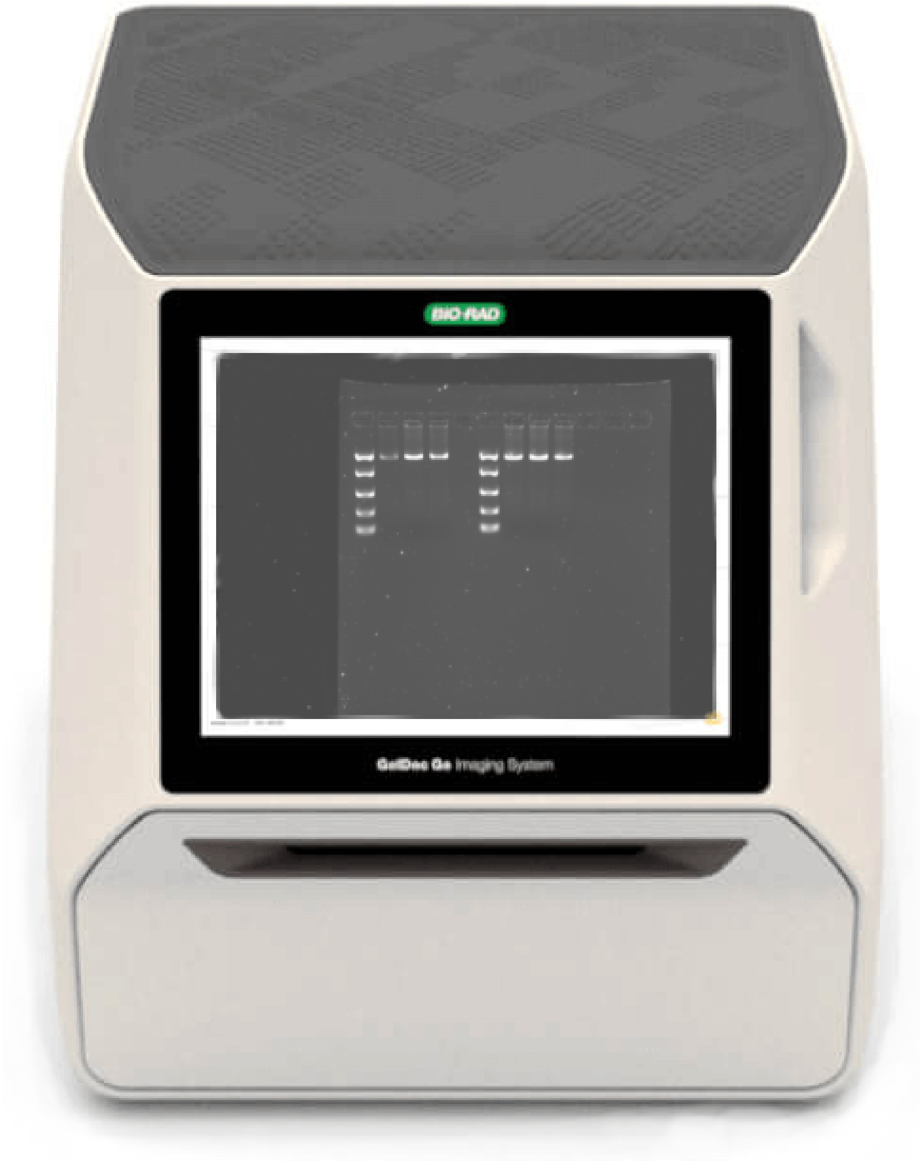

### Gel analysis

You should now see an image of your PCR product run on the gel on the screen of the GelDoc. Discuss the results with a demonstrator while at the GelDoc.

Importantly, has your PCR been successful?

If there were any issues, what could have been the cause?

You have used the EasyLadder I DNA ladder. The bands are of specific DNA fragment lengths. The PCR product is ~ 1,500 bp in length. Does your product appear to be around this length? Your gel should look similar to the image below.

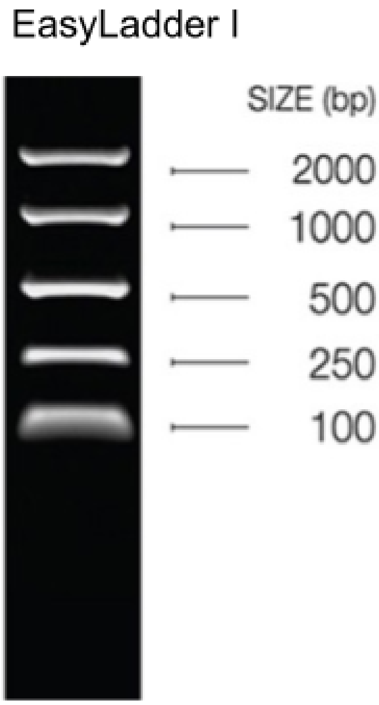


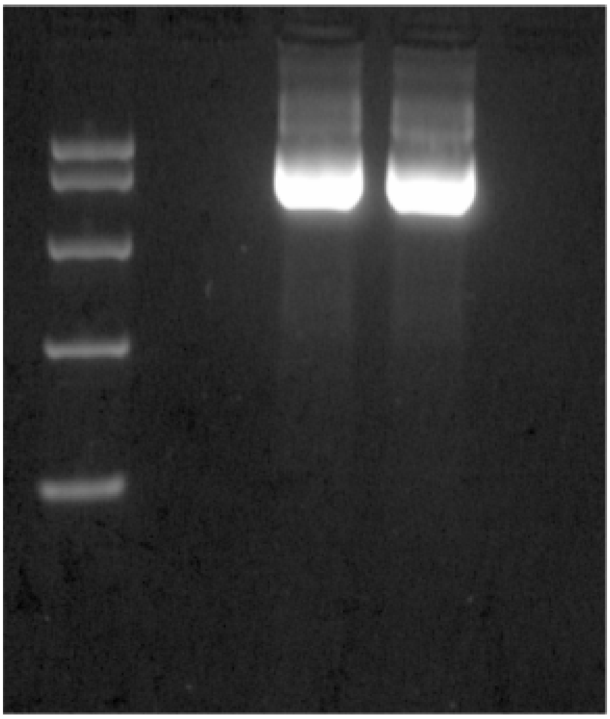

#### Gel images

Your images will be labelled by group number, saved to a USB drive and shared with you via Canvas. Ensure you have the order of samples recorded somewhere to allow you to label the image for future use.

### Day 3 skills

Today you have:

- Loaded and run an agarose gel
- Imaged and analysed the gel

More important laboratory skills for genetic analysis.

### Week 1 achievements so far:

You have now got the following Week 1 achievements:

1. Extract purified DNA from a bacterial culture ✓
2. Quantify the extracted DNA ✓
3. Run a PCR using bacterial 16S markers ✓
4. Analysed PCR results via gel electrophoresis and imaging ✓

WELL DONE!

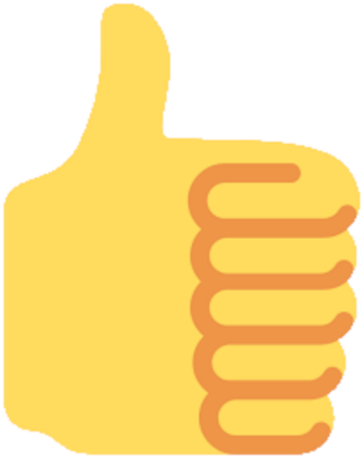

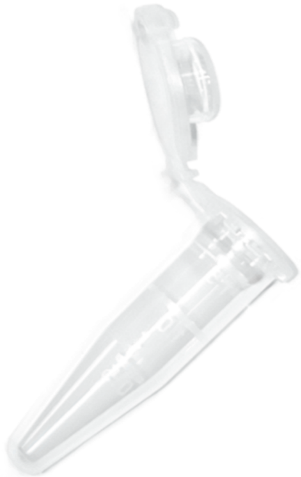

### Tomorrow

The Experiment. Oxford Nanopore DNA sequencing library preparation!

### Genetic Analysis day 4

#### Overview

| Time | Activity |
|---|---|
| 09:00 - 09:30 | Introduction. “The Experiment”.<br>*Overview of the day, health and safety in the lab.* |
| 09:30 - 10:00 | Sequencing library prep. pt1 (PCR).<br>*ONT library preparation PCR.* |
| 10:00 - 13:00 | PCR running on the thermo cycler<br>*2 hrs 30 mins to complete.*<br>*Brief talk and lunch break.* |
| 13:00 - 14:00 | Sequencing library prep. pt2 (gel and Qubit).<br>*Running and visualising a gel.*<br>*Qubit quantification of PCR product.* |
| 14:00 - 15:00 | Practical Week 1 roundup.<br>*Reflection on skills learnt so far.*<br>*ONT sequencing overview.* |

### The Experiment

Today, you will be using all the skills you have learnt so far this week to produce a genetic library for Oxford Nanopore Minion sequencing! The data generated from the sequencing run will be the basis for the bioinformatics analysis of next week's practical. In turn, the results generated from your bioinformatics will be used for the practical writeup assignment.

So, today is a very important day. We begin it with a PCR based Oxford Nanopore Technologies bacterial 16S rRNA DNA sequencing library preparation.

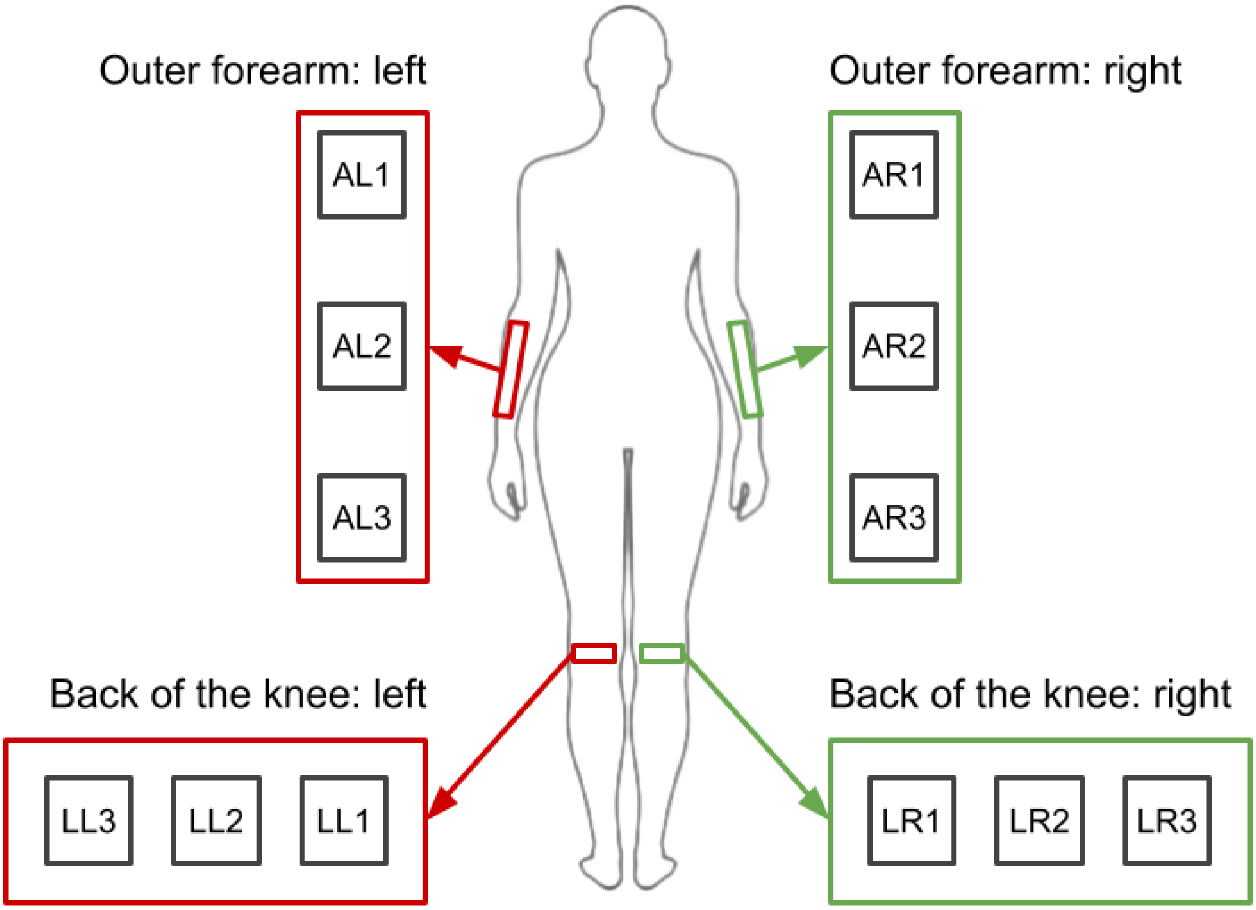


There are **12** human skin microbiome DNA extractions to be sequenced. Each group will be performing the library preparation for **2** of these **12** samples.

---

***IMPORTANT:*** *Sometimes you have a very limited starting material to use. This is one of those occasions. The DNA extracts for the samples are all there is. Each sample is 15 µl in total volume, of which 10 µl is needed for sequencing. This makes them a very precious commodity and they should be treated as such. This does mean there is no margin for error! This is a reality of genetics based lab work, and, hopefully, right now you are acutely aware of it.*

*However, if you do not feel comfortable with the added pressure of adding the samples during the PCR preparation, please ask a demonstrator to do it for you.*

---

### Library preparation

The library preparation will use Oxford Nanopore Technologies 16S Barcoding Kit (SQK-RAB204). It allows for sequencing of up to 12 samples on a single flow cell, a process called multiplexing. This is achieved by the primers having a unique “ID tag” in them, used to separate sequencing reads belonging to each sample.

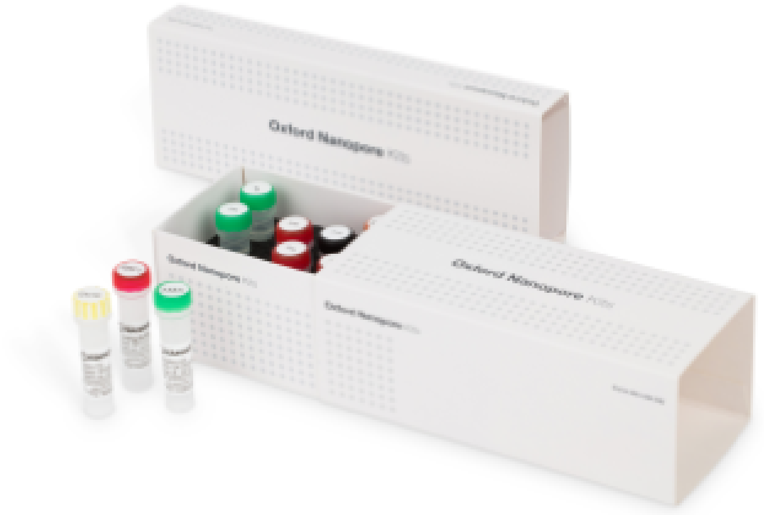

### Materials and methods

The Human microbiome DNA samples and 16S Barcoding Kit primers will be kept on ice at the front of the lab. Once you have finished the PCR preparation you will come to the front and add the relevant primers and DNA template to your PCR preparation.

#### Materials

In front of you you will find the following equipment:

- [ ] P1000, P200, P100, P20 and P10 Gilson pipettes and pipette tips
- [ ] 0.2 ml PCR strip tubes
- [ ] 1.5 ml tubes
- [ ] Ice bucket with PCR reagents
- [ ] Microcentrifuge
- [ ] Vortex mixer
- [ ] DNA ladder

PCR reagents in the ice bucket:

- [ ] Taq DNA Polymerase master mix
- [ ] Molecular grade $H_2O$

At the front of the lab, on ice:

- [ ] 16S Barcode Primer 1-12
- [ ] Human microbiome DNA samples

***NOTE:*** Check if everything required is present. If you cannot see or are missing any of the required materials, ask for assistance - we are here to help.

### Sample and primer sequencing layout

***IMPORTANT:*** Each sample requires its own uniquely tagged primer for sequencing. Refer to the table below for the required primer to be used per sample.

| Sample name | Primer ID |
|---|---|
| LL1 | 16S_01 |
| LL2 | 16S_02 |
| LL3 | 16S_03 |
| LR1 | 16S_04 |
| LR2 | 16S_05 |
| LR3 | 16S_06 |
| AL1 | 16S_07 |
| AL2 | 16S_08 |
| AL3 | 16S_09 |
| AR1 | 16S_10 |
| AR2 | 16S_11 |
| AR3 | 16S_12 |

### ONT 16S Barcoding Kit PCR preparation protocol

Unlike the previous PCR preparation you have performed, you will only need to prepare 2 reactions. As this is for pure library preparation, there will be no negative or positive controls. You only have 2 samples that will be using a specific “ID tagged” primer per reaction, so the need to make a PCR master mix is redundant. It is important to note that the volumes of the reactions are different to those used previously, this is due to the specific purpose of the PCR. Make sure you double check all measurements as you go.

***NOTE:*** Before you start, vortex Taq DNA Polymerase master mix and spin down briefly.

1. Add 25 µl Taq DNA Polymerase master mix to each tube in the PCR strip that is needed
2. Add 14 µl molecular grade $H_2O$ to each tube in the PCR strip that is needed
3. Ensure all caps on the PCR strip tube are closed securely
4. Gently flick PCR strip tubes to mix contents, spin down briefly, place on ice

**Final volume per tube** = 39 µl

Below is the final layout for your samples in the PCR strip tubes:

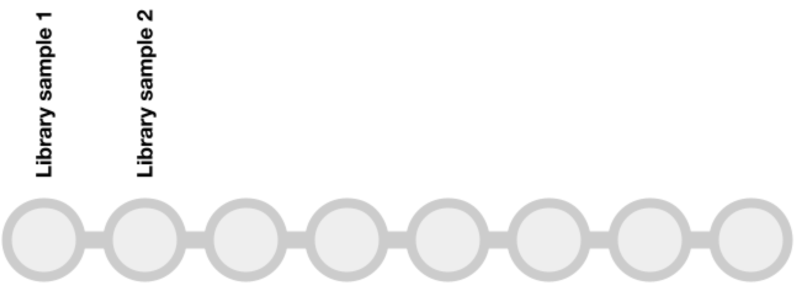


*Hint: pre labelling your tubes is a still a very,very good idea*

---

***IMPORTANT:*** *Ask a demonstrator to confirm you have correctly added the volumes required before continuing.*

---

Now you have prepared the PCR reaction tubes it is time to add the “ID tagged” primers and sample DNA template. Once you have reached this point, let a demonstrator know. When feasible (there may be a queue), you will go to the front of the lab to add in the “ID tagged” primers and sample DNA.

1. Add 1 µl of the relevant “ID tagged” primer to each of your PCR preparations
2. Add 10 µl of the relevant Human microbiome DNA sample to each of your PCR preparations
3. Ensure all caps on the PCR strip tube are closed securely
4. Gently flick PCR strip tubes to mix contents, spin down briefly, place on ice

Once completed your PCR preparations will be placed on ice until all groups have finished.

### Sequencing library preparation PCR

Once all preparations are done, the samples will be placed in the thermal cycler. It's going to take 2 hrs 30 mins for the PCR to be completed.

The thermal cycler conditions used for the library preparation PCR are:

***1 min @ 95°C, 40 X (20 secs @ 95°C, 30 secs @ 55°C, 2 mins @ 65°C), 7 mins @ 65°C.***

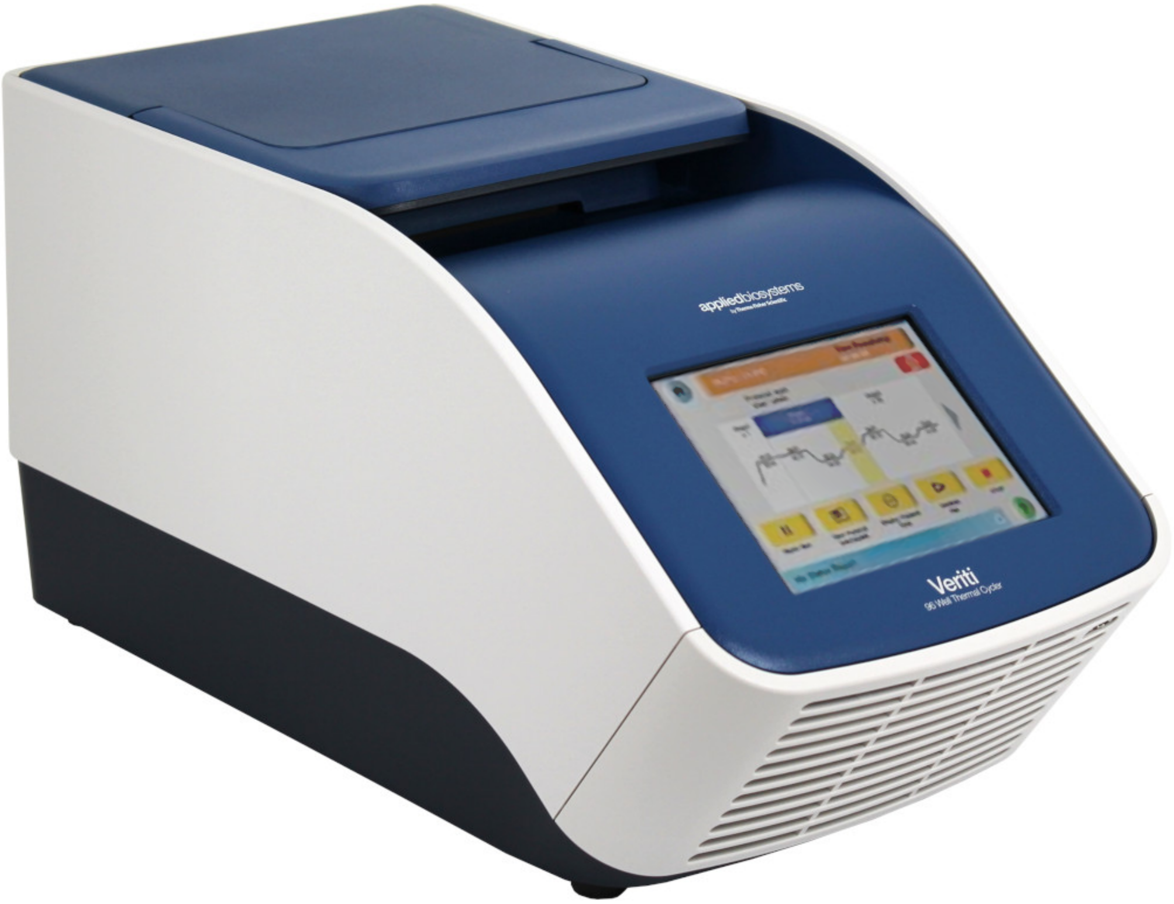


Hello again, Applied Biosystems™ Veriti™ 96-well thermal cycler, we have missed you.

***IMPORTANT PAUSE:*** *BREATH! The difficult bit is done. You have dealt with sensitive samples in an accuracy demanding protocol. WELL DONE!*

*The PCR will take a bit of time. Just chill, everything is cool. In the meantime, while the PCR is running, we shall have a thought provoking, informal talk. After which there will be a lunch break, I promise. We can't let you starve, can we?*

### Gel electrophoresis of sequencing library product

This is the same as running the gel from the other day. You will therefore be familiar with it. But, for completeness, the process is covered here again. Remember the important bits from your previous experience.

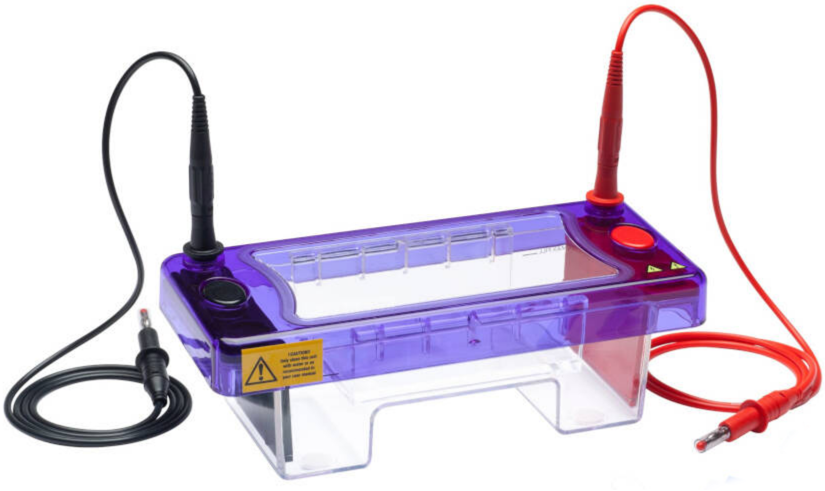

### Materials and methods

An important part of gel electrophoresis is the gel. We will be using a 1.5% agarose gel stained with GelRed® nucleic acid stain. All gels will be premade and there is one gel per group. Each gel has a 12-well comb.

#### Materials

In front of you you will find the following equipment:

- ☐ P1000, P200, P100, P20 and P10 Gilson pipettes and pipette tips
- ☐ 0.2 ml PCR strip tubes
- ☐ Gel electrophoresis tank
- ☐ Premade 1.5% agarose gel
- ☐ Sodium borate buffer
- ☐ Loading dye
- ☐ DNA ladder
- ☐ Microcentrifuge

You will also need your library PCR products.

***NOTE:*** Check if everything required is present. If you cannot see or are missing any of the required materials, ask for assistance - we are here to help.

### Gel electrophoresis protocol

#### Preparing the library PCR product

The simple strip tube sample layout below is recommended.

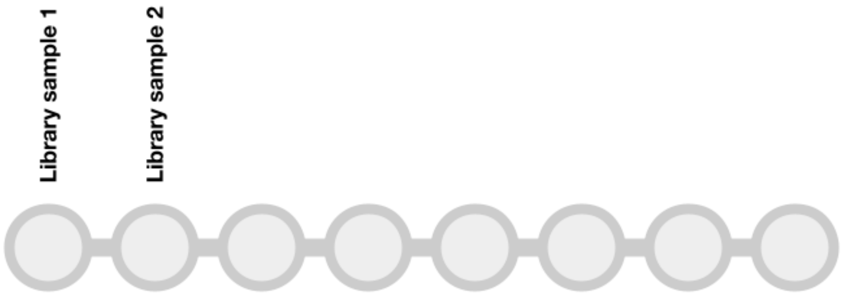


*Hint: again, pre labelling your tubes is still a very sound idea*

1. Add 1 µl loading dye to to each tube in the PCR strip that is needed (samples plus controls)
2. Add 5 µl of each of your library PCR products to the relevant PCR tube
3. Seal all caps securely
4. Gently flick PCR strip tubes to mix contents, spin down briefly

#### Preparing the gel tank

1. Taking care not to damage the gel, remove the sealing tape from both ends of the gel tray
2. Place the gel tray in the electrophoresis tank
3. Slowly fill the electrophoresis tank with sodium borate buffer until the gel is submerged
4. Carefully remove the comb from the gel

#### Loading the gel

1. Add 5 µl of DNA ladder to the first well
2. Add 5 µl of each of your PCR products to the adjacent wells:

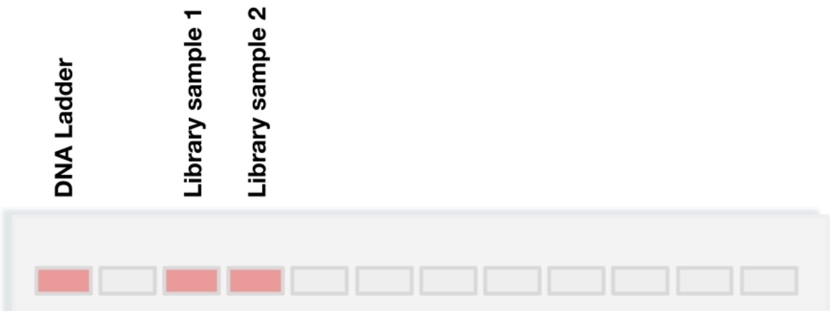


4. Replace the gel tank lid

Once completed let a demonstrator know. They will check your progress and remind you of the final steps to run the gel.

### A short time later…

Your gel will take 30 minutes to run. Take the time to make some notes.

#### Removing the gel

Once complete, disconnect the power pack from the gel tank. Ask a demonstrator if unsure. Carefully remove the gel and tray from the gel tank and place it on a tray.

#### Imaging the gel

Inform a demonstrator and image the gel on the GelDoc imager.

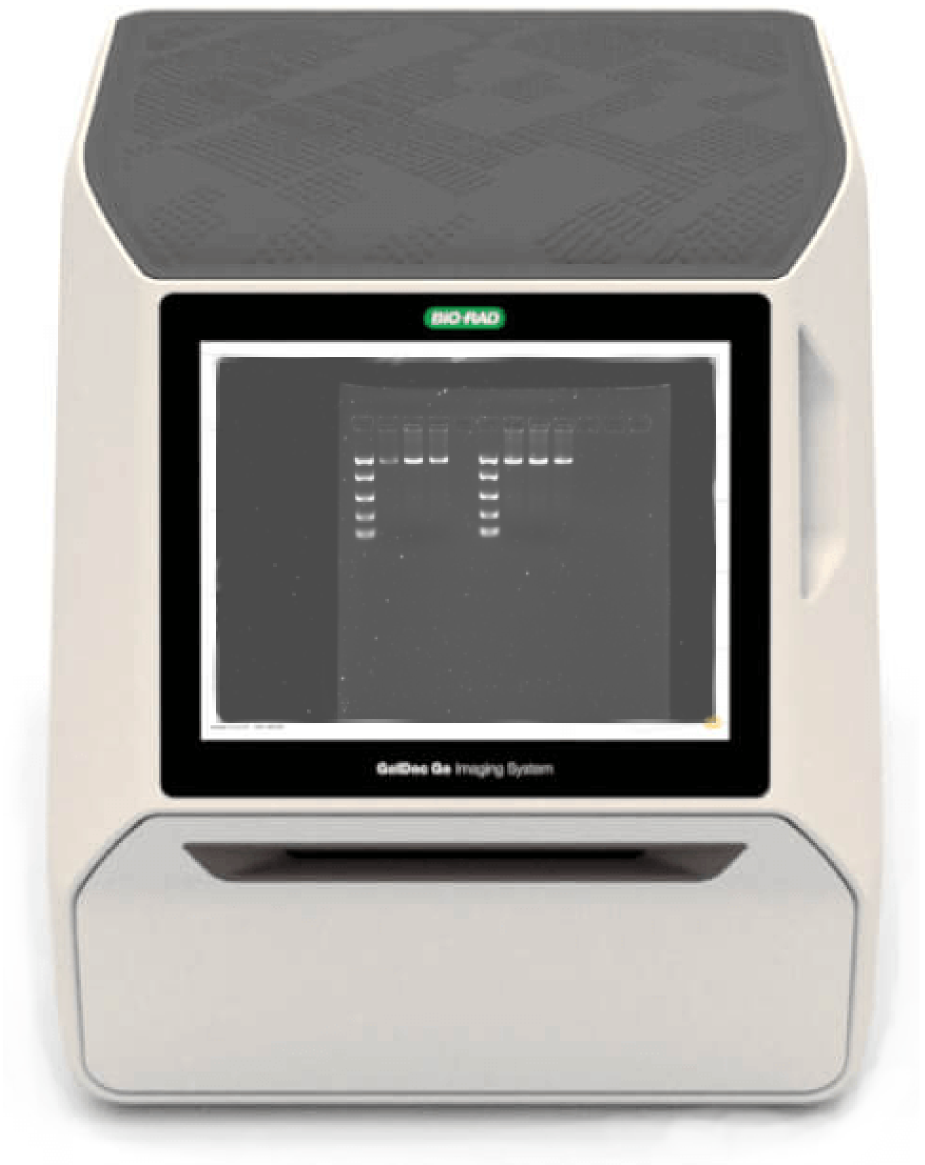


#### Gel images

Your images will be labelled and saved as before, then shared with you via Canvas. Ensure you have the sample names and primers recorded for future use.

### Assessing sequencing library quality

You have now prepared and visualised your sequencing library! Your final product length is ~1,500 bp. You have used EasyLadder I as your ladder, the same as the last time. You should now be familiar with what to expect and how to interpret your results. Your gel should look similar to the image below.

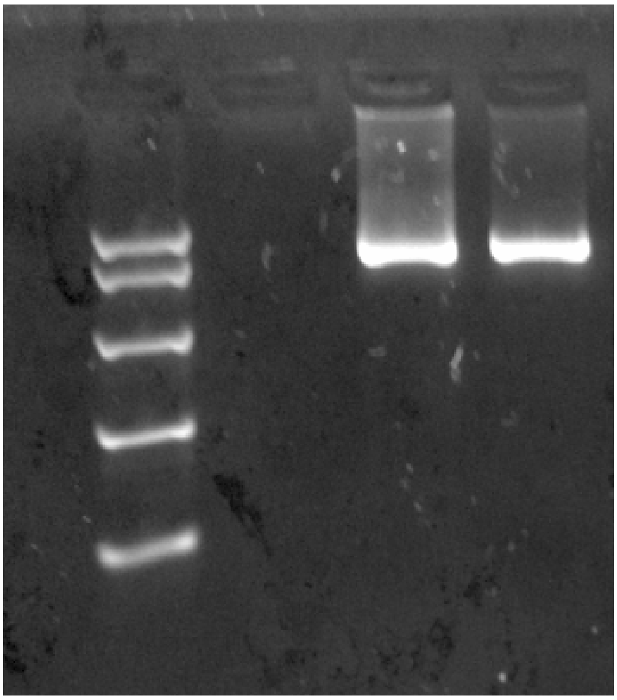

#### Important points to consider

The following questions are key to assessing the quality of your library preparation:

**Has the gel run properly?**

**Are the results clear enough to interpret correctly?**

**Are the PCR products of the samples the expected length?**

**Are all the samples of equal brightness?**

**Are there unexpected bands present in the samples?**

**Can you think of any more questions?**

Discuss your results with a demonstrator. Once you are sure of the results and have interpreted them correctly, let's move on to some calculations.

### Sequencing library quantification

Now we have a library preparation, we need to determine the amount of DNA within the library PCR product. We will use the Qubit fluorometer for this, like before. From the results we can determine the molarity of the samples to be sequenced.

### Materials and methods

As before, all the samples will be measured in a single batch using the same controls. This will make all the results directly comparable. The Qubit buffer master mix and controls required for the process are all pre-prepared at the front of the lab.

For a reminder, here are the volumes required for the Qubit buffer and Qubit dye for a single sample.

Add 1 µl of Qubit dye to 199 µl Qubit buffer. Vortex to mix, spin down briefly.

You will quantify 2 µl of each of your sequencing library products.

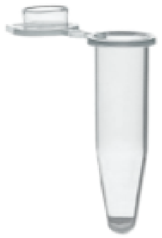

**Qubit tube**

#### Materials

In front of you you will find the following equipment:

- ☐ P1000, P200, P100, P20 and P10 Gilson pipettes and pipette tips
- ☐ Qubit tubes
- ☐ Microcentrifuge
- ☐ Vortex mixer

### DNA quantification protocol

1. Vortex DNA sample to mix thoroughly, spin down briefly
2. Transfer 198 µl Qubit buffer master mix to a Qubit tube
3. Add 2 µl of sequencing library PCR product. Vortex to mix, spin down briefly
4. Incubate at room temperature for 2 mins

Once your sample is ready, ask a demonstrator and they will show you (again) how to quantify it on the Qubit. Make a note of your sample's DNA concentration - this time it's important.

### Molarity calculations

Prior to final library preparation, all samples must be pooled together at roughly equal molarity. This means that each sample will have around the same number of DNA copies to be sequenced, therefore each sample is equally represented in the sequencing output.

To calculate the molarity of your sample you will need to know:

1. The DNA fragment length
2. The DNA concentration

As luck would have it, you already have these values.

#### Calculators at the ready

Using the equation below, calculate the molarity for each of your samples.

$$\text{fmol}/\mu\text{l} = \text{DNA concentration (ng}/\mu\text{l)} / ((\text{length of DNA (bp)} \times 617.96) + 36.04) \times 1{,}000{,}000$$

***NOTE:*** The result will be given in femtomoles per microlitre (*fmol/*µl). A femtomole is $10^{-15}$ moles, so a very, very small measurement.

### Questions

**Are the samples of similar molarity?**

**How would you get them to be the same molarity?**

Discuss your findings with a demonstrator.

#### Bonus: calculating DNA copy number

You can use the fmol/µl values from above calculate the number of DNA copies in your samples:

$$\text{DNA copy number} = \text{fmol}/\mu\text{l} \times 6.022 \times 10^{8}$$

---

***FINAL STAGES OF LIBRARY PREPARATION AND SEQUENCING:*** *Now we know the molarity of your samples, the final stages of library preparation can be performed. This will be done by me as it is a single person job and can be very tedious and demanding.*

*You have all done a brilliant job this week and deserve a break.*

---

### Day 4 skills

Today you have:

- Performed ONT library preparation
- Assessed sequencing library quality

More complex but no less important laboratory skills.

### Week 1 achievements so far:

You have now got the following Week 1 achievements:

1. Extract purified DNA from a bacterial culture ✓
2. Quantify the extracted DNA ✓
3. Run a PCR using bacterial 16S markers ✓
4. Analysed PCR results via gel electrophoresis and imaging ✓
5. Use the above skills to prepare and quantify a genetic library for MinION sequencing ✓

WELL DONE!

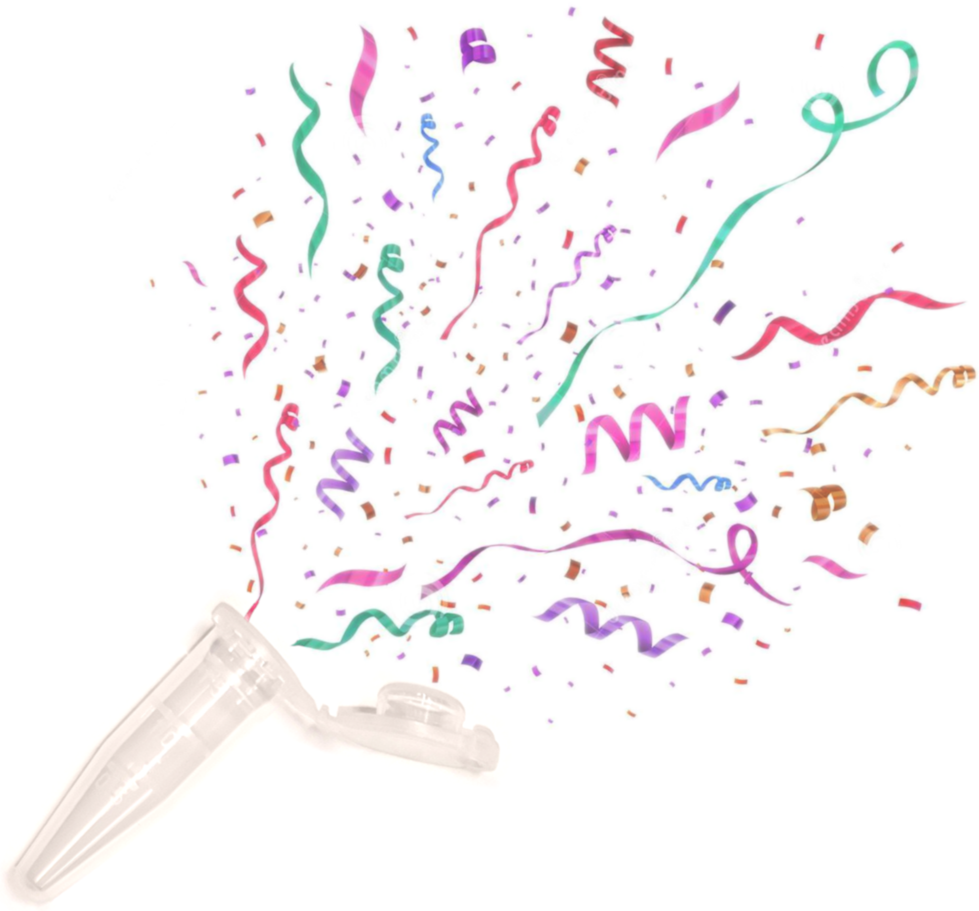

### Next week

Bioinformatics: computational analysis of sequencing data. See you then!

Blank for notes.
