## Supplementary 2: Bioinformatics manuals for "Who Grows There? A Course-based Undergraduate Research Experience to explore the human microbiome through 16S DNA metabarcoding"

### GENETIC ANALYSIS: INTRODUCTION TO THE UNIX COMMAND LINE

#### Overview

This practical will introduce you to the unix command line and some of its basic utilities.
At the end of this practical session you will have experienced, and hopefully appreciate, the power and functionality of using the command line within a unix terminal and understand its wider applications to data management.

#### 1. Introduction to the command line

Below we will give only a very simple overview of the basic utilities of the unix command line. The best way to learn the command line is to Google search for instructions and then try it yourself, experimenting as much as you can.

For this practical, we have embedded the unix command line into a Jupyter notebook so that you have both the instructions and the command line exercises together. A normal command line terminal would be a seperate window, but otherwise it is the same and all the commands you will use in this session are applicable.

In the instructions below we denote commands typed at the terminal with a `$` sign, whereas the output, the info returned by the terminal, has no dollar sign.

```
$ a typed command
output
```

**Important:** When you type out code presented like this, do not include the `$` sign. It will not work and return an error!

The actual terminal where commands are written and run for this practical are the grey cells (boxes) prefixed by [ ]:
Here is one:

In [ ]:
```

```

You can run commands in a terminal cell by clicking in it, typing a command, and then Shift-Enter to run the command. You can also click the triangular `Run` button in the menu bar. Try it below, the 'hello world' `echo` command has been entered for you.

In [ ]:
```
echo 'Hello World!'
```

You should see that the output 'Hello World!' has been printed direcly below the cell. We haven't introduced the `echo` command, but you can probably guess what it does. If not google it.

### 2. Navigating the file system

---

The first thing to do is to learn the basics of moving between places on your computer, checking where you are, checking what files are present and having a quick look at them. Some of the terminology is perhaps slightly new ('directories' rather than 'folders') but using the right words will mean you are speaking the same language as everyone else and make your life easier when Googling for solutions.

#### 2.1 Checking where you are ( `pwd` ), listing contents ( `ls` ) and changing directories ( `cd` )

##### 2.1.1 Where am I? `pwd`

At the command line you need to know "where you are" i.e. which directory (folder) you have open and are working in, called the working directory. The command to find out where you are is `pwd` short for 'print working directory'. NB: 'print' when working at the command line means 'display on the screen' rather than 'write this to a piece of paper'.

Type the command to print working directory `pwd` below, and hit Shift-Enter to run the command.

In [ ]:
```

```

You should see something like this:

```
/home/bs/123456/genetic-analysis-bioinformatics-
practical/01_introduction_to_unix_command_line
```

**Tip: if you want to try something and there isn't an empty terminal cell, use the menu bar `+` to "Insert Cell"**

Try inserting a cell below and check your working directory again:

This is your **file path** which very specifically records the exact location of the file on your system. Unfortunately it's a bit long for this notebook, so I am going to ignore everything before `01_introduction_to_unix_command_line`

It would be useful to know what folders and files are present in your current location.

##### 2.1.2 Listing the contents of a directory with `ls`

A very common thing you will want to do is to display the contents of a directory, i.e. list all the files.

You can list the files (and directories) in your working directory using the `ls` command. Try `ls` in the cell below:

In [ ]:

```

```

You will see the files and directories in your current folder displayed by `ls`.

The jupyter notebook you are using will have the extension ".ipnb". There some directories: **data** and **images**.

##### 2.1.3 Changing directory with `cd`

But maybe this isn’t where you want to be, maybe you need to have a look at your new data, in which case you need to `‘change directory’` and the command for that is `cd`.

```
$ cd data
$ cd my_sequences
$ pwd
```

```
/01_introduction_to_unix_command_line/data/my_sequences
```

**Try navigating down to the `my_sequences` folder:**
Remember you can add extra cells, and use `pwd` and `ls` to help you.

In [ ]:

```

```

You can see that the `/` symbol denotes levels of directories, so that the ‘my_sequences’ directory is contained within the ‘data’ directory which is within the user's 'home' directory, and ultimately in the system’s ‘home’ directory.

These **file paths** can sometimes be long but they are always explicit, which is a very good thing for reproducibility. There is no excuse for failing to remember where the data was stored for your analysis, here it is written out, and in the real world you will want to record this as part of your experiment.

You can go up one level (to '**data**') by using double dots (ensure there is always a space between the `cd` command and the directory you wish to go to).

```
$ cd ..
$ pwd
```

/01_introduction_to_unix_command_line/data/

**Try it for yourself:**

In [ ]:

```

```

What happens if you use the `cd` command without telling the system where you would like to change directories to?
Try it. How can you find out which directory you are now in?

In [ ]:

```

```

Using `cd` command on its own returns you to your **user's home directory**, in this case `/username` from wherever you are. NB it is **your** home directory. **your home directory** is an important concept. In the example below *username* means your username, whatever it is.

The tilde symbol (`~`) is shorthand for your home directory so `username/01_introduction_to_unix_command_line` and `~/01_introduction_to_unix_command_line` refer to the same directory, which saves a little typing.

Another very useful shortcut is `cd -` (dash) which takes you to the previous directory that you were in. This is really useful when you need to swap between directories that are separated by several levels or that have long names.

**Try it for yourself:**

In [ ]:

```

```

Using the `cd` command navigate to `test_files` directory, examine the contents of the directory, then navigate back to the top `01_introduction_to_unix_command_line`

#### 2.1.4 tab completion saves time and errors

You are probably slightly annoyed now that you have to type in all these directory names, which are long and complex. If you make one typo it will result in an error. Real bioinformaticians make use of the tab key a lot. tab will suggest completions of your file path.

If I am in the home directory and I type `$ cd da<tab>` it will autocomplete to `$ cd data`.

Because of the way this Jupyter notebook is set up it may instead give you a list of options, you can navigate using the arrow keys, and hit RETURN when you have the right option.

This saves a LOT of typing and prevents errors.

**Insert a cell below and try it now:**

In [ ]:

##### 2.1.5 Chaining commands together with a semicolon

Above we we're issuing commands one at a time; first `cd` then `pwd`. To chain commands together on the same line separate them with a semicolon ie `cd;pwd`.

Try moving around some directories, printing where you are and listing the contents.

Do this by chaining together some `cd`, `pwd` and `ls` commands.

Are you tab completing the addresses?

In [ ]:

#### 2.2 Making directories and getting organised

I hope you see that we have begun to organise our working directory into sections. For example, we have a `/data` directory, with all our fasta data files in sub-directories.

Organising your bioinformatics experiment is very important, and its easy to get to a chaotic confused state if files are badly named and not in meaningful directories.

It is bioinformatics best practice never to modify files in your `data` directory, instead, if they are to be modified they are saved as a results, ie they have been successfully modified. This means we need to know how to create a `/results` directory.

##### 2.2.1 `mkdir`

The command to create a directory is `mkdir`

In order to use this sensibly it will require you to (a) state the name of the directory to create and (b) say where you would like it created (default is your current working directory). These are normally combined.

Move to the top level `01_introduction...` directory

Create a directory called "test-dir" in `data/test_files`

`mkdir location/dir-name`

Now create a `results` directory at the same level as your `/data` directory

#### 2.3 Summary

```
pwd     print working directory, show where you are
ls      list, show all the files in your current directory
cd      change directory, move to another location
mkdir   make a directory
;       semicolons can chain together commands on one line
```

Test yourself, demonstrate that you can do something using each of these

In [ ]:

```

```

### 3 Inspecting file contents, and searching within text files

#### 3.0 fasta refresher

You may have come across fasta sequence files before, you can Google them, remember Googling is a bioinformatics skills not a cheat. Fasta files contain fasta sequence records, and are a very simple and human-readable way to display sequence data. Each sequence in a fasta file has 1 line of header (a title) beginning with the greater than sign (>), then, on the next line the sequence itself.

Fasta files may contain 1 or very many sequences. Typical file names end in `.fasta` or `.fas` and contents may look like this:

```
>apple DNA sequence 1
ACGCATGCACGTACGATCGATCGATCGATCGACTGATCGCTAGCTAGCTA
>banana DNA sequence 1
TTGCATGCACGTACGATCGATCGATCGATCGACTGATCGCTAGCTAGCTA
>cherry DNA sequence 1
GGGCATGCACGTACGATCGATCGATCGATCGACTGATCGCTAGCTAGCTA
```

#### 3.1 Inspecting files

Much of bioinformatics is the inspection and retrieval of information from text files. This data is the source of all subsequent analyses and we need to make sure it is the right data in the right format.

If we want to inspect a file we can display its contents using several unix programs:

```
cat     to print the whole file to the screen
head    to print the first few lines of the file on screen
tail    to print the last few lines of the file on screen
```

One of the problems with DNA sequence files is that they can be large - several hundred megabytes to a few gigabytes is not uncommon. Viewing these files can be difficult, as the files need to be loaded into memory, and can therefore take a great deal of time for the text editor to read from the disk. The `head` and `tail` commands are very efficient for viewing large files such as these.

**Remember Googling things is encouraged**, it is a great way to learn bioinformatics! Google `unix head command` and `unix tail command` to learn more about them if you wish. Both `head` and `tail` display 10 lines by defualt, but they allow you to specify the number of lines to display, so `head -20` would display the first 20 lines and `tail -5` the last 5 lines.

Remembering tab completion, navigate to the directory `/data/my_sequences`.
Tip; when navigating use `ls` to see what is available.

Use `head`, `tail` and `cat` on the files in `data/test_files`

**Try it now:**

In [ ]:

```
```

The files you have inspected are small and easily readable with the `head`, `tail` and `cat` commands.
Some files can be extremely large and using `cat` may take a long time or even crash the system - not good!

If you are unsure about the size of a file `head` is a good way to have a look at it.

However, if you actually want to know how big a file is you can display this with a version of the `ls` command `ls -lh`. It can be used on files or directories, it is a very useful command.

**Try it:**

In [ ]:

```
```

All the commands above will be very useful for basic bioinformatics work, they are key skills.

#### 3.2 Searching within files

Searching within very large files however can also be problematic, especially using the standard 'find' functions in a text editor, which aren’t optimised for performing searches across very large files. For this reason various tools have been created that allow users to search within large files from the command line, and are highly optimised for their function. One of the most useful utilities for searching within a file is `grep` (global regular expression parser).

`grep` is very simple to use. At the command line you will need to type the word `grep`, followed by the text you are searching for, followed by where (the filename) to look for it. For example, to search for the word 'Bos' in our `UK_vertebrates.fasta` file we do the following:

```
$ cd data/my_sequences/
$ grep 'Bos' UK_vertebrates.fasta
>AB074962.1 9913 Bos taurus
```

This returns the text `>AB074962.1 9913 Bos taurus` which is the single line that contains the word Bos. This is the output of `grep`, the line on which the search phrase is found.

Make sure that you are trying this for yourself, by inserting cells and giving the command

We can confirm that there is only 1 instance by using the **count flag** (`-c`) with `grep` as follows:

```
$ grep -c 'Bos' UK_vertebrates.fasta
1
```

(Flags are useful things, they allow the default behaviour of a command to be refined. You have already used one (-lh) with the `ls` command to make it show you files sizes.)

#### 3.2.1 How many sequences do I have?

A very common question is to ask is 'how many DNA sequence records are in this fasta file?'

A fasta format file shows a sequence over two lines. The first line is the sequence name, the second is the actual DNA sequence.

For example:

```
>ACC01.1 a_sequence
CGACCTTATTATCAGCTTTAACTCAA
```

Remember the sequence name always starts with `>`.

**Now try to see if the sequence file 'UK_vertebrates.fasta' matches your expectations. You already have the skills to do this without loading the whole file into memory, can you remember how? Check above if not.**

In order to find out the number of sequences you could of course search for all the greater than `>` symbols, which is almost certainly the number of records. However, you should really search for all the lines *starting with* `>` rather than the number of times it occurs, as it is possible for a fasta header to contain an internal `>`. 'Line starts with' is represented by the `^` symbol.

Design a `grep` search to count the number of fasta header lines. Do not copy/paste from any web resource. Discuss your solution with the person next to you or check with a demonstrator.

**Here's a cell to try it in:**

In [ ]:

```

```

#### 3.3 Summary

```
head    show the first few lines of a file
tail    show the last few lines of a file
cat     print the whole file to the screen
grep    search, the -c flag counts the matches
```

#### -- PAUSE --

You have just learnt some basic unix command line utilities and programs. Well done!

There are still many, many more things that can be done that we have not covered in this session.

For example:

- using wildcards
- copying and moving files
- writing to a file
- appending to a file

Learning how to do them is as simple as Googling them.

##### Googling is not just OK, it is encouraged!

The internet is a source of knowledge. It is mined by those eager to learn.
It is how all bioinformaticians develop and progress.
There is no simple handbook, there are so many different ways to achieve the same result.

### Outcomes of this Introduction

You have now completed a brief introduction to the unix command line. This is the foundation of bioinformatics and you will use thise skills duering the analysis of your 16S data later in the practical.

You have learned to navigate the file system, examine and search DNA sequence files, and start to get organised with big data.

The reason we use the command line is because of the scale of data and the need for reproducibility. These just could not be achieved with normal graphical user interfaces.

### 4. Future reference

**Important:** Before you leave this jupyter notebook, use the internet browser menu to `print as PDF`. Ask a demonstrator if you need help.

This can then be kept as a reference for you in the future. Something to look back on.

### End of session

**You have now learnt sufficient command line skills for the next session:**

#### ONT sequencing data analysis

### GENETIC ANALYSIS: ONT SEQUENCING DATA ANALYSIS

#### Overview

This practical will introduce you to the bioinformatic analysis pipeline used to generate results from ONT MinION sequencing data.
At the end of this practical session you will have bioinformatically analysed a small test set of sequencing data.
Importantly, you will have experienced the steps used in this kind of analysis.

#### 1. Introduction to the bioinformatic analysis workflow

This section of the practical assumes the completion of the previous section: **Introduction to the Unix command line**.
You will use your newly aquired skills to perform the exploration of the sequencing data and execute the analysis in this section.

##### 1.1. What has happened so far

Last week we created a final baterial 16S rRNA library for sequencing on Oxford Nanopore Technologies' MinION.
The final stages of library preparation have been performed. The prepared DNA sequencing libaray has been loaded onto the MinION and sequenced. The sequencing output has been "basecalled" and we have data in a meaningful file format ready for us to analyse bioinformatically.

##### 1.2. What happens now?

**We can at last analyse the data!**

This is done in 3 stages:

1. Quality control
2. Taxonomic assignment
3. Format outputs

##### 1.3. Workflow programs

As a breif outline, we will use 3 specific command line based programs for data anlysis.
These are all pre-installed and ready to use, so do not worry.

1. `seqkit` is used to look at the stats of the data (read length, read count etc.).
2. `fastp` performs quality control.
3. `kraken2` performs taxonomic assignment.

We will use a custom `python` script to format the outputs.

### 2. Organising the working directory

Before we get started we have to make this look and perform more like a bioinformatic analysis working directory.
We need to create some directories for each stage of the anlysis to output results to.
To do this we will use a new command line utility `mkdir`.

`mkdir` creates a directory where you specify, or in your working directory ( `pwd` ).

#### 2.1. Create a *results* directory

We need to create a **results** directory where all the outputs from the analysis go.
Use the following command:

```
$ mkdir results
```

**Do it now:**

In [ ]:

```

```

#### 2.2. `fastp` and `Kraken2` directories

In the **results** directory we need to create **qc** (quality control) and **kraken2** (taxonominc assignment) directories.

These are for `fastp` and `kraken2` to output to respectively.
Use the following command to make the **qc** directory:

```
$ mkdir results/qc
```

**Note:** you did not have to change directory here to do it. The path to the directory is sufficient. Another unix command line skill learned :)

**Run the required commands to create both the qc and kraken2 directories:**

In [ ]:

You should now have two directories: **qc** and **kraken2**, in your **results** directory. Use your command line skills to check:
*hint* `ls`

In [ ]:

##### Important: Before continuing, check with a demonstrator that you have done this all correctly!

### 3. The bioinformatic analysis workflow

#### 3.1. The fastq format

The data we have is in "fastq" format. This is similar to "fasta" format, but another way to store genetic data.

However, fastq files hold more information - specifically for DNA sequencing quality.

A fastq file has 4 lines per sequence:

1. Sequence name
2. DNA sequence
3. Spacer ("+")
4. Quality score

For example:

```
@sequence_name
GCGAACTTTGCTAGCGGCAAGGCGCTTACAGCAAGTCGAGCGAGGAC
+
%$%%$$$&''&&'((),,*%$%$#$%'''&&&'(((((-/////)((
```

For quality control the quality of each base is an important factor.
Base quality is given as a "Q-Score": a character-based code for the quality of the base.
Go to this link to see the actual values (look at the "Symbol" and "Q-Score" columns):

https://support.illumina.com/help/BaseSpace_OLH_009008/Content/Source/Informatics/

Now you understand the format of the data you need to analyse, use your command line skills to look at the first 4 lines of the `test_sample.fastq` in the `/data` directory:

In [ ]:

Does this look like the format you expected?

Discuss with the person next to you and check with a demonstrator.

### 3.2. Quality control of fastq data

We have the raw reads from the MinION sequencer but we need to know which reads to keep.
Which are of good enough quality to consider for further analysis?

We will use `fastp` to do this. We will also utilise `seqkit` to look at before and after quality control.

#### 3.2.1. `SeqKit` to inspect raw fastq data

Let's look at the vital statistics of the "test_sample.fastq" file using `seqkit` prior to quality control:

In [ ]:
```
seqkit stats data/test_sample.fastq
```

Can you describe exactly what the output is showing? Check with a demonstrator if you are unsure.

#### 3.2.2. `fastp` qc of raw sequencing data

OK, lets quality control the fastq data using `fastp`:

In [ ]:
```
fastp \
    --disable_adapter_trimming \
    --in1 data/test_sample.fastq \
    --out1 results/qc/test_sample.qc.fastq \
    -j results/qc/test_sample.json \
    -h results/qc/test_sample.html \
    --qualified_quality_phred 7 \
    --unqualified_percent_limit 40 \
    --average_qual 7 \
    --length_required 1000 \
    --max_len1 1700
```

There is a lot going on here. **Don't panic!**

`fastp` has a lot of functions and subsequently a lot of "flags".
It is a very powerful tool and well worth knowing how to use for quality control of sequencoing data!

You could refer to the `fastp` docs to see what was just done, but this is optional, you can understand enough from the instructions here:

See if you can understand what the command above did. Check with a demonstrator if unsure.

##### 3.2.3. `SeqKit` to inspect quality controlled fastq data

Let's look at the vital statistics of the quality controlled fastq data we have just generated.
It is now in the **results/qc** directory as "test_sample.qc.fastq".

Shall we see what quality control did? `seqkit` to the rescue:

In [ ]:
```
seqkit stats results/qc/test_sample.qc.fastq
```

Compare the post quality control `seqkit` output to that of the `seqkit` output for the raw sequencing data.

Does this make sense as to what `fastp` has done?

Discuss with the person next to you or check with a demonstrator if unsure.

**TIP**

You can specifify 2 files for `seqkit stats` to display just by listing their paths.
Then you will get both files displayed in a single table, whioch will be much easier to compare.

`seqkit stats path/to/file1 path/to/file2`

Try it to get a much more informative table

In [ ]:
```

```

#### 3.3. Taxonomic assignment with `Kraken2`

The raw reads have been quality controlled with `fastp`. They are now considered of sufficient quality for taxonomic assignment.

`kraken2` is an ideal program to use for analysisng Oxford Nanopore Technologies' MinION sequencing data. It is a "kmer" based taxonomic assignment method. It is again an incredibly powerful tool worth knowing how to use correctly.

In your spare time, and if you are interested, read the supplied literature in the Genetic Analysis module's "to read" section for an insight as to what `kraken2` does. Google for further examples of kmer based methods to get a better overview of the approach.

##### 3.3.1. `Kraken2` database

`kraken2` requires a database with which to perform it's analysis. This is supplied.
The database *is* the directory **bact_db**: a `kraken2` database specifically created for this example anlysis.

##### 3.3.2. `Kraken2` taxonomic assignment

Now we have a database and some quality controlled fastq data.

In [ ]:
```
kraken2 \
    results/qc/test_sample.qc.fastq \
    --db bact_db \
    --report results/kraken2/test_sample.txt \
    --output results/kraken2/test_sample.krk \
    --minimum-base-quality 7 \
    --confidence 0.02
```

Again, here is a lot going on here. **Don't panic!**

The important bit is that it has done it's job. the terminal output should have shown something happen and a lot of text will have been produced. Have a look at this and see if you can make sense of it.

Check with a demonstrator if unsure.

The **results/kraken2/** directory should now have files in.

**Explore them using you command line skills:**

In [ ]:
```

```

#### 3.4. Formatting the outputs

`kraken2` has given us outputs but they are not easilly readable, nor are they ideal for downstream analysis with *R*, for example.

The next step is to modify these outputs to be in a format for our use.

To do this we will use a custom `python` script to transform the output. `python` is it's own coding language, but we have a script (regardless of coding language) that forms an essential urpose. We shall now use it.

**Important - you do not need to know python to run this script, it is just a program, like seqkit is a program, and you can run it without looking at the code**

In the cell below we call the interpretter `python`, the path to the script and the required variables: `-i` and `-o` (input and output).

Run the command below to execute the `python` script:

In [ ]:
```
python scripts/to_tsv.py -i results/kraken2/test_sample.txt -o results/te
```

Can you determine what it has done from the command?

For those who are really interested, look at the script itself, it is located in the **scripts** directory.

#### 3.5. The final output

We have now generated an output that is suitable for our requirements.

Here we will use the `cat` command to view the whole file **results/test_sample.tsv**:

In [ ]:
```
cat results/test_sample.tsv
```

Is this a more approachable format for downstream analysis?

Discuss with the person next to you or with a demonstrator.

#### -- OUTCOMES --

**WELL DONE**

You have just completed many of the key analyses for taxonomic investigation of a bacterial community 16S data.

- You have used `fastp` to quality control data
- You have characterised your sequencing data with `seqkit` and generated data on how it changed through the workflow
- You have run the workflow for taxonomic assignment of sequencing data with `Kraken2``
- Using the command line you have explored the data at different stages of analysis.

### 4. Future reference

**Important:** Before you leave this jupyter notebook, use the internet browser menu to `print as PDF`. Ask a demonstrator if you need help.

This can then be kept as a reference for you in the future. Something to look back on.

### End of session

**You are now ready for the next session:**

#### `R` analysis of sequencing data

### GENETIC ANALYSIS: *R* ANALYSIS OF SEQUENCING DATA

#### 1. Data

*R* is widely used for data analysis. So it is no surprise that we must know how to deal with data when using it.

**Important**: We will use a test dataset initially. It is DNA metabarcoding sequencing data in the same data output format as you will have for your sequencing data. Columns are samples, rows are "species". Each entry in the dataframe is the number of sequenced reads per "species".

##### 1.1. Reading data into R

Data can be read into *R* in different ways. Here we will use the normal method for the kind of input data we have.
Our data is a tab-delimited (.tsv) output from the taxonomic assignment of the sequencing data generated from the lab practical.

First, lets make a dataframe object called "*test_data*" from the `test_data.tsv` located in the `data/` directory:

In [ ]:
```
test_data = read.csv('data/test_data.tsv', sep = '\t', header = T, row.na
```

A short description of what we just did there:

`test_data =` creates a data object (a dataframe in our case) called "*test_data*" which we can recall

We used the `read.csv()` function to read in a data file

`'/data/test_data.tsv'` is the path to the file we wish to read in

`sep = '\t'` makes `read.csv()` read "*test_data.tsv*" as a tab-delimited file (.tsv) (i.e < TAB > between columns)

`header = T` means use the first row of the data as the column headers for the dataframe (logical: T (TRUE) or F (FALSE))

`row.names = 1` uses the first column (column 1) as the rownames of the dataframe

#### 1.2. Exploring the data

We have a data object, we'll explore it a bit with *R*'s basic functions.

First, let's recall the data object, our dataframe:

In [ ]:
```
test_data
```

That's a bit too much to look at!

So, let's just look at the top rows of the data using the `head()` function:

In [ ]:
```
head(test_data)
```

...and now the last rows with the `tail()` function:

In [ ]:
```
tail(test_data)
```

Now look at the column names of the dataframe using the `colnames()` function:

In [ ]:
```
colnames(test_data)
```

You can also use the `names()` function:

In [ ]:
```
names(test_data)
```

Now the row names with `rownames()` :

In [ ]:
```
rownames(test_data)
```

We now know that columns are samples and rows are species.

To be exact it really isn't species, it is OTUs (Operational Taxonomic Units) as some assignments are higher than species (i.e genus etc.). OTU is the better way to describe taxonomic assignment, unless you focus purposefully on the species level.

The last column of the dataframe gives the taxonomic lineage of each OTU.

**How large is our dataframe?** Let's use the `dim()` function to get it's dimensions:

In [ ]:
```
dim(test_data)
```

Our test data is 54 rows by 6 columns (first value is rows, second is columns for the `dim()` output).
Taxonomic lineages (*taxonomy*) forms the final (6th) column.

Instead of using `dim()` you can return just the number of columns with `ncol()` or rows with `nrow()` :

In [ ]:
```
ncol(test_data)
nrow(test_data)
```

#### 1.3. Viewing columns of the data

A column can be isolated from the dataframe and viewed by specifying the column name.
We can find column names using `colnames()` or `names()` (see above).

Look at an individual column of the dataframe like this:

In [ ]:
```
test_data$OP01
```

This shows a list of the values in that specific column. ***Only*** the values, nothing else.

**Question:** How could this be an issue?

### 2. Data manipulation

You now know how to read in some data as a dataframe and look at it's structure. That is an important basic skill for any form of *R* analysis.

The dataframe is loaded into *R*, we have explored it, but now need to manipulate it for further analyses.

We manipulate to look at subsets of interest to us, i.e. pull out the bits we want.
In this section we will look at different methods to do this. The ways that we can organise our dataframe to better suit our needs.

#### 2.1. Dataframe indexing

Indexing is a very powerful method that can be used in multiple ways in *R*.
Indexing uses the values within a set of square brackets ( `[` `]` ) to isolate rows or columns from a dataframe.

It can be very abstract to understand to begin with, but it is a method worth becoming familiar with.
It will be important in the final stages of the *R* analysis of the sequencing data generated in the laboratory sessions.

We have "*test_data*", lets look at the first column of it:

In [ ]:
```
test_data[,1]
```

...now the first row:

In [ ]:
```
test_data[1,]
```

The position of the comma changes which we look at, columns or rows.

Columns 1 to 4:

In [ ]:
```
test_data[,1:4]
```

Rows 1 to 4:

In [ ]:
```
test_data[1:4,]
```

Columns 1 to 4 and rows 3 to 6:

In [ ]:
```
test_data[1:4, 3:6]
```

To put this into context: rows are before the comma, columns after.
Use the `dim()` , `ncol()` or `nrow()` functions to check how many columns and rows you have if needed (see above).

**An alternative:** To look at a column you can also use:

In [ ]:
```
test_data[1]
```

Dropping the comma now shows the row names too! **Note:** This only works with columns from an *R* dataframe.

**Question:** Is this more useful than `test_data$OP01` ? Why? How?

#### 2.2. Subset by indexing

Suppose we need to have the "*test_data*" dataframe without the taxonomy column?
Using dataframe indexing, in combination with the `ncol()` function (see above), we will look at the whole dataframe excluding the taxonomy column.

As "*taxonomy*" is the last column of our dataframe, `ncol(test_data)-1` will give the positional value of the second to last column.

We can now grab column 1 to the second to last column:

In [ ]:
```
test_data[,1:ncol(test_data)-1]
```

Oh look! No taxonomy column! Excellent!

Using this we can get numeric values from our dataframe.
The taxonomy column is strings - letters or words, not numeric values.

Suppose we need to get the sum all rows or columns of the dataframe? Having a string in a row will not work - it requires purely numeric values!
Removing the taxonomy column will achieve this perfectly!

Our "*test_data*" frame has samples as columns. So let's calculate the total sequencing reads of each sample by summing all values in each column using the `colSums()` function:

In [ ]:
```
colSums(test_data[,1:ncol(test_data)-1])
```

This is just what we were after. Nice.

If we would have done the following it would give an Error message as there is still the taxonomy column:

In [ ]:
```
colSums(test_data)
```

What happens if the thing we want to exclude is somewhere else in the dataframe?
The only way we can access this is via explicitic column name exclusiion.

Columns or many things can be called by a matching string (i.e. letter or words).

Lets try this:

In [ ]:
```
test_data[,!colnames(test_data) %in% "taxonomy"]
```

Look at the code above. Consider the indexing and what is happening.

By using `!` we have just said "*don't include*" column names matching, `%in%`, "*taxonomy*".
Combine this with the indexing we learnt above and it is again a powerful tool for data manipulation.
This is a very useful thing to know. Maybe a dataframe can be complicated or even need to exclude certain columns explicitly by name. A point to consider.

Lets remember this as "indexing by `%in%`" - we'll use this later.

#### 2.3. Playing with numbers

If a dataframe is entirely numeric (like "*test_data*" without the "*taxonomy*" column) then there are a multitude of mathmatics we can do to it.

Lets use indexing to crop the dataframe to pure numeric values (we know this because it is not including "*taxonomy*").

We will do this and assign a data object to it:

In [ ]:
```
ralf = test_data[1:5, 1:3]
```

Recall the object "*ralf*":

In [ ]:
```
ralf
```

Sum of all columns in "*ralf*" using the `colSums()` function:

In [ ]:
```
colSums(ralf)
```

Sum of all rows in "*ralf*" using the `rowSums()` function:

In [ ]:
```
rowSums(ralf)
```

It is important to note that a lot can be done with numeric data. This will become apparent in the next section.

Using the cells below, have a play around with the test data to familiarise yourself some more with indexing.

In [ ]:

In [ ]:

In [ ]:

In [ ]:

### 3. Sequencing data analysis

In the first seection learnt how to read in some data as a dataframe and look at it's structure. The second section demonstrated how to manipulate your data to get it to do what you want.

Now we will put all this together in the *R* analysis or the human skin microbiome metabracoding sequencing data you generated in the lab sessions.

This gets a bit complex, but don't panic - work through the stages and ask for assistance if needed.

#### IMPORTANT: Clear *R*'s memory

Before we go any further we should clean up *R*'s memory.
This will make sure there is none of the test data remaining to confuse the actual analysis. We can then begin the next section with a clean slate!

To do this run the cell below:

In [ ]:
```
rm(list = ls())
```

Remember this as you will be using it later.

#### 3.1. Load the data

Just as before we will get the data read into *R*. The file you need is located in the `data/` directory and is called "genetic_analysis_students_raw.tsv".

Lets get this data read in using the `read.csv()` function:

In [ ]:
```
my_data = read.csv('data/genetic_analysis_raw_data.tsv', sep = '\t', head
```

#### 3.1.1 Explore the data

Using the empty cells below, explore the data to get a better feel for it.
Remember the functions such as `head()`, `tail()`, `ncol()` etc.

In [ ]:

In [ ]:

In [ ]:

In [ ]:

In [ ]:

#### 3.2. Cleaning the data

Metabarcoding data is invariably noisy. It will have some OTUs taxonomically assigned that are low read counts and a lot of sequencing data can be left completely unassigned to any taxonomic rank at all.

Before carrying out any analysis of the data we must first clean the dataframe and make it suitable for purpose. Here we shall do just that.

The data needs to be numeric, so let's remove the taxonomy column:

In [ ]:
```
my_data = my_data[,!colnames(my_data) %in% "taxonomy"]
```

For our analysis the dataframe is oriented the wrong way. Rows should be samples and columns OTUs.

We need to transpose the data using the `t()` function.
But to make sure the dataframe remains as a dataframe, we have to make it a datframe again. The `as.data.frame()` function will make it a datafame.

With this in mind, lets transpose our data:

In [ ]:
```
my_data = as.data.frame(t(my_data))
```

Check to see if the data is correctly oriented in the cell below:

In [ ]:
```

```

##### 3.2.1 Removing low frequency data

Now we must deal with the low frequency reads in each sample, the OTUs that have low read counts, as they can be unreliable, and are of relatively little value in our analysis.

In our data, OTUs in a sample with less than 5 reads assigned to them can be considered noise.
These OTUs could be genuinely presnent, but can just as easily be false artefacts from sequencing/taxonomic assignment error. Either way, they are not desirable and need to be removed.

To do this we will apply a threshold to the entire dataframe. The threshold being that any value below 5 is removed and counted as 0.

Using *R* logic (TRUE or FALSE) can determine if a result is < 5. Try it:

In [ ]:
```
my_data < 5
```

***Question:*** Is this what you were expecting to see? Why?

This in combination with indexing can change the values listed as TRUE to equal 0:

In [ ]:
```
my_data[my_data < 5] = 0
```

It is inevitable that we now have OTUs with no reads assigned to them anyehere in the dataframe. We need to remove them now as it will confuse downsteam analysis.

Let's chop these from the data using the `colSums()` function, with a pinch of indexing and logic:

In [ ]:
```
my_data = my_data[!colSums(my_data) == 0]
```

As you have probably seen from exploring the data earlier, there is an OTU called "unassigned". These are the reads that failed to be assigned to any taxonomic rank in the sample and are left "unassigned". These reads form no part in our analysis and need to be removed.

To do this we will use the same indexing methods as when we deleted the "taxonomy" before. However we are removing a row here not a column - remember the comma position while indexing for rows.

We will drop the "unassigned" row:

In [ ]:
```
my_data = my_data[,!colnames(my_data) %in% "unassigned"]
```

Revisit the code above and discuss with the person next to you. The code above should be understandable by you, and you should understand how we are tansformaing the data. If not, please ask a demonstrator or discuss with the person next to you.

#### 3.3. Write out the cleaned data

The data is now clean!
We should really write it out to a file so we can come back to it easily in future without having to run all the code again.

To do this we will use the `write.table()` function.
By now the code required to do this should make some kind of sense to you when looking at it.

So let's write it out:

In [ ]:
```
write.table(my_data, file = 'data/genetic_analysis_cleaned_data.tsv', sep
```

#### Clear *R*'s memory

Here is a good point to clear the memory again.
None of the variables or objects we have justs generated above are needed anymore.

Use the cell below to clear *R*'s memory:

In [ ]:
```

```

#### -- PAUSE --

Before moving on take a moment to think about what you have just done.
Reflect on the data managemnent aspects you have just followed in the sections above.

**Questions:**

- Why did we have to clean the data?
- Can you describe what the important steps in the data cleaning were?
- Can you go over the last section and understand the code involved?

Discuss with the person next to you or ask a demonstrator.

### 4. Formatting the data

For any type of analysis the data needs to be in the correct format.
Here we will do just that: wrangle the data to how we need it.

#### In this section there are some new concepts

The `#` is used as a method to comment in the code.
It is not run, but is used to add notation to what each part is doing.

A lot of code is now grouped in each cell and executed at once.
You will need to read each cell to determine what is happening.

#### 4.1. Proportion reads and presence/absence

For our analysis we will need to create a proportion reads dataframe.
It will show the reads per OTU as a proportion of total reads per sample.

Additionally, we will need to create a presence/absence dataframe.
All the data will be transformed to `0`'s and `1`'s - indicating an OTU's presence or absence in a sample.

**In the cell below we will:**

1. Read in the cleaned data we generated above
2. Create a proportion reads dataframe
3. Create a presence/absence dataframe

In [ ]:

```
# read in the new cleaned dataset:
my_data_cleaned = read.csv('data/genetic_analysis_cleaned_data.tsv', sep

# make a proportion reads dataframe:
my_data_prop = my_data_cleaned
my_data_prop = my_data_prop/rowSums(my_data_prop)

# make a presence/absence dataframe:
my_data_pa = my_data_cleaned
my_data_pa[my_data_pa > 0] = 1
```

The processes used in the cell above should make sense to you now you have manipulated data.
If unsure, discuss with the person next to you or ask a demonstrator.

### 4.2. Subsetting by sample site

We njeed to make an OTU presence/absence dataframe per sample site.
As the data is based on limbs (arms and legs) we will make a list of seperate data frames per limb.

To do this we need to isolate each limb from the presence/absence data we created above.

The `grep()` function (*very powerful*) searches for a match to a regular expression in a data set.
In our data the limbs have a few unique characters associated with them.

Look at the `rownames()` of "*my_data_pa*" in the cell below:

In [ ]:

```
```

You will see that the first two characters of each sample name are distinct to each limb. Right?

Using this we can tell `grep()` what to look for.

Try this to find samples from the left leg:

In [ ]:

```
grep("LL", rownames(my_data_pa))
```

The output of `grep()` has just shown us the row numbers that have "LL" in their name.
These are from the left leg, well "*leg left*" as the sample naming system goes.

Using `grep()` we will now subset the data as we need it.

However, we need it in a manner that we can easily recall for analysis.
For this we will add each subset of the dataframe to a list.

**We will make a list of dataframes!**

Dataframe lists are immensely useful as we will see in the coming sections.

But, for now...

Let's hack the data into body parts!

In [ ]:

```
l_leg = my_data_pa[grep("LL", rownames(my_data_pa)),]
r_leg = my_data_pa[grep("LR", rownames(my_data_pa)),]
l_arm = my_data_pa[grep("AL", rownames(my_data_pa)),]
r_arm = my_data_pa[grep("AR", rownames(my_data_pa)),]

body_parts = list(l_leg, r_leg, l_arm, r_arm)

names(body_parts) = c("left_leg", "right_leg", "left_arm", "right_arm")
```

**Explore the list of dataframes.**

For example:

```
In [ ]: body_parts[1]
```

```
In [ ]:
```

```
In [ ]:
```

```
In [ ]:
```

### 5. Ecological analysis

Now we have the data in multiple suitable formats.
It is now stored in memory and ready to be used.

We can start to look at the data in the ecological manner that we want. We can consider the body parts separate environments just like the larger scale environments you are used to.

#### 5.1. OTU richness

First we need to look at the OTU richness per sample per limb.

The cell below will do just this:

```
In [ ]: # create an empty dataframe based on my_data_pa:
my_data_rich = my_data_pa[FALSE]

# Add columns for species richness - sum of rows from my_data_pa:
my_data_rich$richness = rowSums(my_data_pa)

# Add limbs in as numbers rather than name:
my_data_rich$limb = rep(c(1, 2, 3, 4), each = 3)

# recall the data:
my_data_rich
```

The limbs are numbered. This is done because *R* alphabetically orders things and will mess with our order of samples (legs, then arms).
We need them in the order they are given. This keeps the order of samples and is important for the following plot output.

Let's make our first meaningful plot of the analysis...

##### OTU richness boxplot

Now we will use the following code to make a boxplot of OTU richness per sample per site:

In [ ]:
```
boxplot(my_data_rich$richness ~ my_data_rich$limb,
        # name the individual bars according to sample:
        names = c("left leg", "right leg", "left arm", "right arm"),
        # add y axis label:
        ylab = 'OTU richness',
        # x axis label:
        xlab = 'Sample site',
        # x and y axis label size:
        cex.lab = 1.5,
        # make the axis tick labels horizontal:
        las = 1)
```

What does the code in the above cell do? It has been annotated, butu do ***you*** understand it? Could ***you*** do it by yourself?

**Google for details as required.**

It is important you understand what is going on.

Once you have thought/Googled about it, discuss with the person next to you or ask a demonstrator.

##### Total OTU richness barplot

Now we will use the `body_parts` list object we created earlier. Remember?

For each limb we will now calculate the total OTU richness.
We need to create a list of total OTU richness values for each limb with names to determine which limb it is.

In [ ]:
```
l_leg_rich = sum(colSums(body_parts$left_leg) > 0)
r_leg_rich = sum(colSums(body_parts$right_leg) > 0)
l_arm_rich = sum(colSums(body_parts$left_arm) > 0)
r_arm_rich = sum(colSums(body_parts$right_arm) > 0)

total_richness = c(l_leg_rich, r_leg_rich, l_arm_rich, r_arm_rich)
names(total_richness) = c("left leg", "right leg", "left arm", "right arm
```

Does the above code make sense? It should at least make some. Ask a demonstrator if unsure.

Let's plot the barplot:

In [ ]:
```
barplot(total_richness, beside = T)
```

Now we have another plot. What does it mean?

Discuss the plot results with the person next to you or ask a demonstrator.

In [ ]:
```

```

### 5.2. Vegan analysis

The analysis now gets interestingly large.

#### Vegan

Vegan is an *R* package for ecological and community data anslysis.

**Vegan is the package that will make the important figures!**

Before we can use it, we need to load it.
Remember we are in an enclosed "**environment**" and everything required for the practical is available?
So all we need to do is call the required packages from the "**environment**".

Call in the **Vegan** package from *R*'s library with the `library()` function:

In [ ]:
```
library(vegan)
```

#### 5.2.1. OTU accumulation

We have the `body_parts` list created (from above).

recall `body_parts` in the cell below:

In [ ]:
```
body_parts
```

For the dataframes in `body_parts` we will have to "*loop*" through them and generate a "*species accumulation*" for each limb.

As you may have just guessed, "*looping*" in *R* is a vital skill for analysis.

Lets "*loop*" through something as an example:

In [ ]:
```
# make a concatinated list of numeric values:
limp = c(1, 2, 3, 4)

# loop through them and print each one:
for(i in limp){print(i)}
```

As you can see "*looping*" is logically simple and very useful.

Now we will apply it ot our data and use Vegan to calculate some stuff on the way.

We will use Vegan's species accumulation function: `specaccum()`:

In [ ]:
```
# make an empty list:
species_accum = list()

# specify the "limbs":
limbs = c("left_leg", "right_leg", "left_arm", "right_arm")

# loop through the limbs:
for(limb in limbs){
        # generate "species " accumulation:
```

```
        vegan_accum = specaccum(body_parts[[limb]],
              method = "exact",
              permutations = 100)
        # add to "species_accum" list:
        species_accum[[limb]] = vegan_accum
}
```

There may have been some warnings there, do not worry. All is OK! We have just generated something!

**Question:** What have we generated? What does it mean?

Now let's plot it:

In [ ]:
```
# make the plot have a 2x2 grid of plots:
par(mfrow = c(2, 2))

#list of "limbs":
limbs = c("left_leg", "right_leg", "left_arm", "right_arm")

# loop through limbs and plot:
for(limb in limbs){
    plot(species_accum[[limb]],
        ylim = c(50, 300),
        main = sub('_', ' ', limb),
        xlab = 'Samples',
        ylab = 'OTUs detected',
        las = 1)
    # add thicker lines to the main line of each plot:
    lines(species_accum[[limb]]$richness, lwd = 3)
}
```

We now have accumulation curves for OTUs. Is this meaningful?
Can you determine if the sampling number was sufficient to capture the maxima of OTUs?

Discuss the results with the person next to you or ask a demonstrator.

#### 5.2.2. NMDS ordination

**Now we get to plot a Non-metric multidimensional scaling (NMDS) ordination!**

As already mentioned Vegan has many purposes. This is the one that will show you the most important results.

First we must make an "environment" dataframe - a descriptive set of data for reference. It contains a lot of "environmental" info.
In our case, the sample name, limb, side (left or right) and side the limb is on (side + limb).

Here we use a function, `rep()`, to do a lot of work. **Google** it and see how it works.

The cell below does everything needed:

In [ ]:
```
# create an env dataframe for use in vegan:
veg_env = my_data_prop[FALSE]
# set sample name from rownames of data:
veg_env$site = rownames(veg_env)
# make the limb column:
veg_env$limb = rep(c('leg', 'arm'), each = 6)
# make the side column:
veg_env$side = rep(rep(c('left', 'right'), each = 3), 2)
# make the side/limb clumn:
veg_env$side_limb = paste(veg_env$side, veg_env$limb, sep = '_')
```

##### 5.2.3. Vegan metaMDS

We now feed in the proportion data, `my_data_prop`, to the ordination function of Vegan: `metaMDS()` and assign it to an object `ord`.

Run the code:

In [ ]:
```
ord = metaMDS(my_data_prop, k = 3, try = 100, trymax = 10000, distance =
```

##### Vegan NMDS plot

Now we have the data in the correct formats, lets plot the plot...

Run the wall of code below:

In [ ]:
```
# empty plot of ord, no axes:
plot(ord, disp = "sites", type = 'n', axes = F, xlab = 'NMDS1', ylab = 'N

# add correct x and y axes:
axis(1, at = seq(-1,2,0.5), cex.axis = 1, padj = -0.5, tck = -0.01, lwd =
axis(2, at = seq(-1,1,0.5), cex.axis = 1, las = 1, tck = -0.02, lwd = 0,

# add vertical and horizontal lines to show "0, 0":
abline(h = 0, v = 0, lwd = 2, col = 'grey')

# add ordination circumference elipses with corerect colours:
ordiellipse(ord,
            veg_env$side_limb, kind = "ehull", lwd = 2,
            col = rep(c('green', 'red'), each = 2))

# add sample names as text to points
text(ord$points[,1:2], labels = veg_env$site, pos = 4)

# add points to plot with colours specific to sites:
points(ord,
       disp = "sites",
       pch = ifelse(veg_env$limb == 'arm', 21, 24),
       bg = ifelse(veg_env$side == 'left', 'green', 'red'),
       cex = 2,
       col = 'black',
       lwd=0.5)

# add box around plot:
box('plot', lwd=2)
```

Can you interpret the plot correctly? Can you understand the code that made it?

Discuss the plot results with the person next to you or ask a demonstrator.

### 6. Session complete

#### Before you leave, please make sure you understand what you have just done. Ask a demonstrator.

It is important that you know what you did. Spamming keys is one thing, but knowing what they did is the key.
I'd like you all to walk away slightly wiser, not just dumbstruck!

##### Tomorrow:

We will revisit this and make some plots of the data that you will use in your final report.

### GENETIC ANALYSIS: FIGURE CREATION IN *R*

### 1. Data presentation

Plotting the data as a meaningful figure is an important part of reporting on any analysis work.
This is another form of scientific communication.

Today there are a lot of *R* code blocks to run. We think that you will be able to read this code, and understand what is going on. We have included code comments to help you.

Your work is to run the code on your own data, and then to modify it to improve the figures generated (axes, colours, plot clarity). These figures will be used in your assignment write up.

### The art of science

In this session we will concentrate on making suitable figures to be included in your assignment for the Genetic Analysis practical.

You will modify the existing code used to make the plots from the last session, adding to them and customising as you see fit (within reason OFC).

#### 1.1. Theme

When creating plots for a report or manuscript it is a good idea to have a theme in mind.

By "theme", at a minimum, I mean a colour scheme repeated across all relevant plots. Font size, axis labelling, line width and figure resolution all contribute to the theme too.

Having a theme for your figures will act as a unifying factor, a continuity that will make the manuscript seem more of a purposeful entity rather than a disparate smattering of figures. A "*random-acts-of-plotting*" piece of work is certainly not nice, nor is it easy to look at.

#### 1.2. Base *R*

All the figures will be generated using basic *R*.
There are excellent packages available (e.g. *ggplot*) that can allow for the creation of immensely customisable plots, however for this practical we are keeping it with what you already know with *baseR*.

The limited functionality of basic *R* plotting will introduce you to many concepts of figure production.
These can then be improved on as you progress in your degree or career as a genetic analysist.

### 2. Recreate the data

We are in a new Jupyter Notebook, therefore the variables, dataframes and objects created in the last session do not exist.

We will need to recreate them. This means to basically rerun the entire anlysis to get the data in the correct format and ready to plot again.
This is as simple as putting all the releant code in a single cell and running it.

**In the cell below we will rerun the final anlysis from the last session**

The data will all be formatted and ready for plotting.
**Note:** Comments have been added to the code so we can see which bits are doing what.

**When you are ready, run the cell:**

In [ ]:

```
# ====================================================
# FORMATTING DATA SECTION
# ====================================================

# MAKE PROPORTION READS AND PRESENCE/ABSENCE DATAFRAMES

# read in the new cleaned dataset:
my_data_cleaned = read.csv('data/genetic_analysis_cleaned_data.tsv', sep

# make a proportion reads dataframe:
my_data_prop = my_data_cleaned
my_data_prop = my_data_prop/rowSums(my_data_prop)

# make a presence/absence dataframe:
my_data_pa = my_data_cleaned
my_data_pa[my_data_pa > 0] = 1


# ====================================================

# SUBSET THE DATA BY LIMB

# chopping up the data by limb:
l_leg = my_data_pa[grep("LL", rownames(my_data_pa)),]
r_leg = my_data_pa[grep("LR", rownames(my_data_pa)),]
```

```r
l_arm = my_data_pa[grep("AL", rownames(my_data_pa)),]
r_arm = my_data_pa[grep("AR", rownames(my_data_pa)),]

body_parts = list(l_leg, r_leg, l_arm, r_arm)

names(body_parts) = c("left_leg", "right_leg", "left_arm", "right_arm")


# ====================================================

# CREATE OTU RICHNESS DATAFRAME

# create an empty dataframe based on my_data_pa:
my_data_rich = my_data_pa[FALSE]

# Add columns for species richness - sum of rows from my_data_pa:
my_data_rich$richness = rowSums(my_data_pa)

# Add limbs in as numbers rather than name:
my_data_rich$limb = rep(c(1, 2, 3, 4), each = 3)


# ====================================================

# CREATE TOTAL OTU RICHNESS LIST

# total  richness per limb:
l_leg_rich = sum(colSums(body_parts$left_leg) > 0)
r_leg_rich = sum(colSums(body_parts$right_leg) > 0)
l_arm_rich = sum(colSums(body_parts$left_arm) > 0)
r_arm_rich = sum(colSums(body_parts$right_arm) > 0)

# list of limb OTU richness:
total_richness = c(l_leg_rich, r_leg_rich, l_arm_rich, r_arm_rich)

# add names for list:
names(total_richness) = c("left leg", "right leg", "left arm", "right arm


# ====================================================
# VEGAN SECTION
# ====================================================

# import Vegan library:
library(vegan)


# ====================================================

# VEGAN ENVIRONMENT DATAFRAME

# create an env dataframe for use in vegan:
veg_env = my_data_prop[FALSE]

# set sample name from rownames of data:
veg_env$site = rownames(veg_env)

# make the limb column:
veg_env$limb = rep(c('leg', 'arm'), each = 6)
```

```
# make the side column:
veg_env$side = rep(rep(c('left', 'right'), each = 3), 2)

# make the side/limb clumn:
veg_env$side_limb = paste(veg_env$side, veg_env$limb, sep = '_')


# ====================================================

# LIST OF SPECACCUM OBJECTS PER LIMB

# make an empty list:
species_accum = list()

# specify the "limbs":
limbs = c("left_leg", "right_leg", "left_arm", "right_arm")

# loop through the limbs:
for(limb in limbs){
        # generate "species " accumulation:
        vegan_accum = specaccum(body_parts[[limb]],
              method = "exact",
              permutations = 100)
        # add to "species_accum" list:
        species_accum[[limb]] = vegan_accum
}


# ====================================================

# VEGAN ORDINATION CREATION

# vegan ord:
ord = metaMDS(my_data_prop, k = 3, try = 100, trymax = 10000, distance =
```

That was easy wasn't it? The entirety of the last session in a cell. Nice.

**Questions:**

1. Can you tell exactly what was done in the cell?
2. Do the comments help?

#### Clear *R*'s memory

If you ever feel like having a complete reset of *R*'s memory, run the cell below:

In [ ]:
```
rm(list = ls())
```

### 3. Plot to .png

The simplest way to get a figure from *R* to an image file (in this case a .png) via the code is to use the `png()` function.

For example:

In [ ]:
```
png('plots/test.png', width = 1080, height = 1080, units = 'px', pointsiz

# everything for the plot goes here:
plot(0, 0)

dev.off()
```

A short description of the function:

`'plots/test.png'` is the path to the location where the image is to be saved.

`width = 1080` the width of the image in units.

`height = 1080` the height of the image in units.

`units = 'px'` the units the image is measure in, in this case pixels.

`pointsize = 30` the font size of the image.

**Important:** The function needs to be closed after the plot commands.

`dev.off()` does this.

Have a look at the plot created from the cell above (it is in the "**plots**" directory as "**test.png**" in the file browser to the left of the notebook).

##### Simple .png save

To simply save a plot as an image, the easiest way is to copy the `png()` command from the cell above.
Then paste it as the first line of the cell with the plot code in.
Add `dev.off()` as the very last line.

### 4. Customising your plots

The cells below have the relevant code to produce the plots from the last session's analysis.

The code has been expanded to include all the functionality needed for modification of colour, line width and many other aspects.
Comments have been added to show what each part relates to on the plot.

#### 4.1. Safety first!

Use the `+` button in the top bar of the notebook to create a new cell.
Copy the code of the cell you want to modify into it.
This will stop you from irreversibly ruining the code, and keep your experiment reproducible.

##### 4.1.1. Have fun, go mad

Tweak all the variables! Choose different colours!
Run the cell to look at any changes you have made to the plot. Continue tweaking.

**Remember your "*theme*" and make your plots look good**.

#### 4.2. Your customised plot

When you are happy with your output, check with a demonstrator.

Next, add in the `png()` and `dev.off()` functions in the correct places in the cell and make sure the plot looks good as the final .png image.
Check them via the file browser on the left of the notebook. They should be in the "**plots**" directory if you have included the path correctly.

### 5. The plots

##### OTU richness boxplot

In [ ]:

```
boxplot(my_data_rich$richness ~ my_data_rich$limb,
        names = c("left leg", "right leg", "left arm", "right arm"),
        # y axis title:
        ylab = 'OTU richness',
        # x axis title:
        xlab = 'Sample site',
        # size of axis labels:
        cex.lab = 1.5,
        # size of axis tick labels:
        cex.axis = 1,
        # colours of boxes:
        col = 'grey80',
        las = 1)
```

##### Total OTU richness barplot

In [ ]:

```
barplot(total_richness,
        beside = T,
        # size of axis tick labels:
        cex.axis = 1,
        # size of x axis labels:
        cex.names = 1,
        #y axis label:
        ylab = 'OTU richness',
        # size of y axis label:
        cex.lab = 1.5,
        # bar colours:
        col = 'grey80',
        # colour of bar borders:
        border = 'black,'
        las = 1)
```

```
# add line across the bottom of the plot:
abline(h = 0)
```

#### OTU accumulation plot

In [ ]:

```
# make the plot have a 2x2 grid of plots:
par(mfrow = c(2, 2))

#list of "limbs":
limbs = c("left_leg", "right_leg", "left_arm", "right_arm")

# loop through limbs and plot:
for(i in 1 : length(limbs)){
    plot(species_accum[[limbs[i]]],
        # y axis upper and lower values:
        ylim = c(50, 250),
        # x axis upper and lower values:
        xlim = c(0, 4),
        main = sub('_', ' ', limbs[i]),
        # x axis title:
        xlab = 'Samples',
        # y axis label:
        ylab = 'OTUs detected',
        # size of axis labels:
        cex.lab = 1,
        # colour of lines:
        col = 'black',
        las = 1)

    # add thicker lines to the main line of each plot:
    lines(species_accum[[limbs[i]]]$richness,
          # line width of main line:
          lwd = 3,
          # colour of main line:
          col = 'black')
}
```

#### Vegan NMDS ordination plot

In [ ]:

```
# empty plot of ord, no axes:
plot(ord,
     disp = "sites",
     type = 'n',
     axes = F,
     # x axis upper and lower values:
     xlim = c(-1, 1),
     # y axis upper and lower values:
     ylim = c(-1, 1),
     # x axis label:
     xlab = 'NMDS1',
     # y axis label:
     ylab = 'NMDS2',
     bty = 'n',
     # size of axis labels:
     cex.lab = 1.5)

# add correct x and y axes:
```

```
axis(1, at = seq(-1,2,0.5), cex.axis = 1, padj = -0.5, tck = -0.01, lwd =
axis(2, at = seq(-1,1,0.5), cex.axis = 1, las = 1, tck = -0.02, lwd = 0,

# add vertical and horizontal lines to show "0, 0":
abline(h = 0, v = 0, lwd = 2, col = 'grey')

# add ordination circumference elipses with correct colours:
ordiellipse(ord,
            veg_env$side_limb,
            kind = "ehull",
            # line width of elipses:
            lwd = 2,
            # colours of elipses:
            col = rep(c('green', 'red'), each = 2))

# add points to plot with colours specific to sites:
points(ord,
       disp = "sites",
       # shape of points:
       pch = ifelse(veg_env$limb == 'arm', 21, 24),
       # colour of points:
       bg = ifelse(veg_env$side == 'left', 'green', 'red'),
       # size of points:
       cex = 2,
       # outline colour of points:
       col = 'black',
       # line width of points outline:
       lwd = 0.5)

# add sample names as text to points:
text(ord$points[,1:2],
     # size of text:
     cex = 1,
     # label names to use:
     labels = veg_env$site,
     # position of the text around point:
     pos = 4)

# add box around the plot:
box('plot',
    # width of box line:
    lwd = 2)
```

### 6. Session complete

You have just made some manuscript quality figures for your Genetic Analysis assignment.

#### Well done!

##### Before you leave...

Make sure you have downloaded all your figures.
You don't want to have lost them to the ether!

**Ask a demonstrator if there are any issues.**

#### Scientific Reproducibility

All your figures have been made in an entirely reproducible manner. You could start with the same data and create the same figures just by running your notebook. Other scientists could reproduce your work in the same way. This is a great way to carry out research.
