## Supplementary 3: Risk assessment for "Who Grows There? A Course-based Undergraduate Research Experience to explore the human microbiome through 16S DNA metabarcoding"

| UNIVERSITY OF Hull<br>Department of Biological and Marine Sciences | PROCEDURAL RISK ASSESSMENT AND COSHH FORM | | | Page 1 |
|---|---|---|---|---|
| | ref: BMS101146v1 | Date: 26/10/22 | Group: Teaching | Overall Risk Level: Low |

| PROCEDURE TITLE: | 500697 Genetic Analysis Lab week 1 extraction quantification and PCR of DNA | |
|---|---|---|
| Ethics Approval ☐ YES x NO | Home Office Licence ☐ YES x NO | |
| Reference No: | Home Office No: | Other: |
| Further information must be provided in the appropriate section within the BioCOSHH part of this form. | | |

| Name and status of assessor: | I consider that all precautions listed are adequate and reduce risks to an acceptable level |
|---|---|
| Graham Sellers | Signature: [signature] Date: 26/10/2022 |
| Head of Group (Authorisation): Graham Sellers | I am satisfied that the risk assessment completed is suitable and sufficient<br>Signature: [signature] Review Date: Oct 2023 |
| ☒ School/Departmental Safety Officer<br>☐ University Biological Hazards Officer | I have reviewed this assessment<br>Countersignature: RDonnelly Date: 26/10/2022 |

**Location(s): Hardy South teaching lab**

**Who might be affected:**

☒ Staff ☒ Undergraduate Students ☒ Postgraduate Students ☐ Cleaners ☐ Maintenance Contractors ☐ Visitors ☐ General Public
☐ Young Persons (<18 yrs) ☐ Elderly ☒ New and expectant mothers ☒ Immune-compromised Persons
☐ Other:

| UNIVERSITY OF Hull<br>Department of Biological and Marine Sciences | **PROCEDURAL RISK ASSESSMENT AND COSHH FORM** | | | Page 2 |
|---|---|---|---|---|
| | **ref: BMS101146v1** | **Date: 26/10/22** | **Group: Teaching** | **Overall Risk Level: Low** |

| | | |
|---|---|---|
| **Risk to pregnancy:** ☒ Yes ☐ No | **Lone Working permitted:** ☐ Yes ☒ No | **Out of hours permitted:** ☐ Yes ☒ No |

### PROCEDURE DESCRIPTION

**Technical Summary outlining the Procedure being followed**

**Corresponding Detailed Protocol/SOP Reference No:** **(Where currently not available must be provided within 3 months of the approved date)**

Genetic Analysis lab manual

Students will be performing the following procedures:

- Extract DNA from bacterial isolates (*E. coli* K12 frozen liquid medium culture)
- Assess the quantity of DNA extraction through Qubit fluorometer
- PCR with universal 16S primers and gel electrophoresis
- Library preparation PCR for MinION sequencing

**All waste must be autoclaved by technicians prior to disposal.**

Clinical isolates are of Hazard group 1 only as outlined in the Advisory committee on Dangerous Pathogens “The Approved List of Biological Agents”

**Extra information:**

All microbiological cultures are unlikely to spread to the community and there will be effective prophylaxis or treatment available. Risk is very low – all culture work has been carried out prior to the practical and the bacterial samples are frozen.

**For work to be undertaken by Immunocompromised people/pregnant women: an individual risk assessment must be completed to evaluate the risk on a case by case situation**

| RISK RATING | | | | | |
|---|---|---|---|---|---|
| L = Likelihood of event occurring | Rare = 1 | Unlikely = 2 | Possible = 3 | Likely = 4 | Almost certain = 5 |
| C = Consequences of event occurring | Insignificant = 1 | Minor = 2 | Moderate = 3 | Major = 4 | Catastrophic = 5 |

### EQUIPMENT

| Equipment used | How might someone be harmed? | RISK (L X C) | Control Measures | Residual RISK (L X C) |
|---|---|---|---|---|
| | | **MuDNA extraction** | | |
| Benchtop centrifuge | • Electrocution<br>• Damage to rotor if unbalanced/centrifuge may travel and fall from surface<br>• Flying parts | 3 x 5 = 15 | • Ensure that users are trained in how to use the equipment and understand the need to balance the rotor<br>• Perform visual inspection before use<br>• Annual PAT testing must be within date | 1 X 5 = 5 |
| TissueLyser | • Electrocution | 3 x 5 = 15 | • Perform visual inspection before use<br>• Annual PAT testing must be within date | 1 X 5 = 5 |
| Thermomixer | • Electrocution | 3 x 5 = 15 | • Perform visual inspection before use<br>• Annual PAT testing must be within date | 1 X 5 = 5 |
| Hulamixer | • Electrocution | 3 x 5 = 15 | • Perform visual inspection before use<br>• Annual PAT testing must be within date | 1 X 5 = 5 |
| Vortex | • Electrocution | 3 x 5 = 15 | • Perform visual inspection before use<br>• Annual PAT testing must be within date | 1 X 5 = 5 |
| Pipettes | • Repetitive strain injury | 3 x 3 = 9 | • Ensure pipette action is smooth and light<br>• If hands/wrists start to ache, stop work. | 1 x 3 = 3 |

| PCR and Gel electrophoresis | | | | |
|---|---|---|---|---|
| Benchtop centrifuge | • Electrocution<br>• Damage to rotor if unbalanced/centrifuge may travel and fall from surface<br>• Flying parts | 3 x 5 = 15 | • Ensure that users are trained in how to use the equipment and understand the need to balance the rotor<br>• Perform visual inspection before use<br>• Annual PAT testing must be within date | 1 X 5 = 5 |
| Thermal cycler (PCR machine) | • Electrocution<br>• Burns from heated internal lid | 3 x 5 = 15 | Ensure that users are trained in how to use the machine. Perform visual inspection before use. Users should not open the lid during program operation and should not touch the internal heated lid. Annual PAT testing, must be within date. | 1 x 5 = 5 |
| Microwave | • Overheating and spillage/explosion of agarose in microwave.<br>• Exposure to microwaves<br>• Electrocution | 4 x 5 = 20 | • Do not heat agarose in bottles with lid on.<br>• Take care when taking hot agarose out of the microwave. Use thermal gloves and safety glasses.<br>• Perform visual inspection before use<br>• Annual PAT testing must be within date | 1 x 5 = 5 |
| Balance | • Electrocution | 3 x 5 = 15 | • Perform visual inspection before use<br>• Annual PAT testing must be within date | 1 x 5 = 5 |
| Electrophoresis tank | • Electrocution<br>• Spillage of buffer | 3 x 5 = 15 | • Perform visual inspection before use<br>• Annual PAT testing must be within date<br>• Ensure tank is on a flat surface and the outside of the tank is dry. | 1 x 5 = 5 |
| Powerpack | • Electrocution | 2 x 5 = 10 | • Perform visual inspection before use<br>• Annual PAT testing must be within date | 1 x 5 = 5 |
| UV Gel imager | • Electrocution<br>• UV radiation | 3 x 5 = 15 | • Perform visual inspection before use<br>• Annual PAT testing must be within date<br>• Only to be used by trained user | 1 x 5 = 5 |

| | | | | |
|---|---|---|---|---|
| | | | • Instrument design means that users will not be exposed to UV radiation during normal use | |
| **Qubit quantification** | | | | |
| Qubit fluorometer | • Electrocution | 3 X 5 = 15 | • Perform visual inspection before use<br>• Annual PAT testing must be within date | 1 X 5 = 5 |
| Benchtop centrifuge | • Electrocution<br>• Damage to rotor if unbalanced/centrifuge may travel and fall from surface<br>• Flying parts | 3 x 5 = 15 | • Ensure that users are trained in how to use the equipment and understand the need to balance the rotor<br>• Perform visual inspection before use<br>• Annual PAT testing must be within date | 1 X 5 = 5 |
| Pipettes | • Repetitive strain injury | 3 x 3 = 9 | • Ensure pipette action is smooth and light<br>• If hands/wrists start to ache, stop work. | 1 x 3 = 3 |
| **Nanodrop quantification** | | | | |
| Nanodrop spectrophotometer | • Electrocution | 3 X 5 = 15 | • Perform visual inspection before use<br>• Annual PAT testing must be within date | 1 X 5 = 5 |
| Pipettes | • Repetitive strain injury | 3 x 3 = 9 | • Ensure pipette action is smooth and light<br>• If hands/wrists start to ache, stop work. | 1 x 3 = 3 |

### CHEMICAL SUBSTANCES

**Take note of Concentration and Volumes of Chemicals Used e.g. stock versus working diluted concentrations**

| Substance Name | Quantity Used | WEL Value | Hazardous Properties (Incl. form/route of entry) | Risk (L x C) | Control Measures | Residual Risk (L x C) | Waste Disposal | Emergency Action in the Event of Spillage/Exposure |
|---|---|---|---|---|---|---|---|---|
| **MuDNA extraction** | | | | | | | | |
| Guanidine thiocyanate | 1.3g/100ml | Not available | Liquid/solid ingestion/absorption<br><br>Irritant to skin and mucous membranes.<br><br>Harmful by inhalation<br><br>harmful gas released on contact with acids | 3 x 4 = 12 | Safety glasses, nitrile gloves, lab coat.<br>Ensure adequate ventilation<br>Do not expose to acid | 1 x 4 = 4 | Retain residue in waste bottle contained in fume cupboard. Disposal by incineration through chemistry stores. | Dilute and mop up with towels. Large spills must be disposed of through the Chemistry department. |
| Trisodium phosphate | 6 .5g/100ml | Not available | Liquid/solid ingestion/absorption<br><br>Irritant to skin and mucous membranes.<br><br>Serious eye irritation | 3 x 3 = 9 | Safety glasses, nitrile gloves, lab coat.<br>Ensure adequate ventilation | 1 x 3 = 3 | Retain residue in waste bottle contained in fume cupboard. Disposal by incineration through chemistry stores. | Dilute and mop up with towels. Large spills must be disposed of through the Chemistry department. |
| Sodium chloride | 0.1g/100ml | Not available | • Liquid/solid<br>• none | 1 x 1 = 1 | • n/a | 1 x 1 = 1 | n/a | n/a |

| | | | | | | | | |
|---|---|---|---|---|---|---|---|---|
| Tris/HCl Buffer<br>(Tris(hydroxymethyl)aminomethane) | 5ml | Not available | • Liquid/solid<br>• Ingestion/absorption<br>• Skin irritation<br>• Eye irritation<br>• Specific target organ toxicity<br><br>• May be harmful if inhaled or swallowed. | 3 x 4 = 12 | • Safety glasses, nitrile gloves, lab coat.<br>• Work in a well ventilated area | 1 x 4 = 4 | Retain residue in waste bottle contained in fume cupboard. Disposal through chemistry stores. | Spillage: Sweep up and shovel. Keep in suitable, closed containers for disposal. When liquid: use paper towels to contain spillage. Contaminated paper towels can be disposed of through landfill waste.<br>Fire: use water spray, alcohol resistant foam, dry chemical or carbon dioxide. |
| EDTA 0.2mM | 1.86g in 250mls | Not available | • Liquid/solid<br>• Ingestion/absorption<br>• Eye irritation<br>• May be harmful if inhaled, swallowed or absorbed through skin. | 3 x 4 = 12 | Safety glasses, nitrile gloves, lab coat. | 1 x 4 = 4 | Retain residue in waste bottle contained in fume cupboard. Disposal through chemistry stores. | Spillage: Pick up and arrange disposal without creating dust (dampen with water if necessary) Sweep up and shovel. When liquid: use paper towels to contain spillage.<br>Contaminated paper towels can be disposed of through landfill waste.<br>Large spills must be disposed of through the Chemistry department. |

| | | | | | | | | |
|---|---|---|---|---|---|---|---|---|
| | | | | | | | | Fire: use water spray, alcohol resistant foam, dry chemical or carbon dioxide. |
| Sodium Lauryl Sulphate | 20g/100ml | Not availabl e | • Solid<br>• Flammable<br>• Harmful if swallowed or if inhaled<br>• Causes skin irritation.<br>• Causes serious eye damage.<br>• May cause respiratory irritation. | 3 x 4 = 12 | Safety glasses, nitrile gloves, lab coat.<br>Use in a well-ventilated area<br>Keep away from all sources of ignition | 1 x 4 = 4 | Retain residue in waste bottle contained in fume cupboard. Disposal through chemistry stores. | Spillage: Sweep up and shovel. Keep in suitable, closed containers for disposal through the Chemistry department.<br>Fire: use water spray, alcohol resistant foam, dry chemical or carbon dioxide. |
| Ammonium acetate | 25.69g/100 ml | Not availabl e | Liquid/solid ingestion/absorption<br><br>Irritant to skin and mucous membranes.<br><br>Serious eye irritation | 3 x 4 = 12 | Safety glasses, nitrile gloves, lab coat.<br>Ensure adequate ventilation | 1 x 4 = 4 | Retain residue in waste bottle contained in fume cupboard. Disposal by incineration through chemistry stores. | Dilute and mop up with towels. Large spills must be disposed of through the Chemistry department. |
| Aluminum Ammonium Sulphate Dodecahydrat e | 2.72g/10ml | Not availabl e | Liquid/solid ingestion/absorption<br><br>Possible irritant to skin and mucous membranes. | 3 x 4 = 12 | nitrile gloves, lab coat<br>Ensure adequate ventilation | 1 x 4 = 4 | Retain residue in waste bottle contained in fume cupboard. Disposal by incineration through chemistry stores. | Dilute and mop up with towels. Large spills must be disposed of through the Chemistry department. |

| | | | | | | | | |
|---|---|---|---|---|---|---|---|---|
| Calcium chloride dihydrate | 3g/100ml | Not available | Liquid/solid ingestion/absorption<br><br>Irritant to skin and mucous membranes.<br><br>Serious eye irritation | 3 x 4 = 12 | **Safety glasses**, nitrile gloves, lab coat.<br>Ensure adequate ventilation | 1 x 4 = 4 | Retain residue in waste bottle contained in fume cupboard. Disposal by incineration through chemistry stores. | Dilute and mop up with towels. Large spills must be disposed of through the Chemistry department. |
| Guanidine HCl | 52.54g/100 ml | Not available | Irritant to skin and mucous membranes.<br><br>Harmful by inhalation<br><br>harmful gas released on contact with acids | 3 x 4 = 12 | Safety glasses, nitrile gloves, lab coat.<br>Ensure adequate ventilation<br><br>Do not expose to acid | 1 x 4 = 4 | Retain residue in waste bottle contained in fume cupboard. Disposal by incineration through chemistry stores. | Dilute and mop up with towels. Large spills must be disposed of through the Chemistry department. |
| Ethanol | <2.5 litres | 1000ppm<br>1920mg/m$^3$ | Liquid Absorption/Ingestion<br>• Harmful if swallowed<br>• May be harmful if inhaled<br>• May be harmful if absorbed through the skin<br>• May cause eye irritation<br>• Highly flammable<br>• Mutagenic for mammalian somatic cells | 3 x 4 = 12 | Wear lab coats and nitrile gloves.<br>Avoid contact with the skin.<br>Keep away from all sources of ignition.<br>Work in a well ventilated area. | 1 x 4 = 4 | Ethanol in small amounts can be rinsed down th sink with plenty of water<br>If ethanol is mixed with other chemicals, collect the residues in a bottle and dispose of through the Chemistry department. | Spillage: Wash with copious amounts of water. If a large amount (over 2 litres) evacuate the area and treat as a fire hazard, by raising the alarm.<br>Fire: Use water spray, alcohol resistant foam, dry chemical or carbon dioxide. |

| | | | | | | | | |
|---|---|---|---|---|---|---|---|---|
| Hydrochloric Acid | 2.5ml | | • Causes severe skin burns and eye damage<br>• May cause respiratory irritation | 3 x 4 = 12 | Safety glasses, nitrile gloves, lab coat. | 1 x 4 = 4 | Retain residue in waste bottle contained in fume cupboard. Disposal by incineration through chemistry stores. | Dilute and mop up with towels. Large spills must be disposed of through the Chemistry department. |
| **PCR and Gel electrophoresis** | | | | | | | | |
| TAQ polymerase | <1ml stock 20ul /reaction used | n/a | Liquid.<br>May be harmful by inhalation, skin contact and may cause eye irritation. | 3x2=**6** | Laboratory coats and nitrile gloves must be worn.<br><br>Such small volumes are used <2/1000th of a ml and the method of handling (Gilson pipette and Eppendorf) make contact with the eye and subsequent irritation extremely unlikely. | 1x2=**2** | Clinical waste | FIRE: Any extinguisher<br>SPILLAGE: Wipe up with tissue and dispose of as clinical waste. |
| DNA Ladder | 2ul | n/a | Liquid<br>Contains less than 1% of a substance regarded as hazardous | 2x2 =**4** | It is essential that nitrile gloves and labcoat are worn.<br>Such small volumes are used <2/1000th of a ml and the method of handling (Gilson pipette and Eppendorf) make contact with the substance extremely unlikely. | 1x1=**1** | Gels can be disposed of in the landfill waste. | FIRE: Any extinguisher<br>SPILLAGE: Wipe up with tissue, general waste. |
| Agarose | <2g in 100ml of buffer | Not available | Solid<br>Absorption/Ingestion<br>Respiratory and absorption through skin. | 4 x 3 =**12** | Avoid contact with the skin; use nitrile gloves and lab coat. | 1 x 3 = **4** | Dispose of solidified unstained product through landfill. | Wait until cool and shovel. |

| | | | | | | | | |
|---|---|---|---|---|---|---|---|---|
| | | | May be harmful if inhaled.<br>May cause respiratory tract irritation.<br>May be harmful if swallowed.<br>May be harmful if absorbed through skin.<br>May cause skin irritation.<br>May cause eye irritation. | | Use safety glasses and thermal gloves when handling hot liquid agarose.<br>Vessels used to heat agarose should under no circumstances be covered. | | | |
| Sodium Hydroxide<br>25mM | 8g in 1litre ddH20 when making SB Buffer | 2mg/m 3 | Liquid/solid<br>Ingestion/absorption<br>Skin irritation<br>Eye irritation<br>Specific target organ toxicity<br>May be harmful if inhaled or swallowed. | 3 x 4 = 12 | Wear safety glasses, nitrile gloves, lab coat. | 1 x 4 = 4 | Retain residue in waste bottle contained in fume cupboard. Disposal through chemistry stores. | Spillage: Sweep up and shovel. Keep in suitable, closed containers for disposal.<br>When liquid: use paper towels to contain spillage. Contaminated paper towels can be disposed of through landfill waste. |
| Boric Acid | For 1 litre 5Xsolution use 27.5 g Boric acid | Not available | Solid/LiquidPowder, dissolved in solution.<br>Absorption/Ingestion/Inhalation<br>Toxic when swallowed. **May damage fertility or the unborn child.** May be harmful if inhaled, May be harmful if absorbed though skin,<br>May cause eye irritation. | 3 x 4 = **12** | Wear nitrile gloves, lab coat and safety glasses.<br>Avoid contact with skin.<br>**If you are pregnant or think that you may be pregnant this risk assessment must be re-evaluated with input from the departmental safety officer.** | 1 x 4 = **4** | Dispose of residues down the drain with plenty of water. | Spillage: Pick up and arrange disposal without creating dust. Sweep up and shovel.<br>When liquid, contain spillage by soaking up with tissue that can be disposed of through landfill waste. |

| | | | | | | | | |
|---|---|---|---|---|---|---|---|---|
| Sucrose | 40% in 6x loading buffer. About 1 ul 6X loading buffer per sample | Not available | Solution. Non-hazardous | 1 x 1 = **1** | None | 1 x 1 = **1** | Dilute and wash down the sink | |
| Bromphenol blue | 0.25% in 6x loading buffer. About 1 ul 6x loading buffer per sample | Not available | Solid/Liquid Absorption/Inhalation<br>/Ingestion<br>absorption through skin<br>HARMFUL if inhaled, if swallowed and if absorbed through skin. | 3 x 4 = **12** | Wear nitrile gloves and lab coat.<br>Use in a well ventilated area to avoid inhalation. | 1 x 4 = **4** | Small amounts can be washed down the sink with copious amounts of water. | Spillage: Pick up and arrange disposal without creating dust. Sweep up and shovel. When liquid, contain spillage by soaking up with tissue that can be disposed of trough landfill waste. |
| Xylene cyanol | <5ul per sample | Not available | Liquid. Irritant.<br>Hazardous in case of ingestion.<br>Slightly hazardous in case of eye contact and<br>inhalation. | 2x2=4 | Wear nitrile gloves and lab coat.<br>Keep away from heat. Keep away from sources of ignition.<br>Immediately flush eyes with running water for at least 15 minutes, keeping eyelids open. Cold water may be<br>used.<br>No known effect on skin contact, rinse with water for a few minutes<br>Allow the victim to rest in a well ventilated area. Seek immediate medical attention. | 1x1=1 | Use appropriate tools to put the spilled solid in a convenient waste disposal container. Finish cleaning by spreading water on the contaminated surface and dispose of according to local and regional authority requirements. | Spillage: Pick up and arrange disposal without creating dust. Sweep up and shovel. When liquid, contain spillage by soaking up with tissue that can be disposed of trough landfill waste. |

| | | | | | | | | |
|---|---|---|---|---|---|---|---|---|
| GelRed | 1.5ul in 15ml electrophoresis buffer | Not available | Liquid<br>Non-hazardous | 1 x 1 = 1 | None | 1 x 1 = 1 | Dilute and wash down the sink | |
| **Qubit quantification** | | | | | | | | |
| Qubit ® dsDNA HS Reagent *200X concentrate in DMSO* | 1ul per sample | n/a | • Liquid<br>• Causes mild skin irritation<br>• Causes eye irritation<br>• May cause respiratory irritation | 3 x 2 = 6 | • Wear lab coat and nitrile gloves<br>• Avoid contact with skin | 1 x 3 = 3 | Dispose of small quantities by placing used tubes in clinical waste for incineration | Spillage: Absorb spill with vermiculite or other inert material, then place in container for chemical waste |

Add more lines if required

| | | | | |
|---|---|---|---|---|
| UNIVERSITY OF Hull<br>Department of Biological and Marine Sciences | PROCEDURAL RISK ASSESSMENT AND COSHH FORM | | | |
| | ref: BMS101146v1 | Date: 26/10/22 | Group: Teaching | Overall Risk Level: Low |

| BIOLOGICAL SUBSTANCES | | | | ☐ **NO** Biological Substances Used |
|---|---|---|---|---|
| **Biological Agents or Hazards** | **Human**<br>☐ Cells<br>☐ Tissue/body parts<br>☐ Primary cell culture<br>☐ Continuous cell culture<br>☐ Blood<br>☐ Human excretions/fluids<br>☐ Patient contact | **Animal**<br>☐ Cells<br>☐ Tissue/body parts<br>☐ Primary cell culture<br>☐ Continuous cell culture<br>☐ Blood<br>☐ Animal excretions/fluids<br>☐ Animal contact | **Microorganism**<br>Pathogens listed by ACDP/DEFRA as<br>☒ Hazard Group 1<br>☐ Hazard Group 2<br>☐ Hazard Group 3<br>☐ Not Listed by ACDP/DEFRA<br>☒ Unknown | **Other**<br>☐ Plants<br>☐ Soil<br>☐ Toxins<br>☐ Carcinogens<br>☐ Allergen<br>☒ DNA<br>Other: |
| | ☐ Genetic Modified Organism<br>☐ Lentiviral Transfection | | ☐ Genetic Modified Microorganism | ☐ Unknown Biological Hazards<br>☐ No **known** Biological Hazards |
| **Description, include species, cell line or technical name** | E. coli K12 (provided by teaching lab)<br>**For work to be undertaken by Immunocompromised people/pregnant women: an individual risk assessment must be completed to evaluate the risk on a case-by-case situation.** | | | |
| **Source** | ☐ Unknown<br>☐ Isolated in own Laboratory | ☒ Provided by: Teaching lab<br>☐ Purchased from: High Street Retailer<br>☐ Collection Site: | | |
| **Quantity/Scale** | x Small ☐ Medium ☐ Large | **Risk if maximum quantity used** | X Negligible ☐ Low ☐ Medium ☐ High | |
| **Exposure Route** | ☐ Inhalation (Airborn) ☐ Ingestion ☒ Splash in eyes or mouth ☐ Percutaneous (Skin) ☐ Animal bite or scratch<br>☐ Other: | | | |
| **Exposure frequency** | ☐ Daily ☐ Weekly ☐ Monthly ☒ Other: Once | **Potential of exposure** | x Negligible ☐ Low ☐ Medium ☐ High | |

| | |
|---|---|
| **Risk of Aerosol or Airborne Particle formation** | x Negligible ☐ Low ☐ Medium ☐ High |

| | |
|---|---|
| **Risk to Human Health** | x Effectively Zero ☐ Low ☐ Medium ☐ High<br>Disease or Condition that might be caused upon exposure (specify): |
| **Health Surveillance** | ☐ Not required ☒ Only required upon exposure to risk (incident/accident) ☐ Required |
| **Effective Treatment** | ☐ Do not know if available ☐ Not available ☒ Available (specify): Antibiotics |
| **Immunisation and Prophylaxis** | ☐ Not available ☒ Not required ☐ Optional ☐ Essential<br>☐ Other: N/A |
| **Risk to Animal Health** | x Effectively Zero ☐ Low ☐ Medium ☐ High<br>Disease or Condition that might be caused upon exposure (describe if applicable):<br><br>Effective treatment available: ☒ No ☐ Yes (specify): |
| **Risk to the Environment** | x Effectively Zero ☐ Low ☐ Medium/Low ☐ Medium ☐ High<br>Possible hazards/risks/damage to the Environment (describe if applicable):<br><br>Effective remediation available?<br>☐ No ☐ Yes (specify): N/A |
| **Facility** | x Laboratory ☐ Animal Facility ☐ Plant Facility ☐ Microbiological Containment Facility ☐ Other: |
| **Containment Level** | x No containment required ☐ Containment Level 1 (CL1) ☐ Containment Level 2 (CL2) ☐ Containment Level 3 (CL3) |
| **Biosafety Cabinet** | x No biosafety cabinet required ☐ Class 1 ☐ Class 2 ☐ Class 3 |

| PROCEDURAL RISK ASSESSMENT AND COSHH FORM | | | Page 17 |
|---|---|---|---|
| ref: BMS101146v1 | Date: 26/10/22 | Group: Teaching | Overall Risk Level: Low |

| | | |
|---|---|---|
| **Other Controls** | ☐ Spill tray ☐ Physical barrier to contain splashing ☒ Tubes secured with cap to contain aerosols ☒ Secondary containment<br>☐ Sharps not permitted ☐ Other (specify): | |
| **Long-Term Storage (>24 hours)** | x No storage required ☐ Locked Storage ☐ Cupboard ☒ Refrigerator ☐ -20°C Freezer ☐ -80°C Freezer ☐ Liquid Nitrogen<br>☐ Incubator ☐ Other: | |
| **Transport** | ☐ Allowed, if appropriate containment is used to avoid accidental release during transport<br>Specify containment:<br>☐ Transport according to Dangerous Goods Classification:<br>☐ Transport only allowed with HSE consent/Environmental Agency/DEFRA license<br>License Number or Consent Letter (specify):<br>x Other: n/a | |
| **Inactivation** | ☒ Chemical inactivation ☒ Autoclaving (heat inactivation) ☐ Incineration (heat inactivation) ☐ Fumigation | |
| **Disinfection** | **Surfaces** | ☐ 70% Ethanol ☒ 1% (w/v) Virkon solution ☐ 10% (v/v) Trigene/Distel solution ☐ Clinell sanitising wipes<br>x Other: soap and water |
| | **Hand wash** | ☐ Lab Guard microbial soap ☒ Other: lab hand soap |
| | **Spills** | ☐ Virkon powder ☒ Other: water |
| | **Water bath** | ☐ Aquaresist ☐ Other: N/A |
| **Waste Disposal** | ☐ Clinical waste ☒ Autoclaved waste ☐ Landfill waste Sink Disposal after effective inactivation<br>☐ Other: | |
| **Instruction, Training and Supervision** | x Appropriate Instruction required regarding Biological Safety, including appropriate Standard Operating Procedures (SOP)<br>☒ Basic training required ☐ Specialist training required ☐ Constant supervision required<br>Add details: | |

| | |
|---|---|
| **Consent or License from DEFRA, EA, DEFRA, HSE or Home Office** | x Not required ☐ Required/obligatory, provide HSE consent/DEFRA license number and details: |
| **Emergency Procedure(s)** | x No special requirements for the used biological substances<br>☐ Required (describe): |

| SAFE WORKING PROCEDURE |
|---|
| • Wear suitable protective clothing, laboratory coat, gloves and eye protection where required<br>• Strictly follow procedure protocol as described in Protocol/SOP Ref: |
| • Work to Good Laboratory Practice.<br>• Wear suitable protective clothing, laboratory coat, nitrile gloves and eye protection where required.<br>• Only carry out microbiological work in the designated areas.<br>• Separate risk assessment required for immunocompromised/ pregnant people handling microbes directly. |

| FIRST AID |
|---|
| **Basic First Aid** |
| If skin contact occurs, wash off in running cold water for 10 minutes<br>If chemical ingested, rinse out mouth with cold water and spit out<br>If eye contact occurs, irrigate with cold water for 10 minutes<br>Contact a University first aider |
| **Specific First Aid (if different from above as a result of the risk assessment & COSHH requirements)** |

### EMERGENCY CONTACTS

Please use emergency contacts in the order shown below.
If you are unable to reach the first person on the list, contact the next person, etcetera.

| Role | Name | Telephone Numbers |
|---|---|---|
| Supervisor | **Dr Katharine Hubbard** | 2347 |
| Departmental/Local Safety Officer | Rob Donnelly (BMS)<br>Sonia Jennings (FoSE)<br>Rose Terschak (GEE) | 6213<br>5518<br>2344 |
| University Biological Hazards Officer | Frank Voncken | 5280 |
| Health and Safety Services Team | Secretary<br>Alan Hewett<br>Tim Coldwell<br>Rob McDonald | 5165<br>6571<br>6992<br>2148 |
| University Hull Campus | Emergency Call Centre | 5555 |

| PROCEDURAL RISK ASSESSMENT AND COSHH FORM | | | Page 22 |
|---|---|---|---|
| ref: BMS101146v1 | Date: 26/10/22 | Group: Teaching | Overall Risk Level: Low |

### USER AGREEMENT & AUTHORISATION

User signatures - indicate that you agree to abide by the recommendations indicated above, and have read and understood the associated standard operating procedures where these are referred to.

| **Name** (Block capitals) | **Signature** | **Date** | **Supervisor Name** | **Signature** | **Date** |
|---|---|---|---|---|---|
